# Non-canonical Subthalamic Nucleus projections regulate motor variability

**DOI:** 10.64898/2026.11.12.765751

**Authors:** Oluwatomiwa A. Oladunni, Sophie K. Elvig, Hunter C. Bennis, Laila A. Elzaree, Siddhartha Nandy, Yuting Liu, Snigdhansu Chatterjee, Brian N. Mathur, Steffen B.E. Wolff

**Affiliations:** Department of Pharmacology and Physiology, University of Maryland School of Medicine, Baltimore, MD, 21201, USA; Department of Mathematics and Statistics, University of Maryland Baltimore County, Baltimore, MD, 21250, USA; University of Maryland Medicine Institute for Neuroscience Discovery, University of Maryland Baltimore, Baltimore, MD, 21201, USA

## Abstract

Regulation of motor variability is a key hallmark of motor skill learning, balancing motor exploration with optimization of identified task solutions. Nevertheless, the underlying circuit mechanisms in the mammalian brain remain largely unknown. Subnuclei of the dorsal striatum in the basal ganglia promote stereotyped or variable behavior, respectively, and deep brain stimulation of the downstream subthalamic nucleus (STN) can alter motor variability. We therefore tested whether the STN regulates motor variability via non-canonical projections to striatum. We discovered that silencing STN neurons projecting to dorsolateral striatum impaired motor variability reduction in rats developing complex stereotyped movement patterns. In contrast, silencing STN neurons projecting to the anterior thalamic nuclei, or neurons in these nuclei that receive STN input and project to dorsomedial striatum, accelerated variability reduction. These results deepen our anatomical and functional understanding of subcortical motor circuits and reveal unknown roles for STN’s non-canonical striatal projections in motor variability regulation.

## MAIN

From tying shoelaces to serving a volleyball, our brain’s ability to learn and generate stereotyped, task-specific movement patterns is a crucial part of our daily lives. A key hallmark of developing such motor skills *de novo* is a gradual reduction of trial-to-trial motor variability. Over extended periods of training, initially highly variable movements develop into skilled movement patterns with precise and stereotyped kinematics across trials. While motor variability may be partially due to an inherently noisy nervous system and detrimental to peak performance^1–5^, findings from reinforcement and motor learning show that variability is indeed essential for learning and adaptation. Importantly, variability can even be actively regulated to balance the exploration of the motor space to identify appropriate task solutions and the optimization of successful movement patterns^2–12^. However, the neural circuits and mechanisms underlying such an active regulation of motor variability are poorly understood.

The best-studied example of motor variability regulation comes from songbirds, where the lateral magnocellular nucleus of the anterior nidopallium (LMAN) drives vocal exploration by injecting variability into the song production circuitry^6,13–24^. While no homologous mammalian structure has been identified, LMAN is the output nucleus of a pathway akin to mammalian cortico-basal ganglia-thalamocortical loops^17,25^. In these pathways, the basal ganglia (BG) are particularly promising candidates for motor variability regulation. The BG are critical for behaviors that require regulation of behavioral variability, from action selection^26–28^ to decision-making^29–32^, and, importantly, learning and execution of motor skills^33,34^. Accordingly, many disorders affecting the BG disturb the balance between variable and stereotyped behavior^35–40^, including disruptions of motor variability regulation in Parkinson’s Disease (PD)^12,41–46^. Notably, Deep Brain Stimulation in the subthalamic nucleus (STN) in the BG can bidirectionally affect behavioral and motor variability in PD and other disorders^46–53^. However, whether and how the STN also contributes to the physiological regulation of motor variability is unclear.

While the STN enacts its canonical function in suppressing behaviors^54–57^ via the indirect BG output pathway, it also targets additional structures, including thalamic and brainstem nuclei^58–60^ as well as the, canonically upstream, striatum. The STN does not only directly project to the dorsolateral striatum (DLS)^61–63^, but may also indirectly project to the dorsomedial striatum (DMS): The STN projects to the anterior thalamic nuclei (ATN)^64^, which, in turn, project to the DMS^65–67^. Notably, these STN pathways to the DLS and DMS are poorly understood functionally, and completely untested in the context of motor variability.

In line with their respective general functions in flexible^34,68–70^ and automatic^33,34,68,69^ behavior, a transition from the DMS to the DLS parallels the decrease in motor variability across motor skill learning: the DMS contributes to exploratory behavior early in learning, but becomes dispensable in experts, while the DLS is necessary for the generation of learned, stereotyped movement patterns^33,34,71,72^. Accordingly, the DLS features fine-scaled representations of the detailed learned kinematics^33,71,73,74^ and spiking variability in the DLS is directly correlated with the variability of the generated learned movement patterns^75^. In light of this, we posit that the STN’s non-canonical direct and indirect projections to the striatum may modulate DLS and DMS activity over the course of motor learning to suppress or promote motor variability, respectively.

To directly test this, we trained rats in a task resulting in the development of complex, stereotyped movement patterns and selectively silenced distinct projection neurons along the suggested projection pathways. We found that STN neurons projecting to the DLS promote stereotyped behavior. In contrast, STN neurons projecting to the ATN, ATN neurons that project to the DMS as well as ATN neurons that both receive STN input and target the DMS favor variable behavior. Taken together, our results identify novel circuit mechanisms for the active regulation of motor variability in the mammalian brain and provide new insights into the connectivity and function of the STN, the BG, and the distributed subcortical motor network.

## RESULTS

### Distinct STN neuron populations project to the DLS or to the ATN

While non-canonical projections from the STN to the DLS and to the ATN had previously been individually described in multiple species^61–64^, we first aimed to confirm their existence in the rat and to determine whether they originate from overlapping or distinct neuronal populations. We therefore performed retrograde viral tracing from either the DLS or the ATN in separate animals by injecting retrogradely infecting adeno-associated viruses for the expression of GFP (rgAAV-GFP; **Fig. 1A, B**). Both injections revealed fluorescently labeled neurons in the STN, confirming the existence of DLS-projecting^61–63^ (abbreviated as STN^DLS^ neurons; **Fig. 1A**) as well as of ATN-projecting STN neurons^64^ (STN^ATN^ neurons; **Fig. 1B**) in the rat. Using a similar tracing approach (rgAAV-mCherry in the DMS), we also confirmed that DMS-projecting ATN neurons^65–67^ (ATN^DMS^ neurons) exist in the rat (**Fig. 1C**).

**Figure 1.**
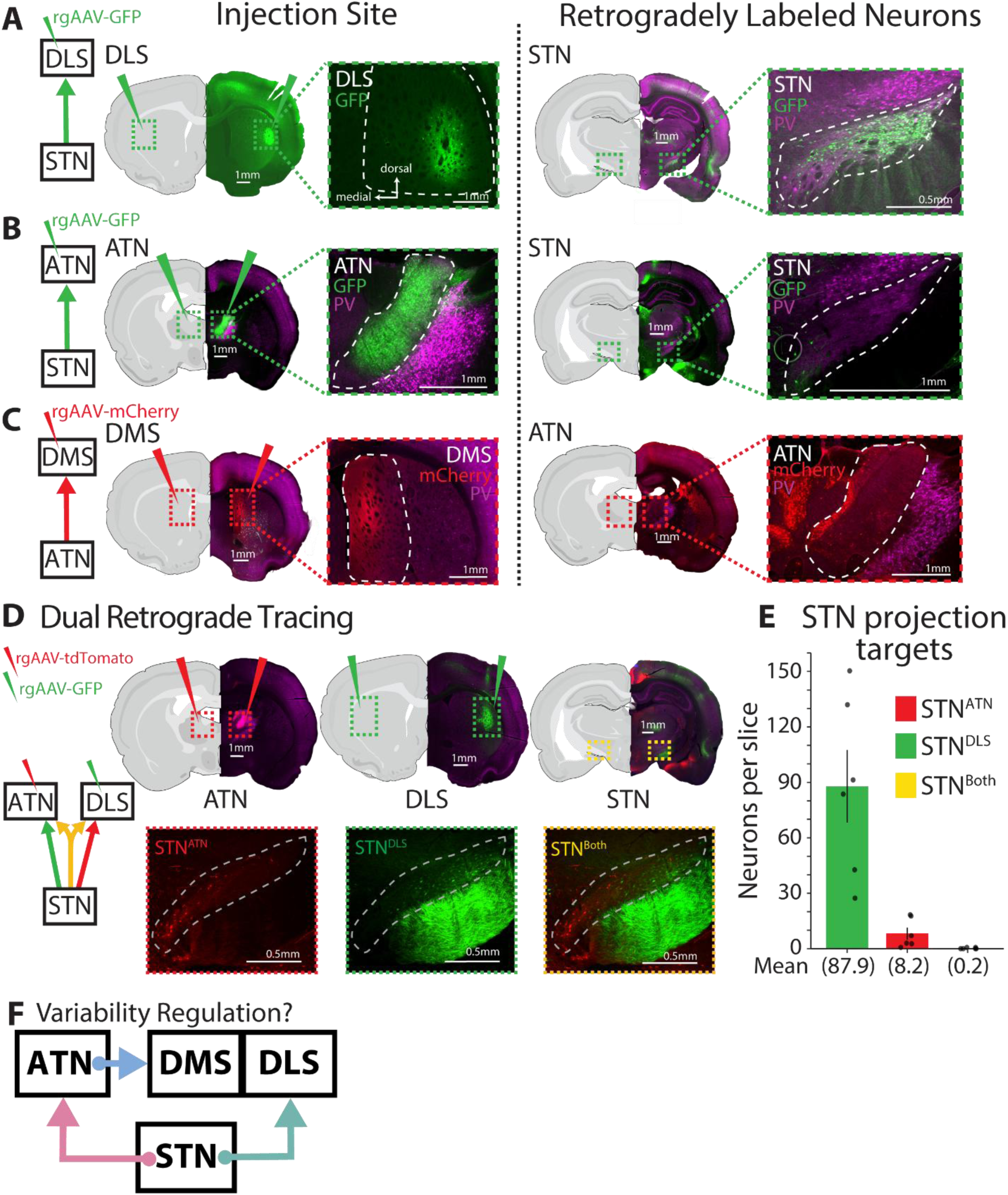
Distinct STN neurons project to the DLS or the ATN, and ATN neurons target the DMS. **(A)** Retrograde viral tracing to label DLS-projecting STN neurons (STN^DLS^). Left: Representative viral injection site in the DLS (outlined in white). Right: Representative image of retrogradely labeled STN neurons (green). **(B)** As (A), but for labeling of STN^ATN^ neurons. **(C)** As (A), but for labeling of ATN^DMS^ neurons. **(D)** Dual retrograde viral tracing of STN^DLS^ and STN^ATN^ neurons. Top: Representative images of viral injection sites in the DLS and ATN and of retrograde labeling of STN neurons. Bottom: Representative images of retrogradely labeled STN neurons. **(E)** Quantification of identified STN^DLS^, STN^ATN^, and STN^Both^ (STN neurons that project to both DLS and ATN) average number of neurons per analyzed brain slice (n = 6 hemispheres; 3 animals, counts averaged across 3 slices per hemisphere). **(F)** Simplified circuit scheme for the hypothesized bidirectional regulation of motor variability by projection pathways originating in the Subthalamic Nucleus. STN: Subthalamic Nucleus, ATN: Anterior Thalamic Nuclei, DMS: Dorsomedial Striatum, DLS: Dorsolateral Striatum.

To determine whether STN^DLS^ and STN^ATN^ neurons are distinct populations, we performed dual retrograde viral tracing (**Fig. 1D, E**): We injected retrograde AAVs for the expression of distinct fluorophores (rgAAV-GFP and rgAAV-tdTomato) into the DLS and the ATN, respectively, allowing us to identify STN neurons that project to either or both target areas (**Fig. 1D**). STN^DLS^ and STN^ATN^ neurons showed not only a difference in abundance, with a majority of labeled neurons projecting to the DLS (**Fig. 1E**), but also distinct anatomical distributions: Across the anterior-posterior axis, STN^DLS^ neurons were predominantly localized in lateral parts of the STN, corresponding to the STN’s motor-related area, while STN^ATN^ neurons were mainly in the medial STN (**Fig. 1D**), corresponding to the STN’s limbic and/or associative segments. Importantly, in line with the different localization patterns, we observed minimal overlap between labeled neurons (**Fig. 1D, E**), suggesting that STN^DLS^ and STN^ATN^ neurons constitute anatomically largely distinct populations. Together, these tracing experiments confirmed the existence of the suggested projection pathways (**Fig. 1F**), allowing us to functionally study them in the context of motor variability regulation in our subsequent experiments.

### STN projection neurons bidirectionally contribute to motor variability regulation

To test whether these distinct STN projection neuron populations contribute to *de novo* motor skill learning in general, and to the regulation of motor variability in particular, we took advantage of a behavioral paradigm we previously used to probe the role of the basal ganglia in motor learning^33,34,76^. Rats are trained to press a lever twice with a specific time interval (inter-press interval (IPI), target 700 ms; **Fig. 2A**) to gain water rewards, scaled by the distance to the target.

**Figure 2.**
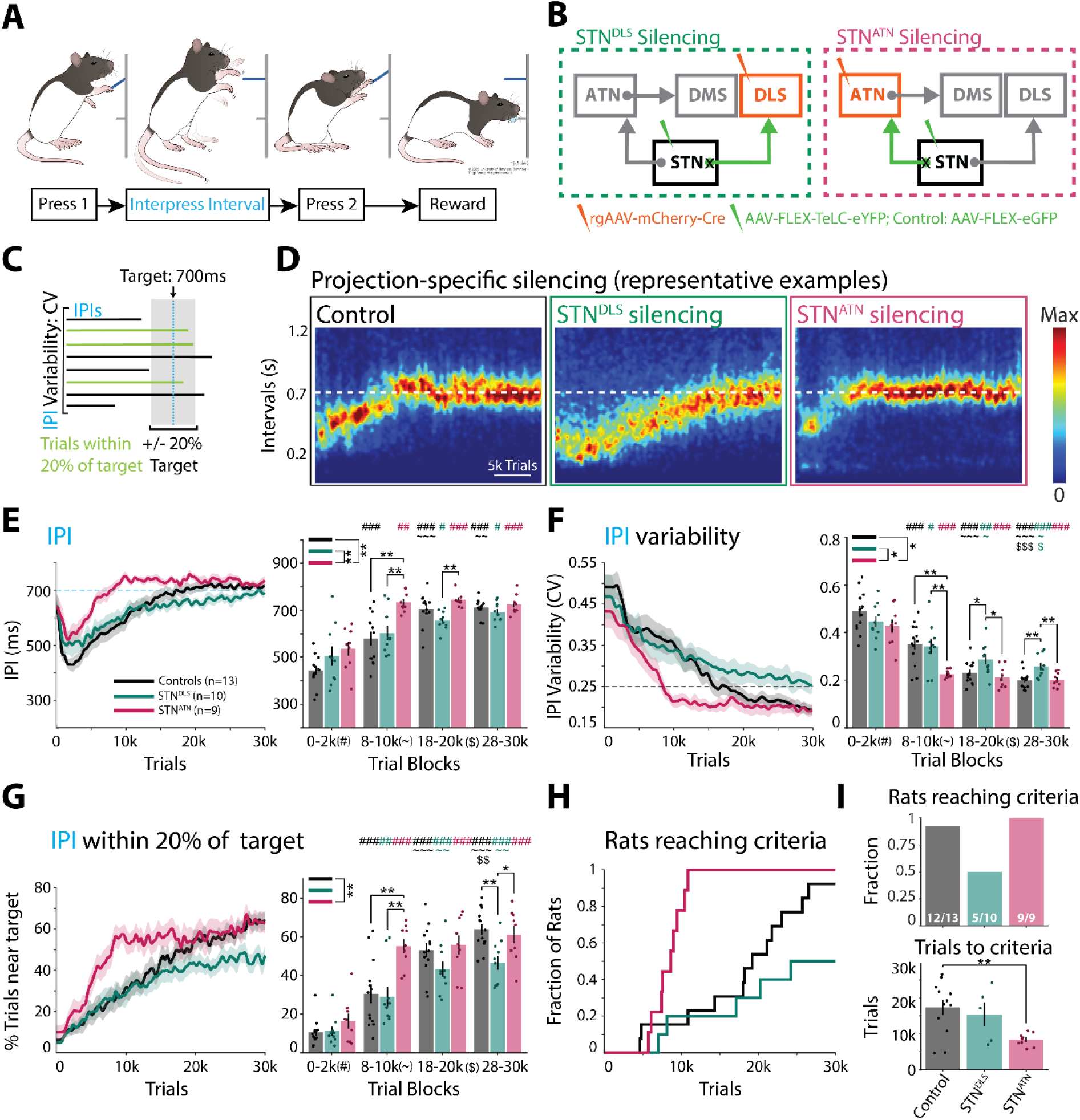
STN^DLS^ and STN^ATN^ neurons bidirectionally contribute to the regulation of motor variability during motor skill learning. **(A)** Motor skill learning paradigm. Rats are rewarded with water for pressing a lever twice with an inter-press interval (IPI) close to a target (700 ms). Rats develop idiosyncratic, complex, and stereotyped movement patterns as task solutions to produce rewarded IPIs. **(B)** Viral approach to chronically silence projection-defined STN^DLS^ and STN^ATN^ neurons. See text for details and **Extended Data** Fig. 1 for injection sites. TeLC: Tetanus Toxin Light Chain, YFP: Yellow Fluorescent Protein, GFP: Green Fluorescent Protein. Animals in the control group express GFP in STN^DLS^, STN^ATN^, or ATN^DMS^ neurons (see Methods and **Extended Data** Fig. 2 for comparison of control cohorts). **(C)** Scheme depicting the quantified performance measures: mean IPI, IPI variability measured as coefficient of variation (CV), and fraction of trials with IPIs near the target (i.e., trials within +/-20% of the target). **(D)** Effects of pre-training manipulations on task performance throughout learning for Controls, STN^DLS^ neuron silencing, and STN^ATN^ neuron silencing. Shown are heatmaps of IPI probability distributions for representative rats. **(E)** Left: Mean IPI across training. Right: Quantification of mean IPI in select trial blocks. Dotted line indicates target IPI. **(F)** As (E), but for CV of the IPIs. Dotted line indicates CV threshold for learning criteria (see text). **(G)** As (E), but for fraction of trials close to the target. **(H)** Fraction of rats reaching learning criteria (see text) across learning. **(I)** Top: Fraction of rats that reached the learning criteria. Bottom: Average number of trials needed to reach the learning criteria. Statistics: (E-G) 2-way repeated measures ANOVA with post-hoc tests (Holm). (H) 1-way ANOVA with post-hoc tests (Holm). For all statistical details, see **Supplementary Table 1**. Bars show means, dots show individual animals, and error bars show SEM. Insets with group colors and stars indicate significance for overall group comparisons. Stars above bars indicate significant differences across groups within a block. Symbols indicate significant differences across blocks for a given group. The symbol indicates the compared block (i.e., # compared to 0-2k block, ∼ compared to 8-10k block, $ compared to 18-20k block), the color indicates group (e.g., black for Controls). *p<0.05, **p<0.01, and ***p<0.001.

Over weeks of operant shaping, rats develop *de novo* idiosyncratic movement patterns, including the lever presses and additional movement elements, that allow them to produce the target IPI^33,34,76^. The developing movement patterns become increasingly complex, spatiotemporally precise and stereotyped, allowing the animals to generate increasingly precise IPIs and more reliably gain rewards^33,34,76^. This task therefore allows us to study the development of variability on the level of the generated IPIs and of the emerging kinematics over weeks of training and tens of thousands of trials^33,34,76^.

To directly probe the contributions of the STN^DLS^ and STN^ATN^ projection neuron populations in this task, we used an intersectional viral strategy to chronically silence synaptic transmission in naïve (untrained) animals: We injected a retrograde AAV for the expression of Cre recombinase (rgAAV-Cre) either into the DLS or into the ATN, and a local AAV for Cre-dependent expression of Tetanus Toxin Light Chain (TeLC) into the STN (**Fig. 2B** *and **Extended Data Fig. 1***). TeLC chronically silences synaptic transmission by cleaving the synaptic protein VAMP2, preventing membrane fusion of synaptic vesicles and neurotransmitter release^77–79^. Our targeting approach restricts TeLC expression to STN neurons projecting to the DLS or the ATN, but we note that these neurons may have axon collaterals targeting other brain areas (see below and Discussion). After recovery from surgery, rats started training in our automated behavioral training system^80^ (see Methods) on our motor skill learning paradigm and completed at least 30,000 trials across ∼4-6 weeks of daily training (2 training sessions per night). Learning and performance of the experimental animals were compared to a cohort of rats that expressed GFP instead of TeLC. Since cohorts of animals with GFP expression in distinct projection neurons did not show any differences in learning (***Extended Data Fig. 2***), we combined them in one control group.

Over weeks of training, control animals learned to produce gradually longer IPIs, eventually reaching a mean IPI length close to the 700 ms target (**Fig. 2C-E**). The generated intervals also became increasingly stereotyped, reflected by a continuous decrease in the coefficient of variation (CV) of the IPI distributions (**Fig. 2C, F**). Together, these changes led to a gradual increase in the fraction of trials close to the target (IPI +/-20% around target) (**Fig. 2C, G**), and allowed almost all control animals to reach our prespecified performance criteria for mastering the task (mean IPI +/-10% of target and CV of IPIs of less than 0.25)^76^ in less than 30,000 trials (average: 17,335 +/-2,123 trials; **Fig. 2H, I**).

Chronic silencing of STN^DLS^ or STN^ATN^ projection neurons did not affect task engagement, or the ability to generate trials with the prescribed two lever presses to receive rewards (***Extended Data Fig. 3A-E***).

However, their silencing dramatically affected performance during motor learning. First, while the average IPI duration was indistinguishable from controls at the beginning and end of training in both experimental groups, STN^ATN^ neuron silencing led to a significantly faster IPI increase in early phases of learning (**Fig. 2E**). Even more pronounced, however, were bidirectional effects on the development of the IPI variability. While initial IPI variability was unaffected in both groups, STN^ATN^ neuron silencing significantly accelerated the decrease in variability early in training, allowing animals to reach the same final variability levels as controls ∼10,000 trials earlier (**Fig. 2F**). In stark contrast, STN^DLS^ neuron silencing impaired the progressive IPI variability decrease, resulting in significantly higher variability levels than observed in controls at later stages of training (**Fig. 2F**). Notably, our manipulations also affected the variability of a task aspect that does not encourage stereotypy: After unrewarded trials, rats need to observe a minimum timeout period (inter-trial interval (ITI), threshold 1.2 s; ***Extended Data Fig. 3H***) before initiating a new trial^76^. Although only the minimum ITI duration, but not ITI precision, is reinforced, control animals reduced ITI variability over training (***Extended Data Fig. 3H, I***). As for IPIs, STN^ATN^ neuron silencing significantly accelerated the ITI variability decrease early in learning, but without affecting average ITI duration (***Extended Data Fig. 3H, I***). STN^DLS^ neuron silencing, on the other hand, caused a noticeable, although not significant, trend towards higher ITI variability levels at the end of training (***Extended Data Fig. 3I***).

Our manipulations also affected measures of overall learning. STN^ATN^ neuron silencing significantly accelerated the increase of the fraction of trials close to the target early in training, while STN^DLS^ neuron silencing impaired its gradual increase, resulting in significantly lower levels at the end of training than in controls (**Fig. 2G**). Finally, after STN^ATN^ neuron silencing, rats reached our learning criteria on average after only 8,338 +/-599 trials, significantly faster than controls (**Fig. 2H, I**). In contrast, after STN^DLS^ neuron silencing, half of the rats never reached the learning criteria in our training period of 30,000 trials (**Fig. 2H, I**). Together, our results show that distinct STN projection neurons bidirectionally contribute to the development of variability over the course of *de novo* motor skill learning, with STN^DLS^ neurons promoting stereotyped and STN^ATN^ neurons promoting variable motor behavior.

### STN^DLS^ and STN^ATN^ neurons are anatomically and functionally distinct from other STN projection neurons

The STN projects to various subcortical areas besides the DLS and ATN^58–60^, but whether these projections originate from overlapping or distinct populations of projection neurons is unclear. This raises the possibility that STN^DLS^ and STN^ATN^ neurons might send potential axon collaterals to other brain areas – which would also be affected by our silencing approach and whose loss might contribute to the observed behavioral effects. Furthermore, since we cannot fully exclude potential spillover of viral injections to thalamic nuclei adjacent to the ATN (cf. ***Extended Data Fig. 1F***), we might have inadvertently silenced additional, previously unidentified STN projection neuron populations targeting these nuclei. We therefore aimed to increase the specificity of our results with additional behavioral and anatomical experiments.

In addition to the suggested indirect pathway from the STN via the ATN to the DMS (**Fig. 1F**), the STN may also directly project to the DMS^61–63^, making it a major potential target for axon collaterals of STN^DLS^ or STN^ATN^ neurons. We therefore performed dual retrograde viral tracing to determine whether this projection exists in the rat and to test for potential overlap. This revealed a medial population of DMS-projecting STN neurons which were distinct both from STN^DLS^ and STN^ATN^ neurons (***Extended Data Fig. 4***). Next, we silenced the DMS-projecting STN neurons in an additional cohort of rats and trained them in our motor task. Notably, silencing these neurons had no effect on variability development or other aspects of motor learning (***Extended Data Figs. 5*, 6A-C**). Together, this demonstrates that STN^DLS^ and STN^ATN^ neurons are anatomically and functionally distinct from DMS-projecting STN neurons, and that the behavioral effects of their silencing cannot be explained by inadvertent silencing of direct STN inputs to the DMS.

Potential spillover from our viral injections into the ATN (***Extended Data Fig. 1F***) might have reached the adjacent rostral intralaminar thalamic nuclei (rILN) or the reticular thalamic nucleus (RTN). Furthermore, STN^ATN^ (or STN^DLS^) neurons could potentially send axon collaterals to these thalamic nuclei. However, to our knowledge, direct STN projections to these nuclei have not been described. Nevertheless, we aimed to test whether inadvertent silencing of such potential projections might have contributed to the effects of STN^ATN^ (or STN^DLS^) neuron silencing (cf. **Fig. 2**). To silence potential STN projection neurons targeting the rILN or the RTN with TeLC, we targeted injections of rgAAV-Cre in separate cohorts of rats to the parts of the rILN and RTN that are immediately adjacent to the ATN and would have been the most likely to be affected by potential spillover (***Extended Data Fig. 6D-I*** *and cf. **Extended Data Fig. 1F***). Importantly, when we trained these animals on our motor task, they did not show any changes in motor variability regulation or other aspects of motor learning (***Extended Data Fig. 7***). These results support the specificity of the behavioral effects of our STN^ATN^ neuron silencing experiments, showing that they are not due to inadvertent silencing of STN neurons projecting to the rILN or the RTN.

Taken together, these results show that the STN^DLS^ and STN^ATN^ neuron populations are anatomically and functionally distinct from other STN populations projecting to the striatum and thalamus and play a specific role in the regulation of motor variability during *de novo* skill learning.

### Variability regulation by STN^DLS^ and STN^ATN^ neurons is trial outcome-dependent

During motor adaptation, motor variability is rapidly modulated by trial outcome, leading to higher variability after unrewarded than after rewarded trials^3,8^. This favors the repetition of successful kinematic solutions and further exploration after unsuccessful attempts. Notably, the STN receives dopaminergic input from the Substantia Nigra pars compacta and the Ventral Tegmental Area^81–83^ and has been implicated in reward and value processing^84–87^. Together, this suggests that STN^DLS^ and STN^ATN^ neurons may contribute to the regulation of motor variability in a trial outcome-dependent manner. To test this, we calculated trial-to-trial IPI changes after rewarded and after unrewarded trials separately (**Fig. 3A**). To account for varying reward contexts, which can significantly affect the baseline levels of motor variability^3^, we used a propensity score matching approach (see Methods), allowing us to isolate the effects of individual rewarded or unrewarded trials on the subsequent IPI variability^3,88^. To further determine whether the contributions of the STN^DLS^ and STN^ATN^ neurons depend on the overall reward context, i.e., whether an individual reward occurs in a stretch of high or low reward rates^3^ (**Fig. 3A**), we split our data into high and low reward context trials. For both contexts, we calculated the IPI difference between each rewarded or unrewarded trial (the trigger trial) and the subsequent trial (ΔIPI; **Fig. 3A**). This showed a decrease over learning for both rewarded and unrewarded trigger trials in controls (**Fig. 3A, B** *and **Extended Data Fig. 8B***). Consistent with our findings above, our manipulations bidirectionally affected the ΔIPI after rewarded trials: STN^ATN^ neuron silencing led to an accelerated decrease of the ΔIPI, while STN^DLS^ neuron silencing resulted in a higher ΔIPI in later training stages (**Fig. 3B***, left*). In contrast, after unrewarded trials, the ΔIPI was not significantly changed by either manipulation (**Fig. 3B***, right*). While slightly more pronounced in the high reward context, these effects were consistent across high and low reward contexts and when all reward contexts were combined (**Fig. 3B** *and **Extended Data Fig. 8A, B***). Together, this suggests that STN^DLS^ and STN^ATN^ neurons modulate motor variability in response to reward.

**Figure 3.**
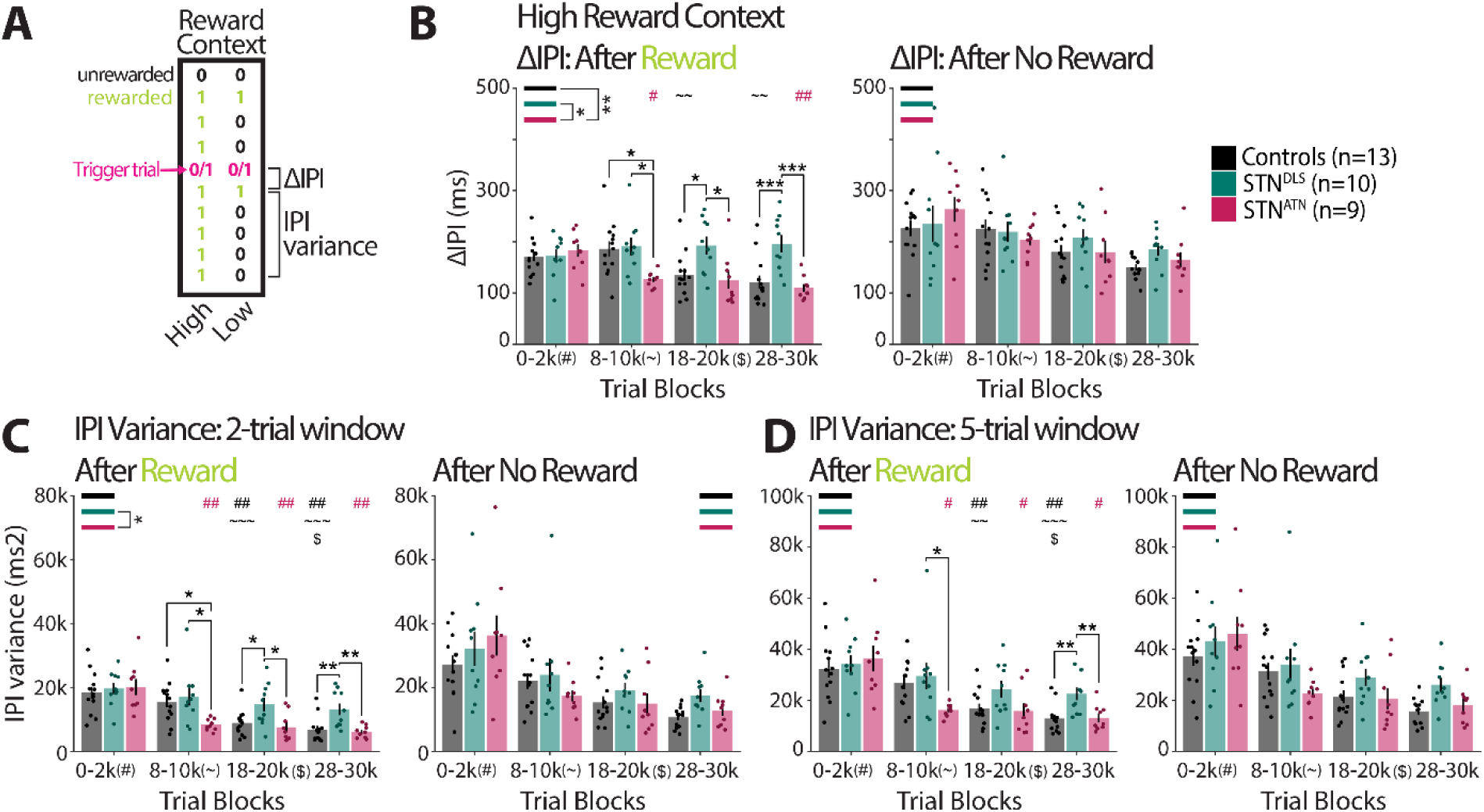
Contributions of STN^DLS^ and STN^ATN^ neurons to variability regulation depend on trial outcome. **(A)** Scheme depicting analysis of trial outcome-dependent variability regulation. For each rewarded or unrewarded trial throughout training (trigger trial, reward 0 or 1), the absolute difference between its IPI and the IPI of the subsequent trial (ΔIPI) as well as the variance of the IPIs across multiple trial windows (2-5 trials) after the trigger trial were quantified. Propensity score matching was used to account for varying reward contexts across individual trials (see Methods) and to divide trials into high or low reward contexts. **(B)** Mean ΔIPI for control and STN^DLS^ and STN^ATN^ neuron silencing groups across select trial blocks after rewarded (left) or unrewarded (right) trials in a high reward context (see **Extended Data** Fig. 8 for all and low reward contexts). **(C)** Like (B), but for the variance of the IPI in a 2-trial window after the trigger trial across all reward contexts. **(D)** Like (C), but for a 5-trial variance window. Statistics: 2-way RM ANOVA with post-hoc tests (Holm). For statistical details, see **Supplementary Table 2.** Bars show means, dots show individual animals, and error bars show SEM. Stars and symbols indicate significance as in Fig. 2.

To test whether STN^DLS^ and STN^ATN^ neurons affect variability also across longer trial periods, we calculated the IPI variance for different windows of trials following the trigger trial (2-5 trials; combined across all reward contexts; **Fig. 3A, C, D** *and **Extended Data Fig. 8C, D***). While we found the same effects of our manipulations on the variance in a 2-trial window as for the ΔIPI after rewarded trials (**Fig. 3C**), the effects started to dissipate for longer trial windows: STN^ATN^ neuron silencing did not significantly affect the variance, and after STN^DLS^ neuron silencing, increased variance levels only became apparent at the very end of training (**Fig. 3D** *and **Extended Data Fig. 8C, D***). This suggests that trial outcome affects the modulation of motor variability by the STN projection neuron populations over short stretches of trials.

Our analyses show that the modulation of motor variability by STN^DLS^ and STN^ATN^ neurons depends on the trial outcome. Both STN neuron populations affect variability after rewarded, but not unrewarded trials, suggesting that they specifically modulate how behavior is adapted in response to successful behaviors.

### STN^DLS^ and STN^ATN^ neurons bidirectionally modulate kinematic variability

So far we have focused on the variability of the generated intervals (**Figs. 2 *, 3***), but our task also allows us to go beyond this proxy for the developing movement patterns and to determine whether STN^DLS^ and STN^ATN^ neurons directly modulate the variability of the movement kinematics themselves^33,34,72^. Using machine learning-aided pose estimation (SLEAP^85^), we tracked the movements of individual body parts throughout training. As we previously described^33,34,72^, the animals’ initially highly variable movement patterns gradually develop into idiosyncratic kinematic solutions that become increasingly precise and stereotyped over the course of training (**Fig. 4A**). To quantify this development, we time-warped all trajectories to the same duration, allowing us to determine kinematic similarity independent of trajectory length, and calculated trial-by-trial correlations. STN projection neuron silencing did not affect general movement aspects like average velocity or acceleration (***Extended Data Fig. 3F, G***), but caused, similar as for the IPIs (**Fig. 2**), dramatic bidirectional effects on the development of kinematic stereotypy. After STN^ATN^ neuron silencing, animals developed highly stereotyped kinematics remarkably early, reaching their maximum correlation levels already after about a third of the training period (2^nd^ trial block), while controls reached similar levels only after about twice as much training (**Fig. 4B**). In contrast, after STN^DLS^ neuron silencing, the developing movement patterns remained variable throughout training, resulting in no apparent increase in kinematic correlations and significantly lower levels than in controls in late training stages (**Fig. 4B**). Notably, consistent with our findings on the IPIs (**Fig. 3**), these effects were partially dependent on trial outcome and more pronounced after rewarded trials. Despite similar trends, STN^ATN^ neuron silencing did not cause significant changes compared to controls after unrewarded trials (Fig. 4C *and see **Extended Data Fig. 9A** for all trials combined*). However, STN^DLS^ neuron silencing led to overall lower correlation levels throughout training also after unrewarded trials (**Fig. 4C**).

**Figure 4.**
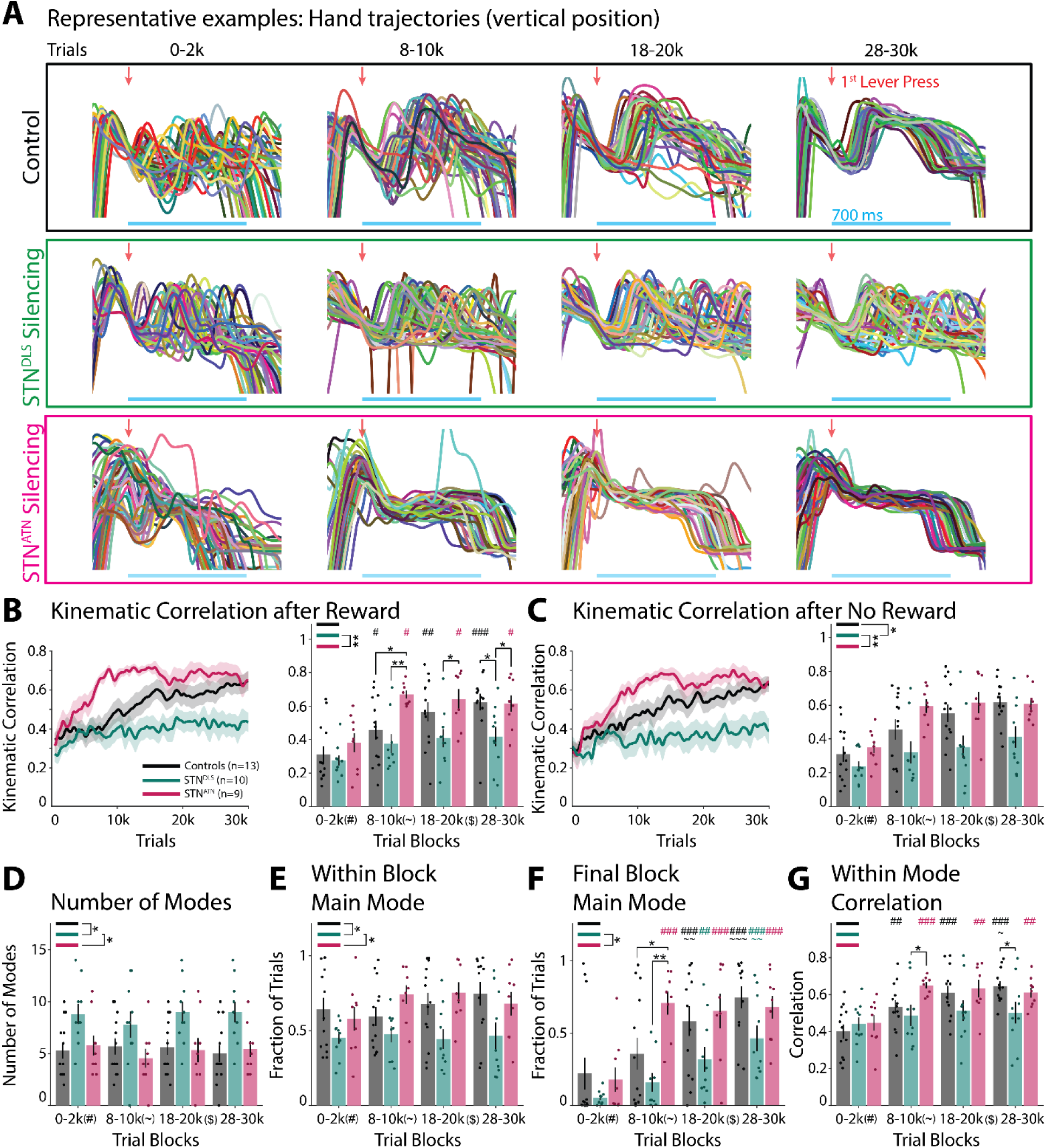
STN^DLS^ and STN^ATN^ neurons bidirectionally contribute to the regulation of kinematic variability. **(A)** Hand movement trajectories (vertical position; hand performing 1^st^ lever press) of representative rats for control and STN^DLS^ and STN^ATN^ neuron silencing in select trial blocks across learning (50 trajectories each). **(B)** Pairwise trial-by-trial correlations between hand trajectories (linearly time-warped, averaged across hands) for all trials after rewarded trials. Left: Population correlations across learning. Right: Mean correlations in trial blocks. **(C)** As (B), but for all trials after unrewarded trials. See **Extended Data** Fig. 9A for all trials combined. **(D)** Number of identified modes, i.e., distinct kinematic solutions to the task (see text for details), in trial blocks. **(E)** Fraction of trials belonging to the most used (main) mode in each trial block. **(F)** Fraction of trials belonging in each trial block to the mode most used at the end of training (main mode in last trial block). **(G)** Weighted average of within-mode kinematic correlations. The average pairwise trial-by-trial kinematic correlations for each mode were averaged for each animal, weighted by mode size in each trial block. Statistics: (B-G) Linear mixed-effects model (LME), with rats as random intercepts. Estimated marginal means from the LME, with Holm correction. For statistical details, see **Supplementary Table 3.** Bars show means, dots show individual animals, and error bars show SEM. Stars and symbols indicate significance as in Fig. 2.

Motor variability in our task is not only reflected by the kinematic similarity across individual trials, but also by the distinct kinematic solutions animals develop. Due to the lack of kinematic constraints, rats often develop multiple related, but qualitatively distinct solutions or ‘modes’ to generate the target IPI^33,34^ (see Methods). Higher levels of motor variability may lead to a broader exploration of distinct ways to solve the task, resulting in a larger number of modes, reduced favoring of particular modes and later settling on a final mode. Notably, the number of modes and the size of the largest mode in a given block remained constant over the course of training, both in controls and after STN projection neuron silencing (**Fig. 4D, E**). After STN^DLS^ neuron silencing, however, animals used significantly more modes and focused less on a particular mode at any given time throughout training (**Fig. 4D, E**). While STN^ATN^ neuron silencing had no overall effects on these measures (**Fig. 4D, E**), animals settled earlier than controls on the mode they favored at the end of training (**Fig. 4F**). Finally, after STN^ATN^ neuron silencing, the trial-by-trial correlations within individual modes also plateaued earlier than in controls, while they did not increase over the course of training after STN^DLS^ neuron silencing (**Fig. 4G**). This suggests that STN^DLS^ neuron silencing leads to the development of more modes that do not become as refined as the modes in controls, while STN^ATN^ neuron silencing leads to an earlier favoring of a particular mode and an earlier increase in mode stereotypy.

Taken together, our findings show that STN^DLS^ and STN^ATN^ neurons contribute to the regulation of motor variability across multiple task dimensions during *de novo* motor skill learning, including not only the duration, but also the kinematic details of the generated movement patterns.

### DMS-projecting ATN neurons promote motor variability

Our results show bidirectional contributions of STN^DLS^ and STN^ATN^ projection neurons to the regulation of motor variability during *de novo* motor skill learning. STN^DLS^ neurons may directly modulate the precise fine-scaled DLS activity that drives the generation of the stereotyped movement patterns^33,71,73,74^. In contrast, it is less clear how STN^ATN^ projection neurons may contribute to variability regulation. Besides a potential role in mediating symptoms in a rodent PD model^64^, the ATN, commonly associated with the hippocampal system^90–92^, have not been implicated in motor behavior. However, previous studies^65–67^ and our own anatomical tracing (**Fig. 1C**) show that the ATN directly project to the DMS, suggesting that STN^ATN^ neurons may modulate motor variability by influencing a downstream projection from the ATN to the DMS (**Fig. 1F**).

Using the same retrograde viral targeting approach as before, we chronically silenced ATN^DMS^ neurons in a separate cohort of rats (***Extended Data Fig. 10A-C*** *for injection sites*) and trained them on our motor skill paradigm. Consistent with our hypothesis, this manipulation affected performance and, in particular, the development of variability in the same way as STN^ATN^ neuron silencing (**Fig. 5B-G** *and cf.* **Fig. 2**). In addition to generating IPIs with a longer average length earlier in training than controls (**Fig. 5C**), these rats reduced the variability of their IPIs significantly faster (**Fig. 5D**), and reached high levels of trials close to the target earlier in training (**Fig. 5E**). Like STN^ATN^ neuron silencing, ATN^DMS^ neuron silencing did not change the final levels of performance measures at the end of training but accelerated when they were reached (**Fig. 5C-E**). Furthermore, the effects of ATN^DMS^ neuron silencing showed the same dependency on trial outcome, with a stronger reduction of variability after rewarded than after unrewarded trials in high reward contexts (**Fig. 5H** *and **Extended Data Fig. 9B-F** and cf.* **Fig. 3**). In contrast to STN^ATN^ neuron silencing, however, ATN^DMS^ neuron silencing did not affect the development of kinematic stereotypy (**Fig. 5I** *and **Extended Data Fig. 9B-G***). Despite this difference, our results show that silencing of ATN^DMS^ neurons largely recapitulates the effects of STN^ATN^ neuron silencing on the regulation of motor variability.

**Figure 5.**
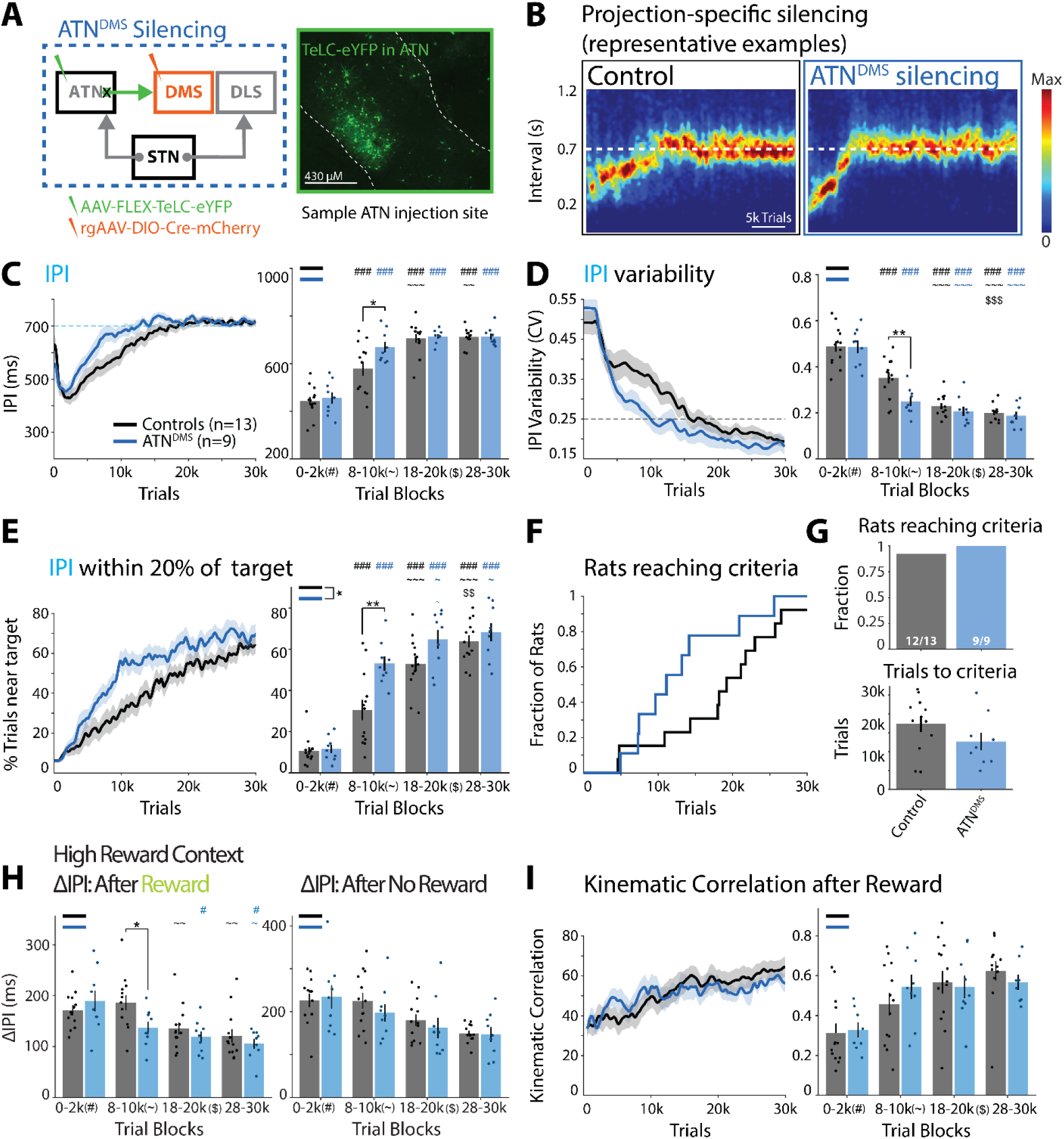
Silencing ATN^DMS^ neurons mimics effects of STN^ATN^ neuron silencing on variability regulation. **(A)** Left: Viral approach to chronically silence ATN^DMS^ neurons. Right: Representative image of TeLC expression in the ATN. See **Extended Data** Fig. 10A**-C** for injection sites. **(B)** Representative example of task performance throughout training as an IPI probability density plot. **(C)** Left: mean IPI across training. Right: Quantification of mean IPI in select trial blocks. **(D)** As (C), but for CV of the IPIs. **(E)** As (C), but for fraction of trials close to the target. **(F)** Fraction of rats reaching learning criteria across learning. **(G)** Top: Fraction of rats that reached learning criteria. Bottom: Average number of trials to reach learning criteria. **(H)** Mean ΔIPI (absolute IPI difference between trigger and subsequent trial; cf. Fig. 3A) across trial blocks after rewarded (left) or unrewarded (right) trials in a high reward context (see **Extended Data** Fig. 8E, F for low and all reward contexts). **(I)** Pairwise trial-by-trial correlations between hand trajectories (linearly time-warped, averaged across hands; see Methods) for all trials after rewarded trials. (See **Extended Data** Fig. 9B, C for trials after unrewarded or all trials.) Left: Population correlations across trials. Right: Mean correlations in select trial blocks. Statistics: (C-E; H) 2-way RM ANOVA with post-hoc tests (Holm). (G) 1-way ANOVA. (I) Linear mixed effects model (LME), random intercepts per rat. For statistical details, see **Supplementary Table 4.** Bars show means, dots show individual animals, and error bars show SEM. Stars and symbols indicate significance as in Fig. 2.

### A disynaptic STN-ATN-DMS pathway exists and promotes motor variability

Our findings are consistent with the existence of a continuous functional STN-ATN-DMS pathway that allows the STN to regulate motor variability by indirectly influencing the DMS. However, to our knowledge, such a disynaptic pathway has so far not been anatomically identified. Alternatively, our results would be consistent with a more complex connectivity, including additional synaptic connections in the ATN (e.g., a STN-ATN-ATN-DMS pathway) or other brain areas. We therefore used a three-component viral targeting strategy to selectively label and silence potential ATN neurons that both receive STN input and project to the DMS (^STN^ATN^DMS^ neurons): We injected an AAV1 for anterograde, transsynaptic^93–95^ expression of FlpO recombinase into the STN, a retrograde AAV for FlpO-dependent expression of Cre recombinase into the DMS, and a local AAV for Cre-dependent expression of TeLC-eYFP into the ATN (**Fig. 6A***, left*). This intersectional approach revealed fluorescently labeled neurons in the ATN, for the first time demonstrating the existence of ^STN^ATN^DMS^ neurons (**Fig. 6A***, right, and **Extended Data Fig. 10D-F***).

**Figure 6.**
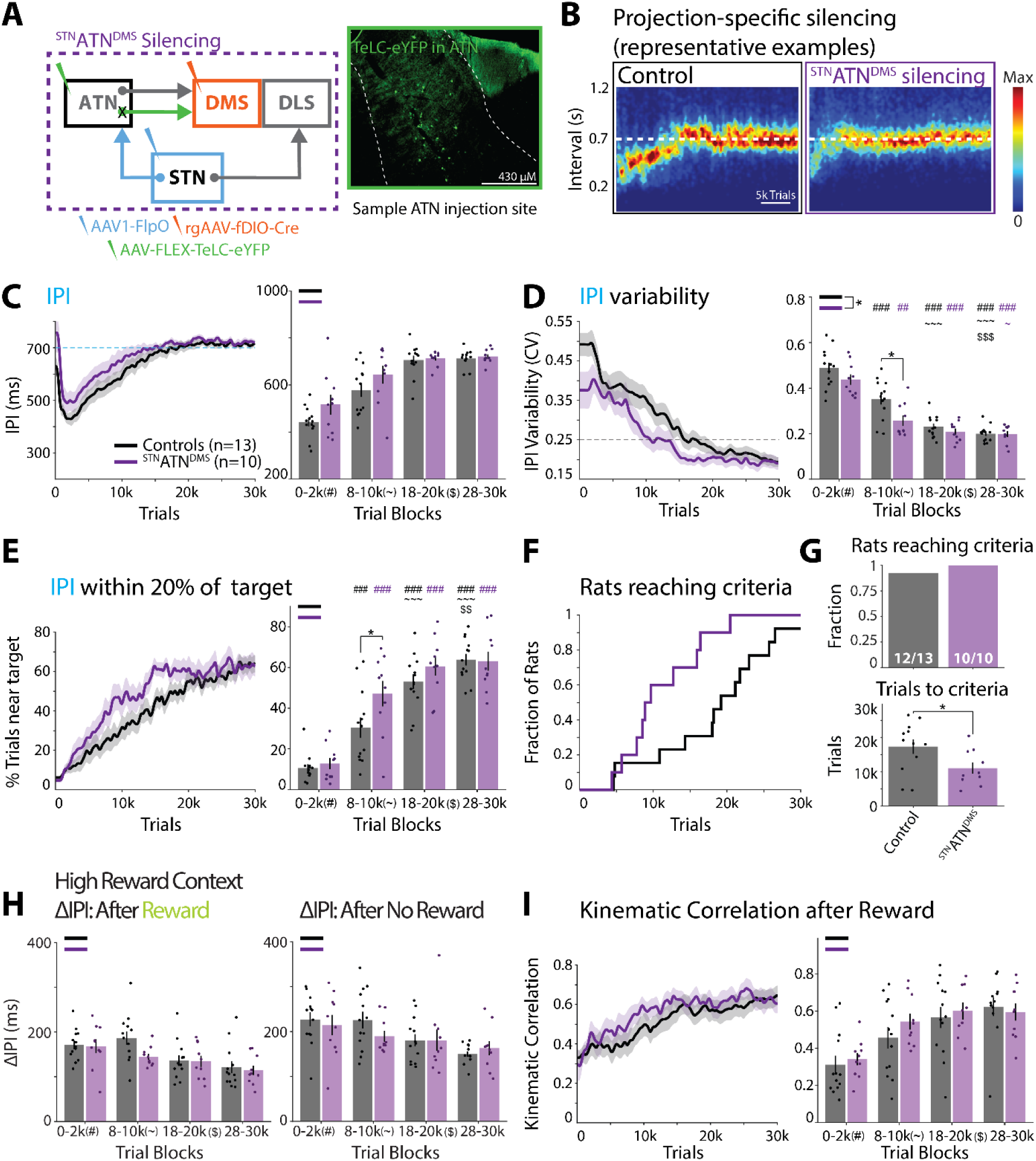
Silencing ^STN^ATN^DMS^ neurons mimics effects of STN^ATN^ neuron silencing on variability regulation. **(A)** Left: Viral approach to chronically silence input-and projection-defined ^STN^ATN^DMS^ neurons. Right: Representative image of TeLC expression in the ATN. See text for details and **Extended Data** Fig. 10D**-F** for injection sites. **(B)** Representative example of task performance throughout training as an IPI probability density plot. **(C)** Left: mean IPI across training. Right: Quantification of mean IPI in select trial blocks. **(D)** As (C), but for CV of the IPIs. **(E)** As (C), but for fraction of trials close to the target. **(F)** Fraction of rats reaching learning criteria (see text) across learning. **(G)** Top: Fraction of rats that reached learning criteria. Bottom: Average number of trials needed to reach learning criteria. **(H)** Mean ΔIPI (absolute IPI difference between trigger trial and subsequent trial; cf. Fig. 3A) across select trial blocks after rewarded (left) or unrewarded (right) trials in a high reward context (see **Extended Data** Fig. 8I, J for low and all reward contexts). **(I)** Pairwise trial-by-trial correlations between hand trajectories (linearly time-warped, averaged across hands; see Methods) for all trials after rewarded trials across learning. (See **Extended Data** Fig. 9G, H for trials after unrewarded or all trials.) Left: Population correlations across trials. Right: Mean correlations in select trial blocks. Statistics: (C-E; H) 2-way RM ANOVA with post-hoc tests (Holm). (G) 1-way ANOVA with post-hoc tests (Holm). (I) Linear mixed effects model (LME), random intercepts per rat. For statistical details, see **Supplementary Table 5.** Bars show means, dots show individual animals, and error bars show SEM. Stars and symbols indicate significance as in Fig. 2.

When we trained these rats in our motor learning paradigm, we found that ^STN^ATN^DMS^ neuron silencing recapitulated the main effects of silencing STN^ATN^ (**Fig. 2**) or ATN^DMS^ (**Fig. 5**) neurons: The reduction of IPI variability was significantly faster than in controls (**Fig. 6D**), and rats reached high levels of trials close to the target as well as our learning criteria earlier than controls (**Fig. 6E-G**). These effects, however, did not depend on trial outcome (**Fig. 6H**, *and **Extended Data Fig. 8I-L***), and like ATN^DMS^ neuron silencing, we did not observe changes in the development of kinematic stereotypy (**Fig. 6I** *and **Extended Data Fig. 9H-M***). While these findings do not exclude contributions of additional circuit elements, they show that a disynaptic pathway from the STN via the ATN to the DMS exists, and that ^STN^ATN^DMS^ neurons promote variability in *de novo* motor skill learning.

## DISCUSSION

We identified two anatomically and functionally distinct STN populations, defined by their axonal targeting of either the DLS or the ATN, and showed that they contribute to the bidirectional regulation of motor variability in a trial-outcome dependent manner (***Figs. 1-4***). While STN^DLS^ neurons promote development of stereotypy, STN^ATN^ neurons promote variable behavior and kinematics (***Figs. 2-4***). While the former are necessary to achieve high levels of stereotypy in late stages of training, the latter promote high variability early in training, favoring the initial exploration of the motor space to find suitable task-specific kinematic solutions. We further identified a novel, anatomically and functionally continuous, disynaptic pathway, connecting the STN via the ATN to the DMS, which may allow STN^ATN^ neurons to enact their influence on motor variability (**Figs. 5 *, 6***). Together, our results reveal critical roles for the STN and ATN as players in novel circuit mechanisms for the regulation of motor variability during *de novo* skill learning in the mammalian brain. These results shine a new light not only on the regulation of a critical aspect of motor learning, but also on the complex connectivity and function of the STN, the basal ganglia, and distributed subcortical networks.

### Potential mechanisms for the contributions of STN projection neurons to motor variability regulation

While previously no circuit mechanism for the regulation of motor variability in the mammalian brain had been described, transitions of behavioral influence between the DMS, associated with variable, flexible behavior^34,68–70^, and the DLS, associated with stereotyped behavior^33,34,69^, are likely candidates for contributing to this regulation. While DMS and DLS mainly receive cortical^68,96^ and thalamic^34,97^ input, our results are consistent with a concerted modulation of their activity by the STN, contributing to regulating their balance^96,98–101^ and, in turn, motor variability. The STN receives broad cortical and subcortical input^58,102–105^, potentially allowing for an integration of striatal output with cortical motor plans, reward history, and internal and external states. Based on these integrated inputs, the STN might provide differential excitatory drive to DMS and DLS to modulate their balance and promote variable or stereotyped behavior, respectively. However, the intricacies of the suggested STN projection pathways suggest more complex and possibly distinct influences on DMS and DLS activity.

In the DLS, the STN preferentially targets parvalbumin positive inhibitory interneurons (PV neurons)^62^, suggesting that it may not simply drive overall DLS activity, but modulate its output. Driving PV neurons might suppress DLS activity broadly, thereby favoring the generation of previously reinforced activity patterns that underlie well-practiced movement patterns over other kinematic solutions. Alternatively, STN input may support PV neurons in sharpening the firing windows and promoting precise spiking activity of their target neurons^106–109^. Thereby, STN may support the generation of the precise DLS firing patterns that drive the generation of stereotyped kinematics in learned skills^33,71,73,74^. These possibilities are not mutually exclusive, and they both suggest a larger importance of STN’s influence on DLS activity late in training, when task-specific movement patterns have been developed and the DLS drives stereotyped behavior – consistent with our findings that silencing of STN^DLS^ neurons affects motor variability late, but not early in learning (***Figs. 2-4***).

How the STN may influence DMS activity and motor variability via the novel STN-ATN-DMS pathway is less clear. Our finding that STN neurons that directly project to the DMS do not contribute to variability regulation (***Extended Data Fig. 5***), suggests that the ATN are not merely a relay, but provide critical information. While this is, to our knowledge, the first study to implicate the ATN in motor behavior, their intimate connections with the hippocampal system^90–92^ suggest that they may contribute information about general task context. The degree of familiarity with the current context likely influences the level of uncertainty regarding task and reward structure, which in turn, influences overall levels of motor variability, with higher uncertainty promoting higher variability^3^. Such context information may be integrated with STN inputs carrying information about striatal output and reward history to favor high or low variability levels. Nevertheless, it also remains unclear how the ATN, and specifically, ATN neurons that receive STN input, may modulate DMS activity. While the ATN have been shown to target spiny projection neurons (SPNs) and PV neurons^110,111^, but not cholinergic interneurons^66^ in the DMS, the specific targets of the STN-ATN-DMS pathway have not been explored. In a simple model, depending on the familiarity of the current task context and reward history, this pathway might provide more or less excitatory drive to SPNs to support the DMS in outcompeting the DLS^96,98–101^ and in driving variable, exploratory behavior. This would suggest a preferential role for this pathway under high levels of uncertainty, e.g., at the beginning of task training – consistent with our finding that silencing of this pathway leads to an accelerated reduction of variability early in learning, without changes in final variability levels (***Figs. 2-6***).

### Variability regulation by the STN-ATN-DMS pathway might have additional complexity

While we show that an anatomically continuous STN-ATN-DMS pathway exists (**Fig. 6**), and that silencing nodes in this pathway causes largely consistent effects on motor variability regulation (***Figs. 2-6***), silencing of ATN^DMS^ neurons or of ^STN^ATN^DMS^ neurons does not recapitulate the accelerated development of kinematic stereotypy we observed after STN^ATN^ neuron silencing (***Figs. 4-6***). This could be due to a low yield of infected ^STN^ATN^DMS^ neurons in both intersectional silencing experiments (**Figs. 5 *, 6*** *and **Extended Data Fig. 10***). Alternatively, another population of ATN neurons that receive STN input, but do not project to the DMS, might specifically contribute to the regulation of kinematic stereotypy. A candidate target for such a hypothetical STN-ATN-X pathway could be the DLS^65^. A sparse projection from the ATN targets the DLS^65^, where it could, for example, drive inhibitory interneurons to suppress DLS activity underlying kinematic stereotypy, acting in concert with ATN^DMS^ neurons promoting DMS activity. We note, however, that it remains unclear whether DLS-projecting ATN neurons receive STN input and which neurons they target in the DLS. Further anatomical tracing and functional experiments will be necessary to distinguish between these possibilities and to identify potential other relevant targets of STN-targeted ATN neurons.

### Limitations

While our chronic silencing approach allowed us to study the role of projection neurons that are defined by a specific projection target over weeks of motor learning, we note its technical limitations. First, since this approach also affects potential axon collaterals of the infected neurons, we cannot fully exclude that effects on other target brain areas than the intended ones have contributed to our findings. Importantly, however, our careful control experiments at least partially mitigate this caveat. We showed that STN^DLS^ and STN^ATN^ neurons are anatomically and functionally distinct from each other (**Fig. 1D, E**) and from STN neurons that project to the DMS, or to the thalamic rILN or RTN (***Extended Data Figs. 4-7***), excluding major targets for potential axon collaterals that might contribute to behavioral effects. Nevertheless, we cannot fully exclude contributions of potential axon collaterals to other brain areas, e.g., to other thalamic nuclei or the Globus Pallidus. We note, however, that it has been reported that STN’s projection to the ATN is distinct from its projection to the Globus Pallidus internus^64^. Due to temporal limitations of chronic silencing, we could also not fully determine the timescale of motor variability regulation. While our results suggest that the STN projection pathways contribute to trial-to-trial variability regulation in an outcome-dependent manner (**Figs. 3 *, 4***), they might also contribute to prolonged high or low variability states. Future experiments will be needed to overcome spatial and temporal limitations of our approach, e.g., by using acute manipulations of axon terminals in select target regions.

## Conclusions

Our results have revealed novel circuit mechanisms in the mammalian brain for the regulation of motor variability, consistent with a model in which the STN modulates DMS and DLS activity in concert to favor variable or stereotyped motor behavior. Future experiments will be necessary to reveal the detailed impact of STN’s projections on striatal activity and how they themselves are driven and coordinated. Our results also raise the question whether these mechanisms are limited to motor variability during *de novo* motor learning or whether they also contribute to motor adaptation of learned skills and more broadly to the balance between exploration and exploitation across different forms of behavior.

## METHODS

### Animals

The care and experimental treatments of all animals were reviewed and approved by the University of Maryland Baltimore Institutional Animal Care and Use Committee. For all experiments, subjects were female, wild-type Long Evans rats (Charles River). For behavioral experiments, rats were 10-12 weeks old at start of training (n = 84, 8-10 per group). Because the behavioral effects of our treatments could not be specified in advance, we chose sample sizes that would allow us to identify outliers and validate reproducibility. For histological tracing experiments, rats underwent stereotaxic surgery at 10-16 weeks of age (n = 9).

### Projection-Specific Silencing of Synaptic Transmission

To study how the loss of specific projection neuron populations, defined by their projection targets, affects motor skill learning, we used a two-component viral strategy for selective chronic neuronal silencing in naïve rats prior to training on a motor skill learning paradigm (see ‘Behavioral Training’). To selectively silence neurons that send axons to a target area of interest (e.g., STN neurons that send an axon to the DLS; see ***Table 1*** for all probed projection neurons), we combined retrograde viral infection with Cre-dependent expression. In the target area (see ***Table 2*** for injection coordinates), we injected a retrogradely infecting adeno-associated virus (rgAAV) which can infect neurons by entering axon terminals. This rgAAV drove the expression of Cre recombinase and the fluorophore mCherry (rgAAV-EF1a-mCherry-IRES-Cre, AddGene). In the source area, we injected a locally infecting AAV for Cre-dependent conditional expression of Tetanus Toxin Light Chain (TeLC) coupled to eYFP (AAV8 or AAV5-FLEX-TeLC-eYFP; DNA construct from AddGene 135391 and custom virus production by University of Maryland Baltimore School of Medicine viral vector core). To mark both injection sites, we co-injected a small amount of Fluorogold (1:8 in the virus solution; Biotium). TeLC cleaves the synaptic protein VAMP2, thereby preventing the fusion of synaptic vesicles with the membrane and the release of neurotransmitters, effectively and chronically silencing synaptic transmission in infected neurons^77–79^. While the locally infecting AAVs nonspecifically infect neurons in the source region, TeLC is only expressed in neurons expressing Cre, i.e., neurons that are also infected by the retrograde rgAAV. This strategy therefore allowed us to target STN or ATN projection neurons that send an axon to our target region of interest. To control for potential axon collaterals and potential spillover of viral injections into neighboring thalamic nuclei, we probed additional projection neurons, including STN neurons projecting to the thalamic rILN or RTN (***Extended Data Figs. 6, 7***). The injections into these thalamic nuclei were mainly targeted to the parts immediately adjacent to the ATN which were most likely to be affected by potential viral spillover (***Extended Data Figs. 1, 6***). To control for the effects of surgeries, viral infections, and expression of recombinant proteins, we performed the same injections but using an AAV for expression of GFP (AAV5-hSyn-DIO-eGFP; Addgene) instead of TeLC-eYFP. Injections were performed as previously described^33,34^. Briefly, rats were anesthetized with 2% isoflurane and placed in a stereotactic frame (Kopf; Model 1900). After incision of the skin and skull cleaning, bregma was located and small craniotomies were performed above the target regions. A thin glass pipette connected to a Nanoject II microinjector (Drummond) was lowered through the craniotomies to the injection sites and viruses were injected in 10 nl increments (see ***Table 2*** for injection coordinates and volumes). For certain injections, the pipette was slowly retracted across the injection range. After the injections, the pipette was retracted by ∼100 μm and left there for at least 3 min to allow for diffusion and to prevent backflow of the virus. After all injections, the skin was sutured and rats received painkillers (Carprofen, 5 mg/kg for 3 days). Surgeries were performed in naïve rats and they were given 7 days to recover prior to the start of training. Correct targeting and spread of injections were determined post hoc (see ‘Histological Analysis’).

**Table 1.** Targeted projection neurons for behavior groups, respective virus injection sites and associated figures.

| Projection | Target<br>rgAAV Injection Site | Source<br>TeLC/GFP Injection Site | Anterograde (AAV1) Injection Site | Figures |
|---|---|---|---|---|
| STN<sup>DLS</sup> | DLS | STN |  | 1-4, S1-4, S8-9 |
| STN<sup>ATN</sup> | ATN | STN |  | 1-4, S1-4, S8-9 |
| ATN<sup>DMS</sup> | DMS | ATN |  | 5, S8-10 |
| <sup>STN</sup>ATN<sup>DMS</sup> | DMS | ATN | STN | 6, S8-10 |
| STN<sup>DMS</sup> | DMS | STN |  | S4-6 |
| STN<sup>RTN</sup> | RTN | STN |  | S6-7 |
| STN<sup>rILN</sup> | rILN | STN |  | S6-7 |

**Table 2.** Injection site coordinates and volumes. Anterior-Posterior (A-P) and Medial-Lateral (M-L) coordinates measured from Bregma; Dorsal-Ventral (D-V) measured from the top of the brain.

| Injection<br>Site | A-P<br>(mm) | M-L<br>(mm) | D-V<br>(mm from brain) | Volume<br>(nL) |
|---|---|---|---|---|
| DLS1 | +1.00 | +/-3.20 | -4.1→-3.6 | 290 |
| DLS2 | +0.40 | +/-3.80 | -3.9→-3.2 | 300 |
| DLS3 | -0.60 | +/-4.10 | -4.2→-3.2 | 350 |
| ATN1 | -1.80 | +/-1.40 | -4.8 | 70 |
| ATN2 | -1.80 | +/-0.70 | -5.6 | 40 |
| STN | -3.50 | +/-2.40 | -7.325 | 40 |
| DMS1 | +0.50 | +/-1.90 | -4.0→-3.0 | 350 |
| DMS2 | +0.10 | +/-1.90 | -4.5→-3.5 | 350 |
| DMS3 | -0.40 | +/-2.20 | -4.1→-3.1 | 350 |
| rILN | -1.90 | +/-0.60 | -4.750 | 120 |
| RTN | -1.60 | +/-0.95 | -5.8→-5.1 | 100 |

### Projection-Specific Silencing of Disynaptic Circuits

To silence ATN neurons that both receive STN input and project to the DMS (^STN^ATN^DMS^) we used a three-component viral strategy, combining antero- and retrograde viral approaches (see ***Tables 1, 2*** for injection sites and coordinates). In the STN, we injected an AAV1 for the anterograde, transsynaptic expression^93–95^ of FlpO recombinase (AAV1-CAG-FlpO; Addgene), and in the DMS we injected a retrograde AAV for FlpO-dependent expression of Cre (rgAAV-fDIO-Cre; Addgene). Together with the injection of an AAV for Cre-dependent TeLC expression (see above) in the ATN, this strategy allowed for expression of Cre and, in turn, of TeLC in DMS-projecting, STN-targeted ATN (^STN^ATN^DMS^) neurons. All injections were performed in the same surgery, and animals were allowed to recover for 7 days to allow for viral expression before the start of training.

### Circuit Tracing

To determine the potential overlap between different STN projection neuron populations, we performed dual-site retrograde viral tracing. Naïve rats were injected as described above with two retrograde rgAAVs for expression of tdTomato (rgAAV-CAG-tdTomato; Addgene) or GFP (rgAAV-CAG-GFP; Addgene) into two distinct target areas (e.g., ATN and DLS; n=6 hemispheres/ projection pair). After 3 weeks, animals were perfused, brains were extracted and processed, and labeling overlap was analyzed (see ‘Histological Analysis’). See ***Table 2*** for injection coordinates.

### Histological Analysis

At the end of training (behavior groups) or 3 weeks after surgery (circuit tracing), rats were given inhalant anesthesia (5% isoflurane) and perfused with phosphate-buffered saline (PBS) followed by 4% paraformaldehyde (PFA) in PBS. Their brains were harvested for histological analysis, sectioned into 65 μm coronal slices on a vibratome (Leica VT1000S), and underwent immunofluorescent staining at room temperature to determine the extent and location of injection sites and TeLC-eYFP expression. Sections were placed on a rocker for 2 hours in blocking solution (BS; 1% bovine serum albumin and 0.3% Triton-X 100 in PBS) and subsequently incubated overnight on a rocker with primary antibodies for GFP (1:1000 in BS, Invitrogen, A11122, raised in rabbit), RFP (1:1000 in BS, Sigma-Aldrich, AB3528, raised in chicken), and Parvalbumin (PV, to stain thalamic and subthalamic landmarks; 1:1000 in BS; Sigma-Aldrich, P3088, raised in mouse). After subsequent washing (3x 10 min in PBS), sections were incubated for 2 hours with fluorescently labeled secondary antibodies (1:1000 in BS; anti-rabbit Alexa Fluor 488, Invitrogen, A11008; anti-chicken Alexa Fluor 555, Invitrogen A21437; anti-mouse AlexaFluor 647, Invitrogen A21236). Sections were then washed (3x 10 min in PBS) and mounted with Fluoromount-G mounting media (Invitrogen).

### Histology: Behavior Experiments

Using a BZ-X710 All-in-One Fluorescence Microscope (Keyence) or Ti2-E microscope (Nikon), images of rgAAV injection sites were acquired at 4x magnification (5-6 whole brain slices/animal) and Z-stacks of TeLC-eYFP injection sites at 10x magnification (3 slices/animal). To determine and compare the extent and localization of the rgAAV injections across all animals, we used the DMC-BrainMap extension in Napari^112^ to register and segment whole brain images. First, we identified fluorescently labeled neurons (mCherry expression from the rgAAV-Cre-mCherry or co-injected Fluorogold for RTN injections) using the Rab5a model from the Allen Cell and Structure Segmenter^113^. After automatic detection of objects with cell size and shape above a set fluorescent threshold, we manually removed false positive objects in regions distant from the injection site. The brain slices with identified putative neurons were then registered to a common rat brain atlas (Waxholm Space Atlas of the Sprague Dawley Rat Brain; whs_sd_rat_39um), allowing for direct comparisons across rats and for localization of identified neurons to individual brain areas. The segmented and registered slices were then exported to Adobe Illustrator and overlaid for each behavioral group (see ***Extended Data Figs. 1, 6, 10***). To quantify the extent of TeLC-eYFP expression, fluorescently labeled neurons were manually counted using ImageJ cell counter. Rats with mis-targeted injections were excluded from behavioral analysis.

### Histology: Circuit Tracing

Images of rgAAV injection sites were acquired as above and targeting was confirmed visually. Hemispheres with mis-targeted injections were excluded from analysis. In hemispheres with confirmed injection sites, Z-stacks of the STN were taken at 10x magnification and total RFP-positive, GFP-positive, and overlap neurons were manually counted using ImageJ.

### Behavioral Training

Rats were trained on a lever-pressing task using a fully automated home-cage training system, as previously described^76,80^. In short, after initial acclimatization and pre-training, water-restricted rats were rewarded with water for pressing a lever twice within performance-adjusted boundaries around a fixed inter-press interval (IPI) target of 700 ms. Reward boundaries were initially set to 200 ms and 1100 ms. To shape the animals’ behavior toward the target interval while maintaining their motivation to perform the task, the boundaries were dynamically and automatically adjusted between sessions, tightening when the reward rate exceeded 40% and expanding when it was less than 30%. To further incentivize IPIs close to the target, a linearly scaled reward landscape with five evenly spaced levels was established between the target (highest reward, 100 µL water) and each boundary (lowest reward, 20 µL water). Intervals outside the boundaries were not rewarded and triggered a time-out in which animals had to withhold pressing for at least 1.2 s before initiating a new trial (Inter-trial interval; ITI). Over weeks of training, control rats developed increasingly complex and stereotyped movement kinematics between lever presses (see ‘Kinematic Tracking’), allowing them to maintain the prescribed IPI target with increasing accuracy and precision. We considered animals as ‘experts’ in the task, i.e., having learned the task structure and stabilized performance, when they met our previously established learning criteria^76^ (see ‘Criteria Performance’).

### Kinematic Tracking

Over the course of training, rats developed increasingly complex, spatio-temporally precise, and stereotyped *de novo* movement patterns which span the interval between the lever presses as solutions to our task^33,34,76^. Raspberry Pi cameras mounted on both sides around the lever captured high-speed videos (90 Hz) of all lever-press-related movements throughout training. The recorded videos were automatically cut into 2-second clips and aligned to the first lever press in each trial of the task. To track the learned movement patterns and their development over the course of training, we took advantage of machine learning-aided pose estimation, similar to our previous work^33,34^. In short, we used the pose estimation software SLEAP^89^ to train a deep neural network on the position of multiple body parts (hands, elbows, ears, nose) on ∼200 video frames per camera view for each rat (sampled from timepoints throughout training). The trained network was used to predict body part positions across all frames of all trial videos for the particular rat. We used a Kalman filter to smooth the resulting trajectories and to linearly interpolate the positions of occluded body parts for up to 5 consecutive frames. Trajectories with more than 5 consecutive occluded frames were excluded from analysis.

## Data Analysis

### Behavioral Performance Metrics

All behavioral analyses were performed using MATLAB 2024a (MathWorks). Performance metrics were determined from lever-press timing in our task. The IPI is the time between the first and second press in a trial, and the ITI is the time between the last press in a non-rewarded trial and the next lever-press. The fraction of trials close to the target (trials in the IPI range of 700 ms ± 20%) and the coefficient of variation (CV) of the IPI distribution were calculated over a moving window (100 trials), and the moving average was low-pass filtered with a 300-trial boxcar filter. Across behavioral analyses, we quantified performance in select trial blocks, based on a global trial index across all training sessions, evenly covering the entire training period: Block 1: Trials 1-2,000, Block 2: Trials 8,001-10,000, Block 3: Trials 18,001-20,000, Block 4: Trials 28,001-30,000.

### Learning Criteria

We considered animals as experts in our task when their CV was less than 0.25 and the mean of their IPI distribution was in a range of 700 ms ± 10% for a 3,000-trial sliding window. These criteria were previously established and capture the learning performance of intact animals in our _task_33,34,76.

### Propensity Score Matching Analysis

To determine the impact of trial outcome on the contributions of the different STN projection neurons, we took advantage of a propensity score matching procedure^3,88^ to account for varying reward context: Let *T_ij_* be the number of valid trials on the *j^t^*^ℎ^ training session of the *i^t^*^ℎ^ animal. We define *R^j^* ∈ {0, 1}, a binary reward variable that takes 0 if the rat was unrewarded on trial t, and 1 otherwise (pooling all reward sizes). For each training session, we fit a logistic regression model, as previously described by Dhawale et al.^3^ (equation (9)) to predict the probability of receiving a reward on trial t, based on reward outcomes of past and future surroundings. We choose a surrounding index around a particular trial *S^j^* = {*t* − *p_ij_*, …, *t* − 1, *t* + 1, …, *t* + *p_ij_*}, as the reward context. Here, *p_ij_* is chosen to avoid the influence of extreme fitted logits along with an adaptive threshold of 5–20 trials (depending on the number of trials in the *j^t^*^ℎ^ training session) to overcome excess exclusion of the extremes on both ends. For the *j^t^*^ℎ^ training session of the *i^t^*^ℎ^ animal, we fit the following model with an adaptive window:

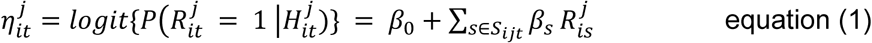

where *H^j^_it_ = {R^j^_is_: s ⋳ S^j^_it_*}, for all *t* ∈ *S^j^_it_*.

After estimating the propensities in equation (1), we focused our analysis on the same 4 trial blocks we used for our other behavioral analyses (see ‘Behavioral Performance Metrics’). In each block, trials are ordered according to their estimated probability of receiving reward. The ordered trials are divided into 10 approximately equal sized bins based on their estimated reward propensities. Within each bin, we compute the mean absolute IPI difference between each trial (the trigger trial) and the subsequent trial (ΔIPI) and the variance of IPIs (vIPI) with a moving window of sizes 2, 3, 4 and 5, separately for rewarded and unrewarded trials. For vIPIs, the moving window starts from the trigger trial. Bin-specific scores are combined by a weighted average, with weights determined by the number of rewarded and unrewarded trials in each bin. These measures characterize the trial-to-trial IPI change and the IPI variability in response to rewarded or unrewarded trials. In a separate analysis, we classified the trials according to their estimated reward probabilities. Trials with estimated probabilities of less than 0.45 or greater than 0.55 are considered to be low or high propensity, respectively. The same binning scheme and weighted scores are computed for both strata. This stratification allows us to examine whether response patterns vary similarly between rewarded and unrewarded trials under low and high propensity conditions (i.e., reward contexts).

### Movement Trajectory Analysis

All kinematic analyses were performed using Python (version 3.11.9). To determine manipulation effects on the development of stereotyped movement kinematics over the course of learning, we compared the trajectories of both hands (horizontal and vertical dimension) of all tracked animals (see ‘Kinematic Tracking’) in the same four trial blocks as for our other analyses (see ‘Behavioral Performance Metrics’). We split our analyses into trials following either rewarded or unrewarded trials. This allowed us to compare trials with the same starting positions as animals move down to, and back up from, a reward port underneath the lever after successful trials, and generally stay at the lever following unrewarded trials^33,34,76^. Furthermore, this allowed us to determine whether manipulation effects differ in response to rewards (cf. ‘Propensity Score Matching Analysis’). To be able to directly compare the kinematic details of the generated trajectories, independent of the individual trials’ IPI durations, we sub-selected trials in an IPI range of 700 ms ± 200 ms and linearly time-warped the trajectories to the same duration (700 ms) by interpolating between the lever-presses. We then calculated in each trial block pairwise trial-by-trial correlations for both hands in both the horizontal and vertical dimensions and averaged them to yield a single correlation value per block for each animal. In addition, we generated continuous correlation plots across the entire learning period by calculating pairwise trial-by-trial correlations in a sliding window of 500 trials, using the same trial selection and time-warping procedures.

### Identification of Distinct Kinematic Solutions or ‘Modes’

Individual rats often develop multiple related, but distinct movement patterns as solutions to the task, which we term ‘modes’ (for details, see^33,34^). To identify these modes, we performed unsupervised clustering of all task-related kinematics from individual rats. The kinematic data included horizontal and vertical components of position, velocity, and acceleration of both hands between 200 ms before the first and 200 ms after the second lever press for all trials with at least two lever presses. To account for trial-to-trial IPI variability, we linearly time-warped the kinematic data to the duration of the target interval (700 ms). The variance of each kinematic feature was standardized by dividing by its standard deviation estimated across all time points and horizontal and vertical components for both hands. We then performed two-dimensional uniform manifold approximation and projection (UMAP) embedding^114^ of the kinematic data associated with each trial. The UMAP embedding was created using the trial data of all main behavior groups (STN^ATN^, STN^DLS^, ATN^DMS^, and ^STN^ATN^DMS^ neuron silencing and the main control group (n = 51 rats)) to embed all trials from every rat in the same low dimensional space. Within the shared embedding, we identified the modes of each animal separately: For each animal, we extracted the low dimensional coordinates of its trials, computed all pairwise Euclidean distances, and performed agglomerative hierarchical clustering with Ward’s linkage. The dendrogram was cut with the Dynamic Hybrid Tree Cut algorithm^115^ (Python implementation: dynamicTreeCut), using a minimum cluster size of 100 trials and a deepSplit parameter of 2. Due to the shared embedding, clustering was comparable across animals, allowing us to directly compare the resulting modes.

### Mode Analysis

We quantified the number of modes for each group in select trial blocks (see ‘Behavioral Performance Metrics’). We also determined the fraction of trials belonging to the largest mode in each individual trial block as well as the fraction of trials in each block belonging to the mode that is the largest in the last training block. To quantify within-mode stereotypy, we calculated trial-to-trial correlations as above (see ‘Movement Trajectory Analysis’) for each mode used in a given trial block by an animal and then calculated the average of these correlations across all modes, weighted by the proportions of their trials in that block.

### Statistics

To test for statistical significance of the effects of projection neuron silencing, performance and kinematic measures were divided into 4 blocks of trials across training (see ‘Behavioral Performance Metrics’). For statistical comparisons of performance measures, we performed two-way repeated-measures analysis of variance (ANOVA) tests with trial block as the within-subjects factor and manipulation group (e.g., STN^DLS^ neuron silencing) as the between-subjects factor. If statistically significant group or block differences or group x block interactions were detected, we performed post-hoc tests, using the Holm correction for multiple comparisons, to identify the individual significantly different comparisons. This statistical analysis revealed that there were no significant differences across our different control cohorts (GFP expression in distinct projection neurons; ***Extended Data Fig. 2***) in any of our behavioral measures. We therefore combined them into larger control groups, i.e., we combined the STN^DLS^, STN^ATN^ and ATN^DMS^ control cohorts into a control group for our main figures. As a control group for silencing of STN^DMS^ neurons, we combined striatum-targeted control cohorts (STN^DMS^ and STN^DLS^; ***Extended Data Fig. 5***). Finally, as a comparison for the thalamic injection controls, we combined thalamus-targeted control cohorts (STN^rILN^, STN^RTN^ and STN^ATN^; ***Extended Data Fig. 7***). Due to technical challenges, we could not generate reliable trajectory data for all animals in every block, leading to a small set of missing data (n=4 data points from 4 animals missing out of a total of 128 datapoints; n=1 sham, n=2 STN^ATN^, n=1 STN^DLS^). To determine significant effects of our manipulations on our kinematic measures, we therefore used linear mixed effects models, with manipulation group and trial blocks as fixed effects, and individual animals as a random intercept. To explore significant group, block or group x block effects, we computed estimated marginal means using Satterthwaite’s approximation for degrees of freedom and performed post-hoc tests with Holm correction. For other comparisons, we used a one-way ANOVA with subsequent post hoc tests (Holm correction; e.g., **Fig. 2G**). All data is expressed as mean +/-standard error of the mean (SEM). We used the statistics software JASP (0.95.4, 2025, University of Amsterdam) for statistical analysis. Significant differences are indicated in the respective figures, and statistical details for behavioral analyses are provided in supplementary tables associated with individual figures.

## DATA AVAILABILITY

All data will be made publicly available upon publication of this manuscript.

## CODE AVAILABILITY

All analysis code will be made publicly available upon publication of this manuscript.

## ACKNOWLEDGEMENTS

We thank all members of the Wolff lab for technical support and feedback, in particular Tara Foley, Jian Kwon, and Rebecca Weimer. We thank Ramesh Chandra and the University of Maryland School of Medicine’s Viral Vector Core for help with virus production. We thank also the McCarthy and Poulopoulos labs for generously sharing their microscopes and biological illustrator Tina Wang for her contributions to Figure 2.

## AUTHOR INFORMATION

### These authors contributed equally

Oluwatomiwa A. Oladunni and Sophie K. Elvig

## Authors and Affiliations

Department of Pharmacology and Physiology, University of Maryland Baltimore School of Medicine, Baltimore, MD, USA

Oluwatomiwa A. Oladunni, Sophie K. Elvig, Hunter C. Bennis, Laila A. Elzaree, Brian N. Mathur, Steffen B.E. Wolff

Department of Mathematics and Statistics, University of Maryland Baltimore County, Catonsville, MD, USA

Siddhartha Nandy, Yuting Liu, Snigdhansu Chatterjee

University of Maryland Medicine Institute for Neuroscience Discovery, University of Maryland Baltimore, Baltimore, MD, USA

Brian N. Mathur, Steffen B.E. Wolff

## Contributions

The study was designed by O.A.O., S.K.E., and S.B.E.W. Viral injection surgeries were completed by O.A.O. and S.K.E. Behavioral training was conducted by O.A.O. and S.K.E. Tissue processing and histological analysis were completed by O.A.O., S.K.E., and L.A.E. Models for kinematic tracking were trained by O.A.O., L.A.E., and H.C.B. Kinematic analyses were completed by H.C.B. Statistical analyses were performed by O.A.O. Propensity score analyses were performed by S.N., Y.L., and S.C. Figures were composed by O.A.O, S.K.E., and S.B.E.W. The project was supervised by B.N.M. and S.B.E.W. The manuscript was written by O.A.O, S.K.E., B.N.M., and S.B.E.W. with input from all authors. All authors read and approved the final version of the manuscript.

## Corresponding Author

Correspondence to Steffen B.E. Wolff

## EXTENDED DATA FIGURES

**Extended Data Figure 1:**
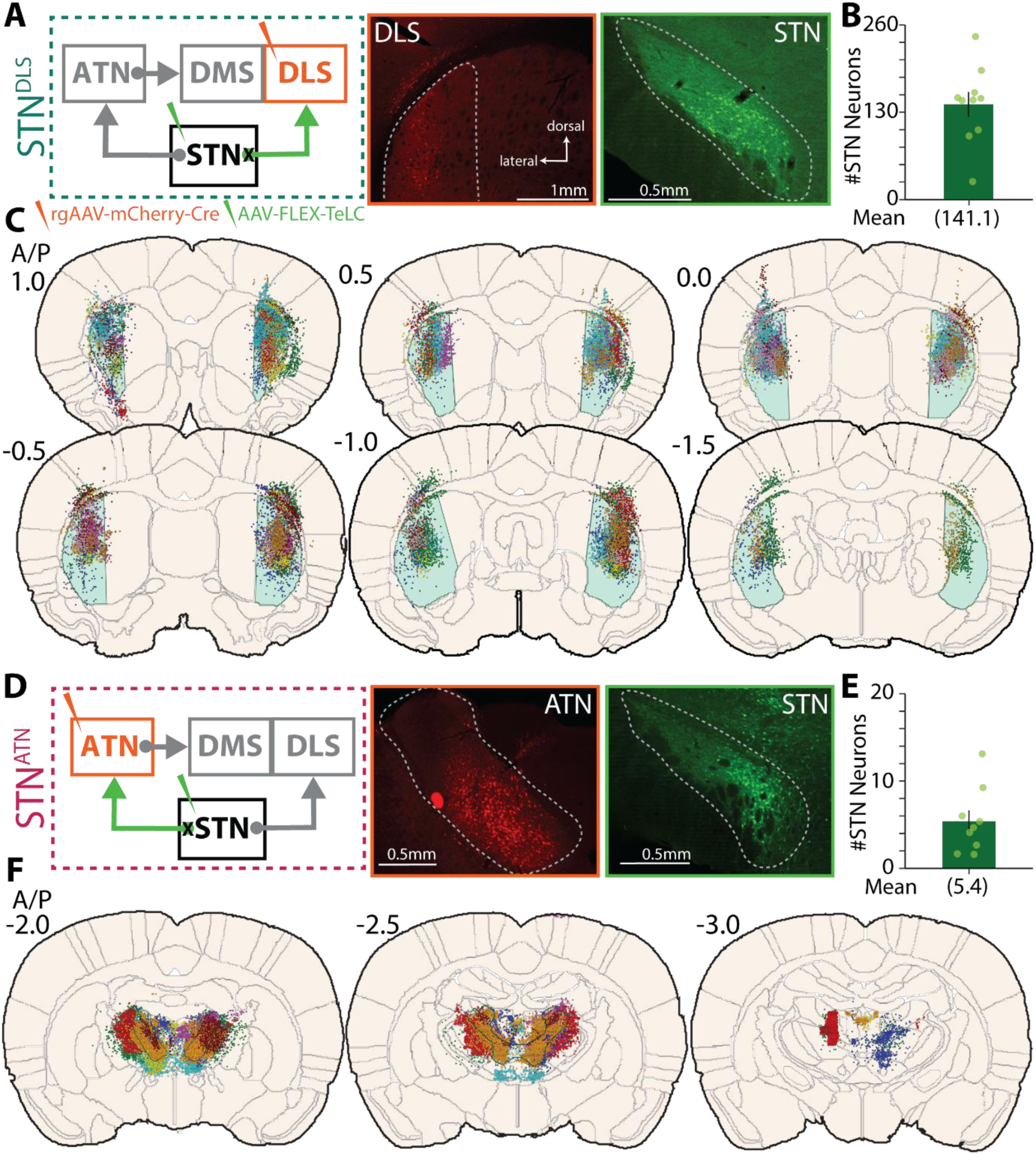
Viral injection sites for STN^DLS^ and STN^ATN^ neuron silencing. **(A)** Left: Circuit scheme for chronic silencing of STN^DLS^ neurons. Right: Representative images of viral injections in the DLS (rgAAV-mCherry-Cre) and in the STN (AAV-FLEX-TeLC-eYFP). Targeted areas are outlined by dotted white lines. **(B)** Quantification of silenced STN^DLS^ neurons per slice (n=10 animals, counts averaged between hemispheres across 5-6 slices per animal). **(C)** Overlay of viral injection sites in the DLS (light blue) for STN^DLS^ neuron silencing on schematic coronal brain sections (rgAAV-mCherry-Cre). Dots represent automatically detected fluorescent neurons (see Methods) and colors indicate individual animals. **(D-F)** As (A-C), but for STN^ATN^ neuron silencing (n=9 animals, counts averaged between hemispheres across 3 slices per animal).

**Extended Data Figure 2.**
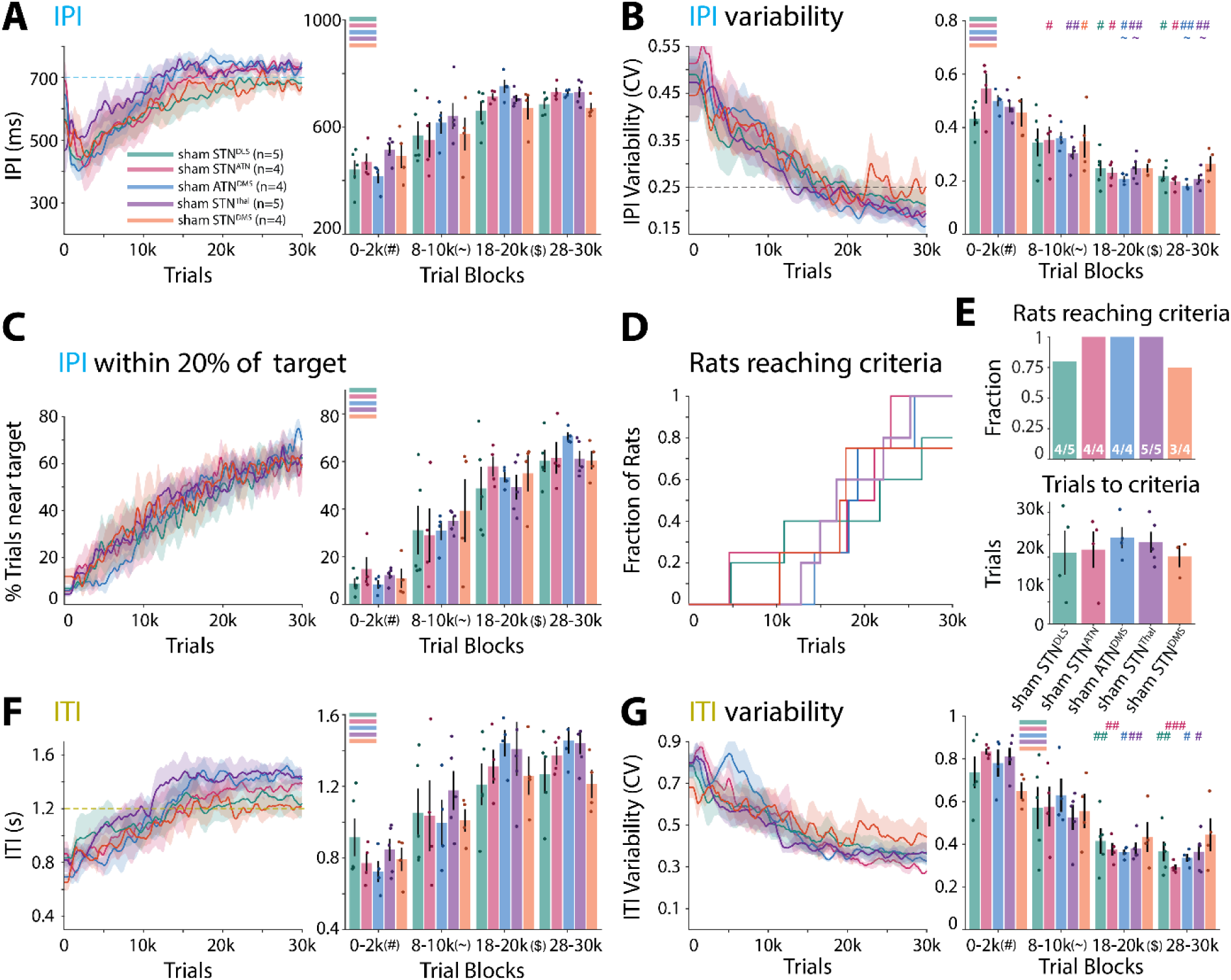
Individual control groups for silencing of distinct projection neurons do not differ in task performance. **(A)** Quantification of task performance across learning for individual control groups, expressing GFP in distinct projection neurons: STN^DLS^, STN^ATN^, ATN^DMS^, STN^DMS^, and STN^Thalamus^ (targeting the rostral intralaminar nuclei and the Reticular thalamic nucleus). Left: mean IPI across training. Right: Quantification of mean IPI in select trial blocks. **(B)** As (A), but for CV of the IPIs. **(C)** As (A), but for fraction of trials close to the target. **(D)** Fraction of rats reaching learning criteria (see text) across learning. **(E)** Top: Fraction of rats that reached the learning criteria. Bottom: Average number of trials needed to reach the learning criteria. **(F)** As (A), but for the mean Inter-Trial Intervals (ITI). After unrewarded trials, rats need to keep a minimum time-out period (1.2 s) before initiating a new trial (see Methods). **(G)** As (A), but for the CV of the ITIs. Statistics: (A-C, F-G) 2-way repeated measures ANOVA with Post-hoc tests (Holm) when appropriate. (D) 1-way ANOVA. For all statistical details, see **Supplementary Table 6**. Bars show means, dots show individual animals, and error bars show SEM. Stars and symbols indicate significance as in Fig. 2.

**Extended Data Figure 3:**
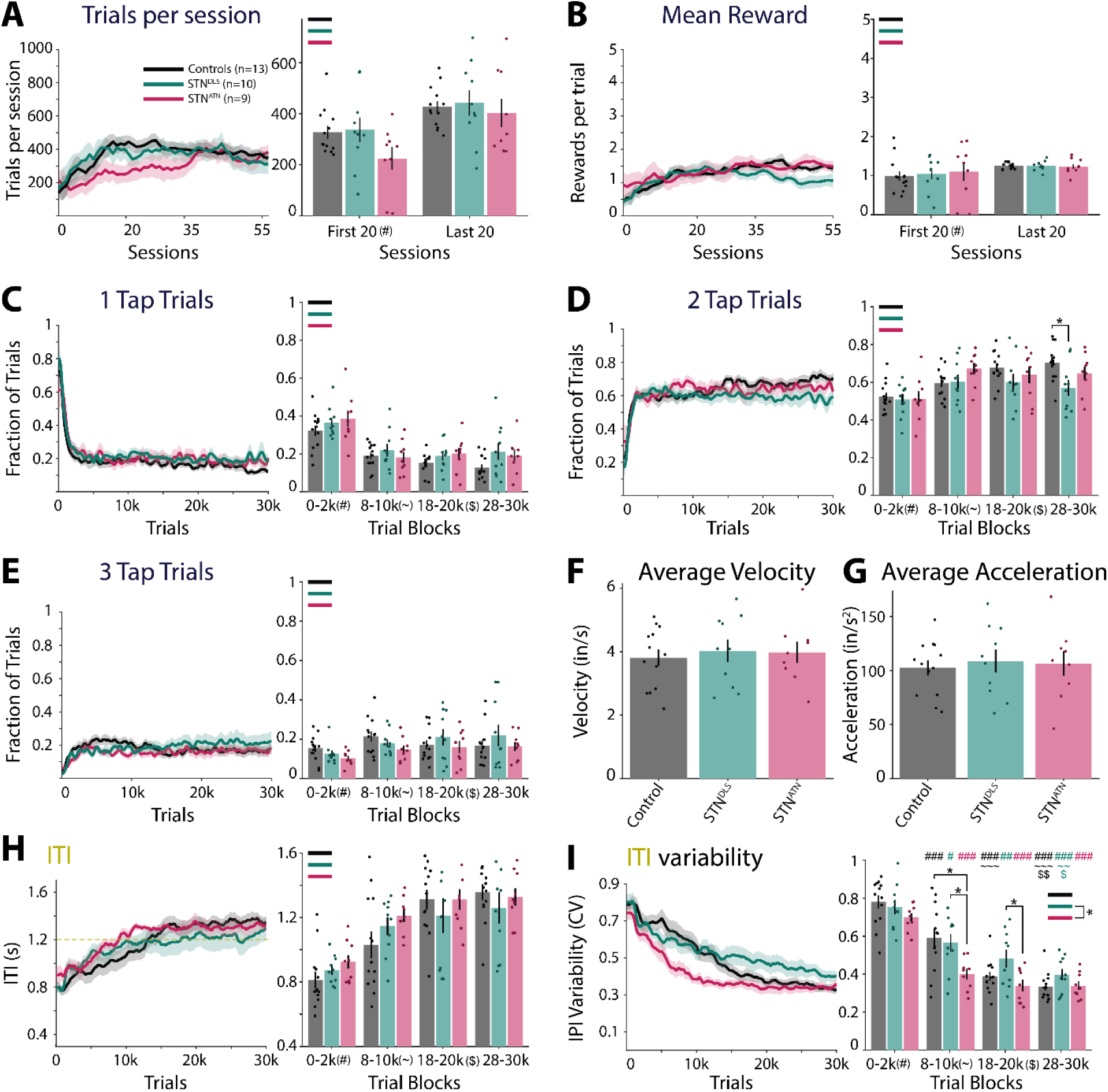
Quantification of additional task parameters after STN^DLS^ and STN^ATN^ neuron silencing. **(A)** Left: Average number of trials performed per training session across learning. Right: Average trials per session for the first and last 20 sessions of training for individual animals. **(B)** As (A), but for the mean reward per trial, averaged across individual training sessions. **(C-E)** Our motor learning task requires rats to press the lever exactly twice (with a specific IPI) to receive a reward – trials with 1, or more than 2 lever presses are not rewarded (see Methods). Shown are the fractions of trials with 1 (C), 2 (D), or 3 or more (E) lever presses across training (left) and in select trial blocks (right). **(F)** Average velocity of task-related hand movements (averaged across hands and vertical and horizontal dimensions) across all trials for each animal. **(G)** As F, but for acceleration of hand movements. **(H)** As (C), but for the mean ITIs. **(I)** As (H), but for the CV of the ITIs. Statistics: (A-E,H-I) 2-way repeated measures ANOVA with post-hoc tests (Holm). (F-G) 1-way ANOVA. For all statistical details, see **Supplementary Table 7.** Bars show means, dots show individual animals, and error bars show SEM. Stars and symbols indicate significance as in Fig. 2.

**Extended Data Figure 4:**
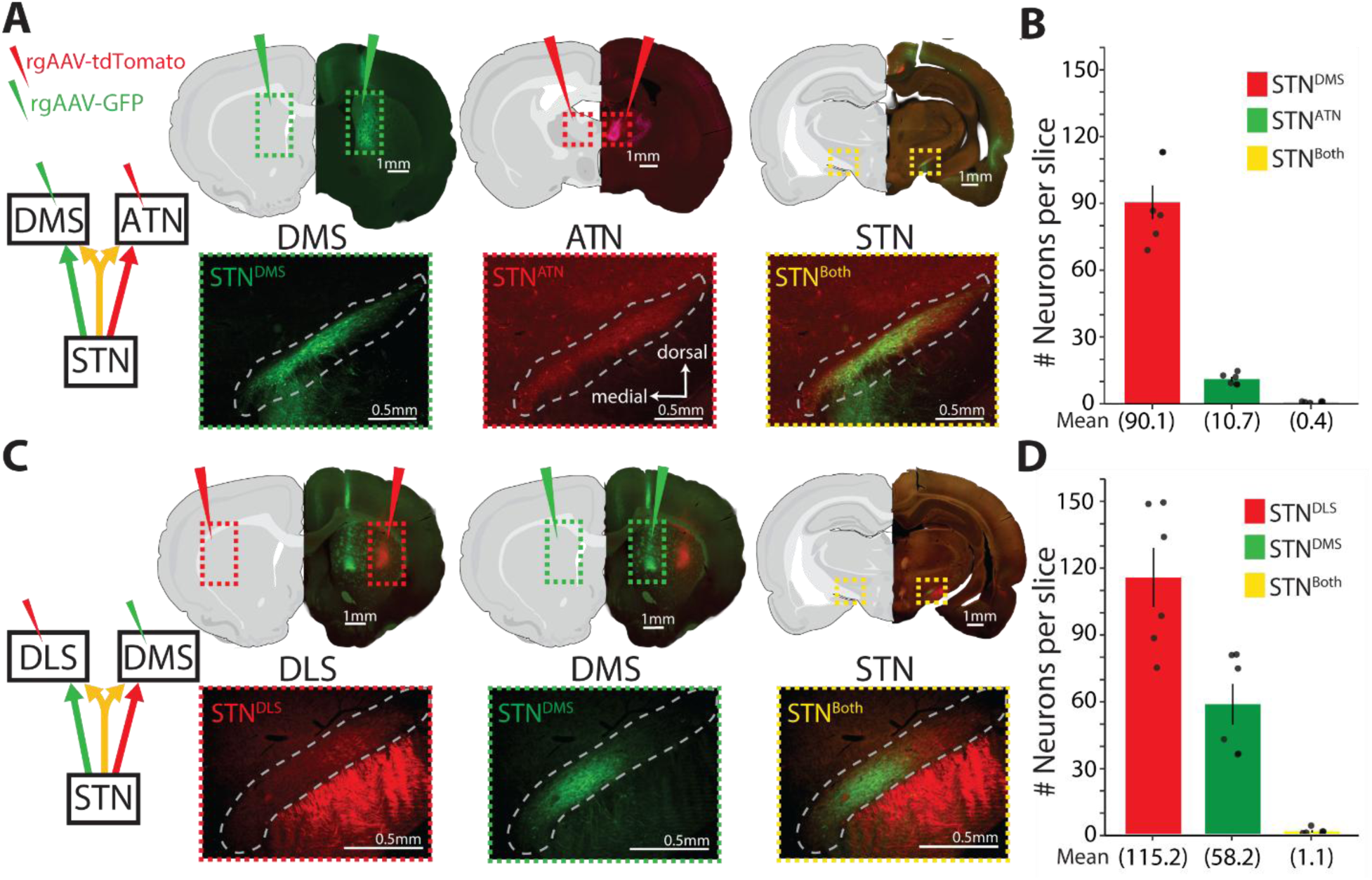
Dual viral retrograde tracing shows no overlap between STN^DMS^ neurons and either STN^DLS^ or STN^ATN^ neurons. **(A)** Dual retrograde viral tracing of STN^DMS^ and STN^ATN^ neurons. Top: Representative images of viral injection sites in the DMS and ATN and of retrograde labeling of STN neurons. Bottom: Representative images of retrogradely labeled STN neurons. **(B)** Quantification of identified STN^DMS^, STN^ATN^, and STN^Both^ (STN neurons that project to both DMS and ATN) neurons per analyzed brain slice (n=5 hemispheres; 3 animals, counts averaged across 3 slices per hemisphere). **(C-D)** As (A-B), but for dual tracing between STN^DMS^ and STN^DLS^ neurons (n=6 hemispheres; 3 animals, counts averaged across 3 slices per hemisphere).

**Extended Data Figure 5:**
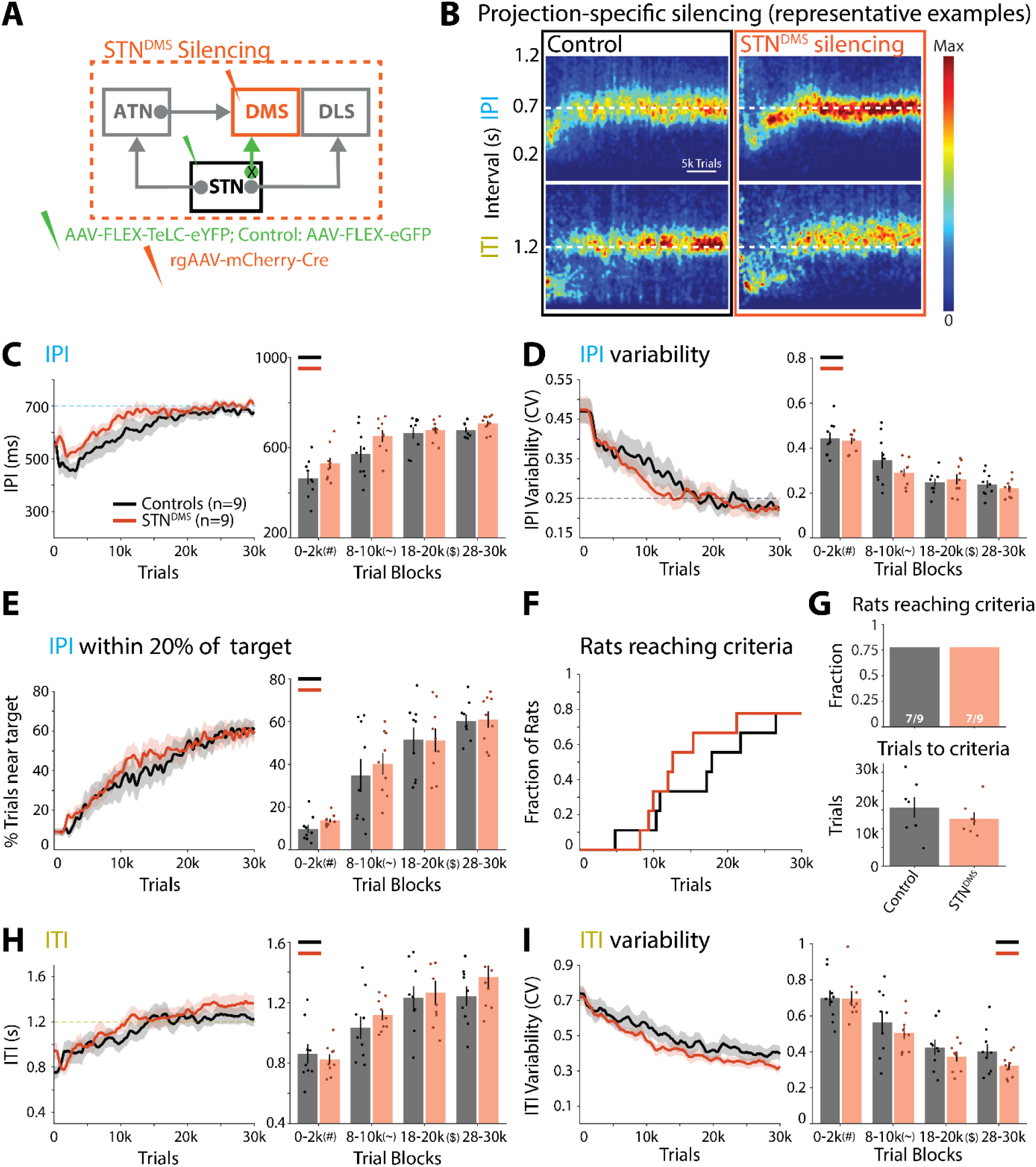
Silencing STN^DMS^ neurons does not impact motor skill learning. **(A)** Left: Viral approach to chronically silence projection-defined STN^DMS^ neurons. See text for details and **Extended Data** Fig. 6A**-C** for injection sites. Animals in the control group express GFP in STN^DLS^ or STN^DMS^ neurons (see Methods and **Extended Data** Fig. 2 for comparison of control cohorts). **(B)** Representative example of task performance throughout training as IPI (top) and ITI (bottom) probability density plots. **(C)** Left: mean IPI across training. Right: Quantification of mean IPI in select trial blocks. **(D)** As (C), but for CV of the IPIs. **(E)** As (C), but for fraction of trials close to the target. **(F)** Fraction of rats reaching learning criteria (see text) across learning. **(G)** Top: Fraction of rats that reached the learning criteria. Bottom: Average number of trials needed to reach the learning criteria. **(H)** As (A), but for the mean ITIs. **(I)** As (A), but for the CV of the ITIs. Statistics: (C-E; H-I) 2-way repeated measures ANOVA with post-hoc tests (Holm). (G) 1-way ANOVA. For all statistical details, see **Supplementary Table 8.** Bars show means, dots show individual animals, and error bars show SEM. Stars and symbols indicate significance as in Fig. 2.

**Extended Data Figure 6:**
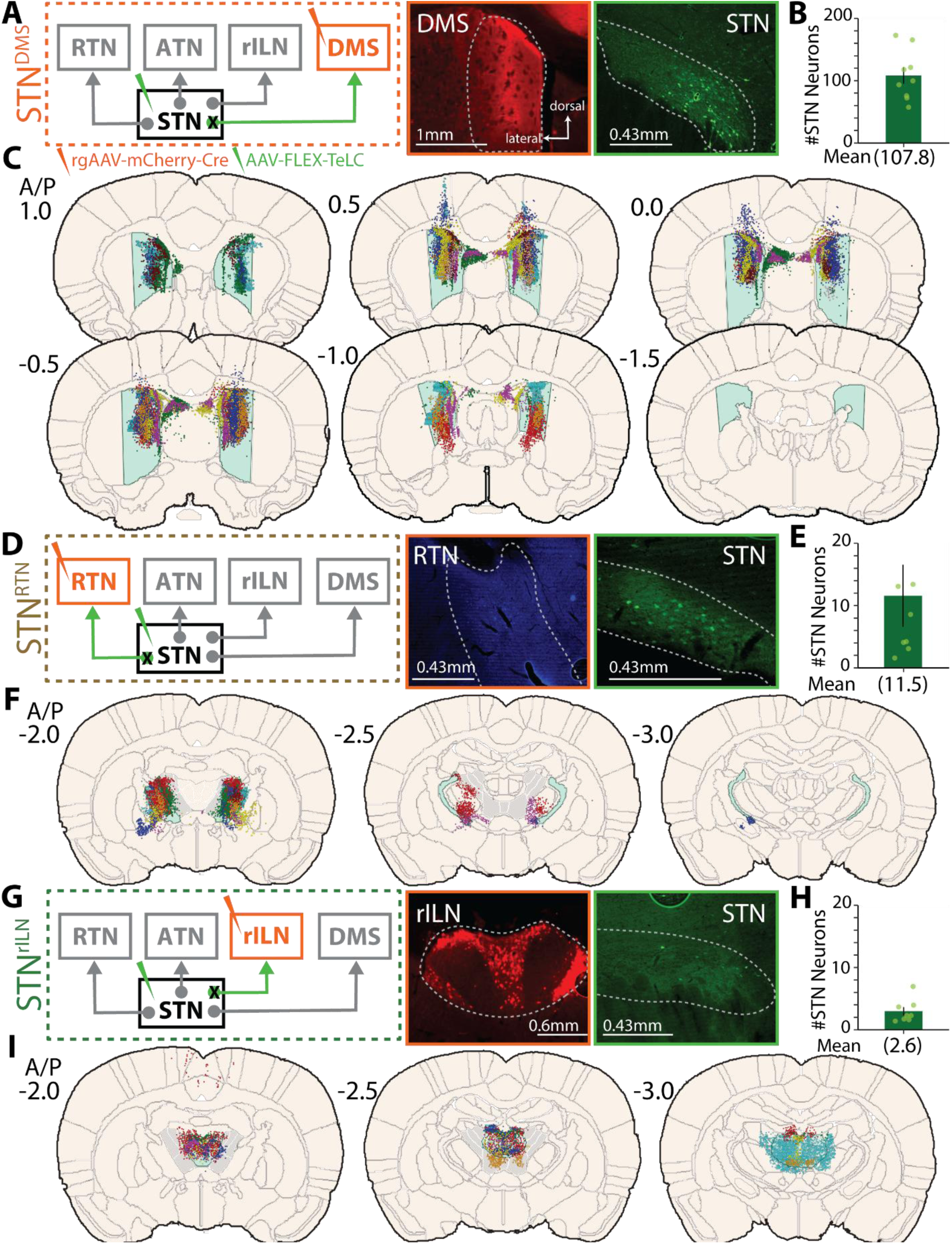
Viral injection sites for silencing of STN neurons projecting to the DMS, or ATN-adjacent thalamic nuclei. **(A)** Left: Circuit scheme for chronic silencing of STN^DMS^ neurons. Right: Representative images of viral injections in the DMS (rgAAV-mCherry-Cre) and in the STN (AAV-FLEX-TeLC-eYFP). Targeted areas are outlined by dotted white lines. **(B)** Quantification of silenced STN^DMS^ neurons per slice (n=9 animals, counts averaged between hemispheres across 3 slices per animal). **(C)** Overlay of viral injection sites in the DMS (light blue) for STN^DMS^ neuron silencing on schematic coronal brain sections (rgAAV-mCherry-Cre). Dots represent automatically detected fluorescent neurons (see Methods) and colors indicate individual animals. **(D-F)** As (A-C), but for STN^RTN^ neuron silencing (n=7 animals, counts averaged between hemispheres across 3 slices per animal). For determining injection sites in the RTN (F), co-injected Fluorogold was used to identify labeled neurons, to avoid inclusion of neurons retrogradely labeled by rgAAV-Cre-mCherry in adjacent thalamic nuclei. **(G-I)** As (A-C), but for STN^rILN^ neuron silencing (n=8 animals, counts averaged between hemispheres across 3 slices per animal).

**Extended Data Figure 7.**
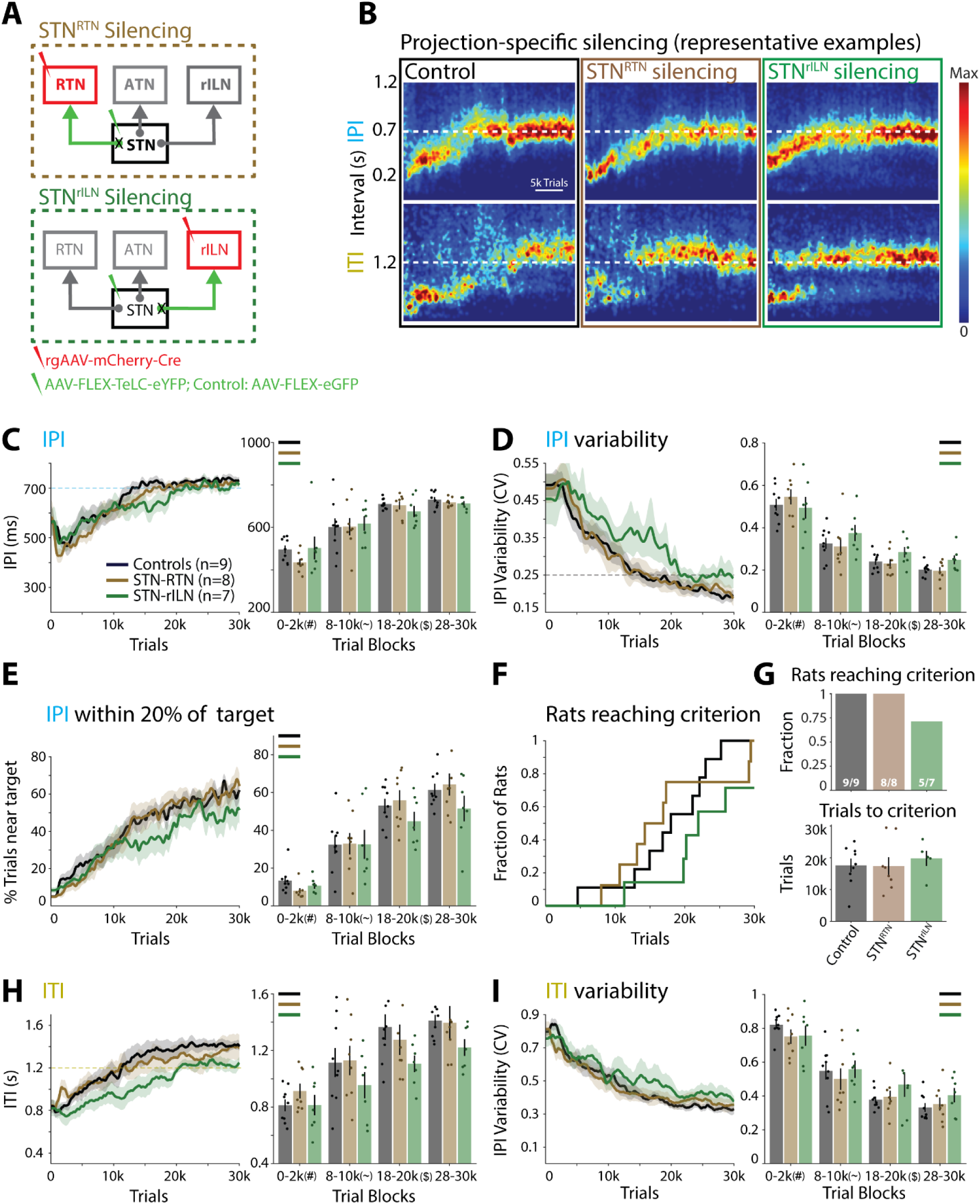
Silencing STN neurons projecting to neighboring thalamic regions does not affect motor skill learning. **(A)** Left: Viral approach to chronically silence STN neurons projecting to the rostral intralaminar thalamic nuclei (STN^rILN^) or to the reticular thalamic nucleus (STN^RTN^). See text for details and **Extended Data** Fig. 6 for injection sites. Animals in the control group express GFP in STN^rILN^, STN^RTN^, or STN^ATN^ neurons (see Methods and **Extended Data** Fig. 2 for comparison of control cohorts). **(B)** Representative example of task performance throughout training as IPI (top) and ITI (bottom) probability density plots. **(C)** Left: mean IPI across training. Right: quantification of mean IPI in select trial blocks. **(D)** As (C), but for CV of the IPIs. **(E)** As (C), but for fraction of trials close to the target. **(F)** Fraction of rats reaching learning criteria (see text) across learning. **(G)** Top: Fraction of rats that reached the learning criteria. Bottom: Average number of trials needed to reach the learning criteria. **(H)** As (A), but for the mean ITIs. **(I)** As (A), but for the CV of the ITIs. Statistics: (C-E; H-I) 2-way repeated measures. (G) 1-way ANOVA. For all statistical details, see **Supplementary Table 9.** Bars show means, dots show individual animals, and error bars show SEM. Stars and symbols indicate significance as in Fig. 2.

**Extended Data Figure 8:**
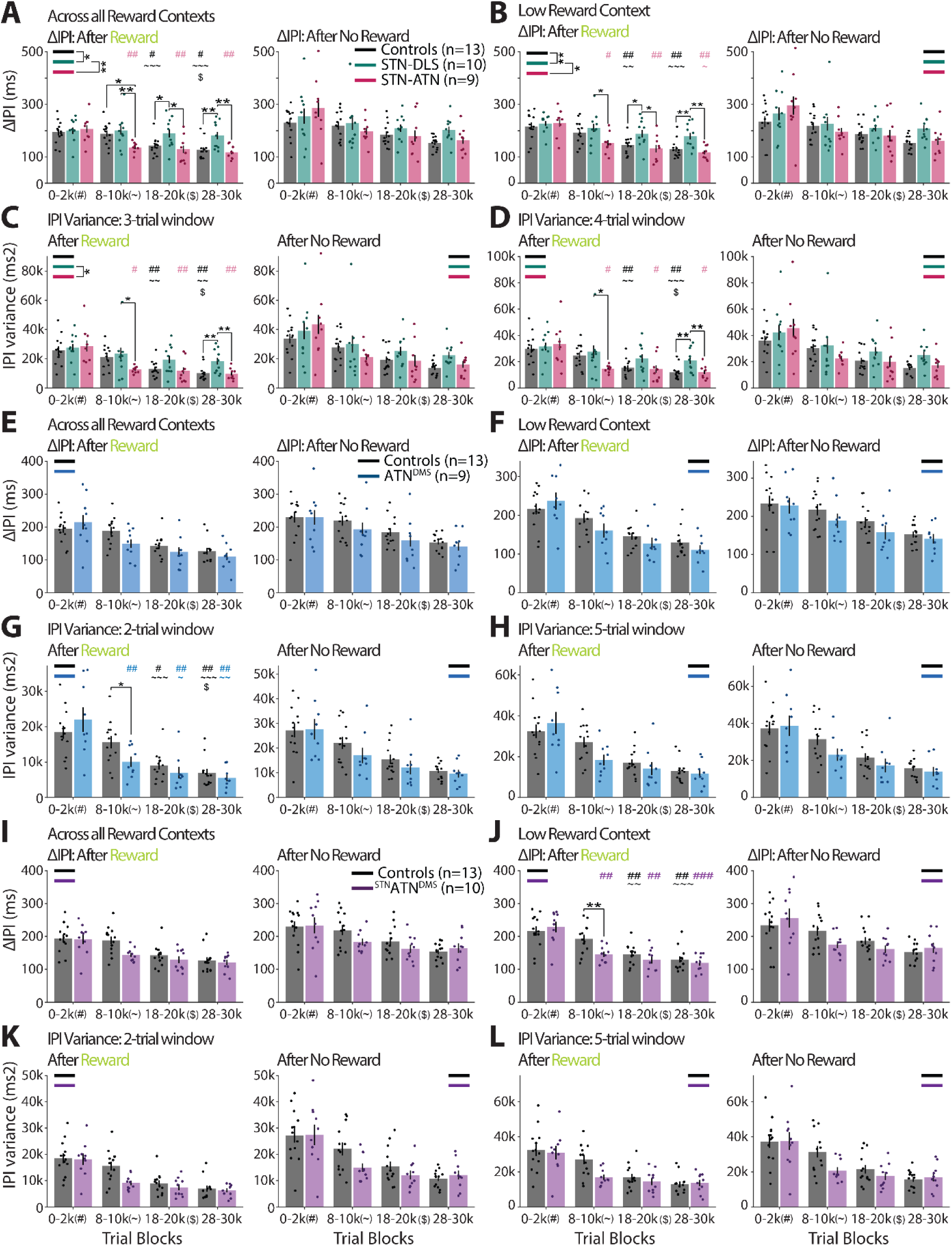
Contributions of STN and ATN projection neurons to variability regulation depend on trial outcome across reward contexts and trial windows. (A-B) To determine whether the trial outcome-dependency of the contributions of STN^DLS^ and STN^ATN^ neurons (Fig. 3) also depends on the reward context, we split all trials into those occurring in a context of high or low reward. (A) Mean ΔIPI for control and STN^DLS^ and STN^ATN^ neuron silencing groups across select trial blocks after rewarded (left) or unrewarded (right) trials across all reward contexts. (B) As (A), but for a low reward context (see Fig. 3B for high reward context). (C) Like (A), but for the variance of the IPI in a 3-trial window after the trigger trial (cf. Fig. 3A, C-D), across all reward contexts. (D) Like (C), but for a 4-trial variance window. (E-F) Like (A-B), but for ATN^DMS^ neuron silencing (see Fig. 5H for high reward context). (G) Like (C), but for ATN^DMS^ neuron silencing in a 2-trial window. (H) Like (G), but for a 5-trial window. (I-L) Like (E-H), but for ^STN^ATN^DMS^ neuron silencing. Statistics: 2-way repeated measures ANOVA with post-hoc tests (Holm). For all statistical details, see Supplementary Table 10. Bars show means, dots show individual animals, and error bars show SEM. Stars and symbols indicate significance as in Fig. 2.

**Extended Data Figure 9:**
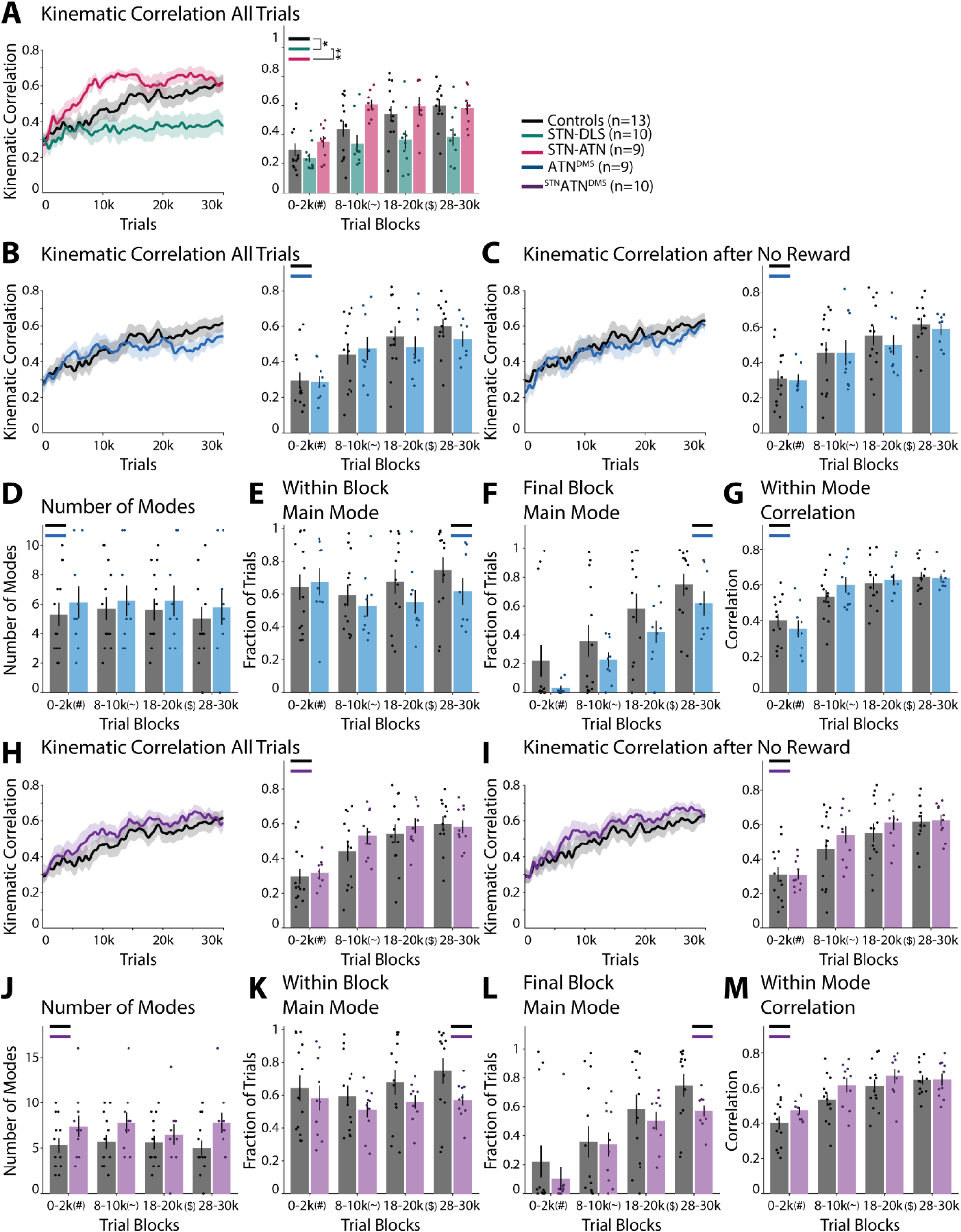
STN or ATN projection neuron silencing contributes to the regulation of kinematic variability. **(A)** Pairwise trial-by-trial correlations between hand trajectories (linearly time-warped, averaged across hands) for all trials, independent of the outcome of the previous trial, for controls and STN^DLS^ and STN^ATN^ neuron silencing groups. Left: Population correlations across learning. Right: Mean correlations in trial blocks. **(B)** As (A), but for ATN^DMS^ neuron silencing. **(C)** As (B), but only for trials after unrewarded trials. **(D)** Number of identified modes, i.e., distinct kinematic solutions to the task (see text for details), in trial blocks for ATN^DMS^ neuron silencing. **(E)** Fraction of trials belonging to the most used (main) mode in trial blocks for ATN^DMS^ neuron silencing. **(F)** Fraction of trials belonging in each trial block to the mode most used at the end of training (main mode in last trial block). **(G)** Weighted average of within-mode kinematic correlations. The average pairwise trial-by-trial kinematic correlations for each mode were averaged for each animal, weighted by mode size in each trial block. **(H-M)** As (B-G), but for ^STN^ATN^DMS^ neuron silencing. Statistics: (A-M) Linear mixed-effects model (LME), random intercept per rat. Post hoc: estimated marginal means from the LME, with Holm correction. For statistical details, see **Supplementary Table 11.** Bars show means, dots show individual animals, and error bars show SEM. Stars and symbols indicate significance as in Fig. 2.

**Extended Data Figure 10:**
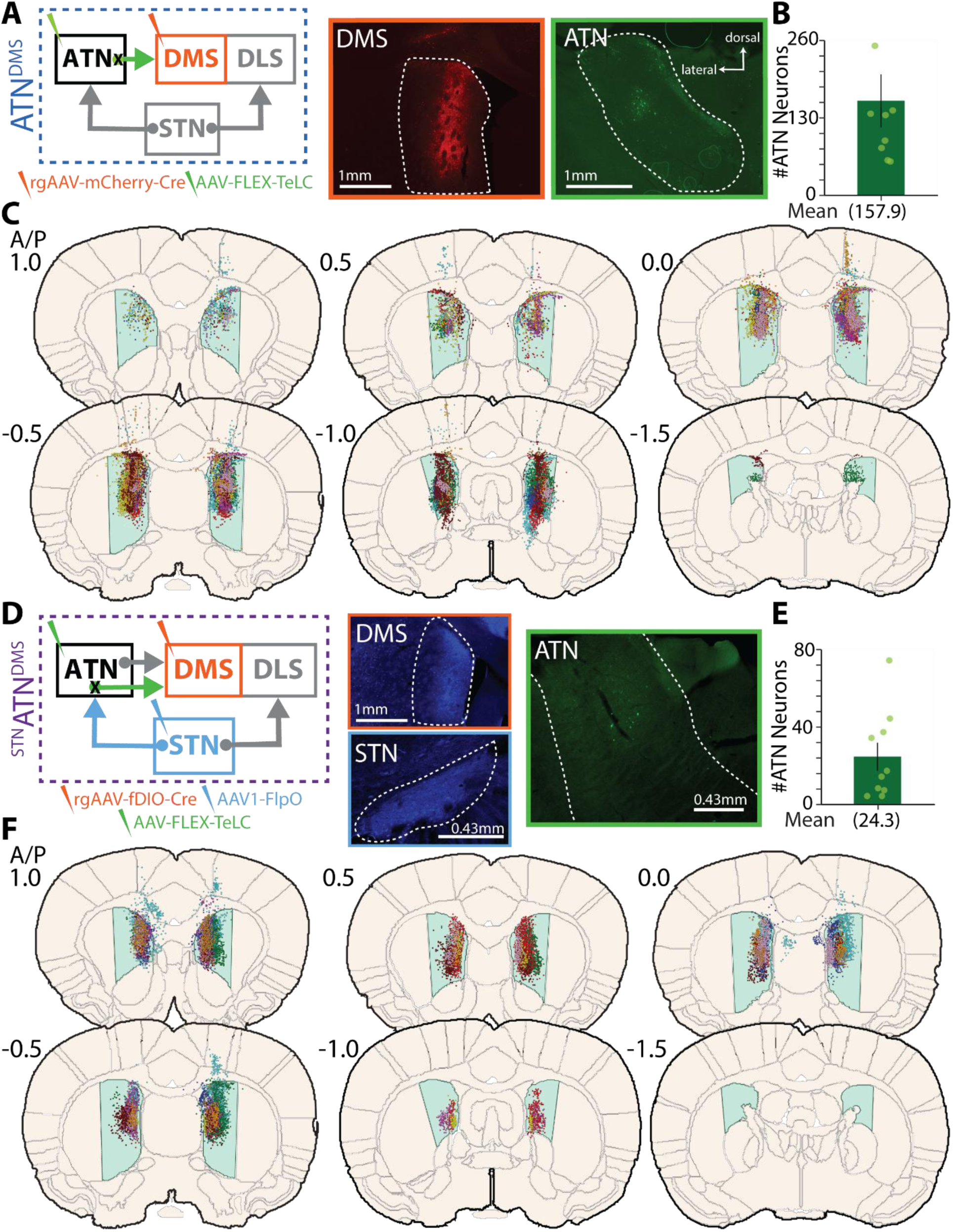
Viral injection sites for silencing of DMS-projecting ATN neurons or STN-targeted, DMS-projecting ATN neurons. **(A)** Left: Circuit scheme for chronic silencing of ATN^DMS^ neurons. Right: Representative images of viral injections in the DMS (rgAAV-mCherry-Cre) and in the ATN (AAV-FLEX-TeLC-eYFP). Targeted areas are outlined by dotted white lines. **(B)** Quantification of silenced ATN^DMS^ neurons per slice (n=9 animals, counts averaged between hemispheres across 3 slices per animal). **(C)** Overlay of viral injection sites in the DMS (light blue) for ATN^DMS^ neuron silencing on schematic coronal brain sections (rgAAV-mCherry-Cre). Dots represent automatically detected fluorescent neurons (see Methods) and colors indicate individual animals. **(D-F)** As (A-C), but for ^STN^ATN^DMS^ neuron silencing (n=10 animals, counts averaged between hemispheres across 3 slices per animal).

## REFERENCES

1. Stein, R. B., Gossen, E. R. & Jones, K. E. Neuronal variability: noise or part of the signal? Nat Rev Neurosci 6, 389–397 (2005).

2. Dhawale, A. K., Smith, M. A. & Ölveczky, B. P. The Role of Variability in Motor Learning. Annual Review of Neuroscience 40, 479–498 (2017).

3. Dhawale, A. K., Miyamoto, Y. R., Smith, M. A. & Ölveczky, B. P. Adaptive Regulation of Motor Variability. Current Biology 29, 3551–3562.e7 (2019).

4. Sternad, D. It’s not (only) the mean that matters: variability, noise and exploration in skill learning. Current Opinion in Behavioral Sciences 20, 183–195 (2018).

5. Therrien, A. S., Wolpert, D. M. & Bastian, A. J. Increasing Motor Noise Impairs Reinforcement Learning in Healthy Individuals. eNeuro 5, (2018).

6. Ali, F. et al. The basal ganglia is necessary for learning spectral, but not temporal, features of birdsong. Neuron 80, 494–506 (2013).

7. Tumer, E. C. & Brainard, M. S. Performance variability enables adaptive plasticity of ‘crystallized’ adult birdsong. Nature 450, 1240–1244 (2007).

8. Wu, H. G., Miyamoto, Y. R., Castro, L. N. G., Ölveczky, B. P. & Smith, M. A. Temporal structure of motor variability is dynamically regulated and predicts motor learning ability. Nat Neurosci 17, 312–321 (2014).

9. Zai, A. T., Lorenz, C., Srikantharajah, S., Giret, N. & Hahnloser, R. H. R. Estimating the motor exploration in reinforcement learning. iScience 29, (2026).

10. Uehara, S., Mawase, F., Therrien, A. S., Cherry-Allen, K. M. & Celnik, P. Interactions between motor exploration and reinforcement learning. J Neurophysiol 122, 797–808 (2019).

11. Bacelar, M. F. B., Lohse, K. R., Parma, J. O. & Miller, M. W. Reinforcement learning in motor skill acquisition: using the reward positivity to understand the mechanisms underlying short-and long-term behavior adaptation. Front. Behav. Neurosci. 18, (2024).

12. Pekny, S. E., Izawa, J. & Shadmehr, R. Reward-dependent modulation of movement variability. Journal of Neuroscience 35, 4015–4024 (2015).

13. Kao, M. H., Doupe, A. J. & Brainard, M. S. Contributions of an avian basal ganglia– forebrain circuit to real-time modulation of song. Nature 433, 638–643 (2005).

14. Kao, M. H., Wright, B. D. & Doupe, A. J. Neurons in a forebrain nucleus required for vocal plasticity rapidly switch between precise firing and variable bursting depending on social context. J Neurosci 28, 13232–13247 (2008).

15. Hessler, N. A. & Doupe, A. J. Singing-related neural activity in a dorsal forebrain-basal ganglia circuit of adult zebra finches. J Neurosci 19, 10461–10481 (1999).

16. Aronov, D., Andalman, A. S. & Fee, M. S. A Specialized Forebrain Circuit for Vocal Babbling in the Juvenile Songbird. Science 320, 630–634 (2008).

17. Ölveczky, B. P., Andalman, A. S. & Fee, M. S. Vocal Experimentation in the Juvenile Songbird Requires a Basal Ganglia Circuit. PLOS Biology 3, e153 (2005).

18. Bottjer, S. W., Miesner, E. A. & Arnold, A. P. Forebrain lesions disrupt development but not maintenance of song in passerine birds. Science 224, 901–903 (1984).

19. Kao, M. H. & Brainard, M. S. Lesions of an Avian Basal Ganglia Circuit Prevent Context-Dependent Changes to Song Variability. Journal of Neurophysiology 96, 1441–1455 (2006).

20. Scharff, C. & Nottebohm, F. A comparative study of the behavioral deficits following lesions of various parts of the zebra finch song system: implications for vocal learning. J Neurosci 11, 2896–2913 (1991).

21. Garst-Orozco, J., Babadi, B. & Ölveczky, B. P. A neural circuit mechanism for regulating vocal variability during song learning in zebra finches. eLife 3, e03697 (2014).

22. Hahnloser, R. H. R., Kozhevnikov, A. A. & Fee, M. S. An ultra-sparse code underlies the generation of neural sequences in a songbird. Nature 419, 65–70 (2002).

23. Long, M. A. & Fee, M. S. Using temperature to analyse temporal dynamics in the songbird motor pathway. Nature 456, 189–194 (2008).

24. Ölveczky, B. P., Otchy, T. M., Goldberg, J. H., Aronov, D. & Fee, M. S. Changes in the neural control of a complex motor sequence during learning. J Neurophysiol 106, 386–397 (2011).

25. Reiner, A., Yamamoto, K. & Karten, H. J. Organization and evolution of the avian forebrain. Anat Rec A Discov Mol Cell Evol Biol 287, 1080–1102 (2005).

26. Dudman, J. T. & Krakauer, J. W. The basal ganglia: from motor commands to the control of vigor. Current Opinion in Neurobiology 37, 158–166 (2016).

27. Park, J., Coddington, L. T. & Dudman, J. T. Basal Ganglia Circuits for Action Specification. Annual Review of Neuroscience 43, 485–507 (2020).

28. Graybiel, A. M. & Grafton, S. T. The Striatum: Where Skills and Habits Meet. Cold Spring Harb Perspect Biol 7, a021691 (2015).

29. Goulet-Kennedy, J., Labbe, S. & Fecteau, S. The involvement of the striatum in decision making. Dialogues Clin Neurosci 18, 55–63 (2016).

30. Rios, A. et al. Dorsomedial striatum monitors unreliability of current action policy and probes alternative one via the indirect pathway. Science Advances 11, eadt4652 (2025).

31. Hiebert, N. M. et al. Striatum in stimulus-response learning via feedback and in decision making. Neuroimage 101, 448–457 (2014).

32. Balleine, B. W., Delgado, M. R. & Hikosaka, O. The Role of the Dorsal Striatum in Reward and Decision-Making. Journal of Neuroscience 27, 8161–8165 (2007).

33. Dhawale, A. K., Wolff, S. B. E., Ko, R. & Ölveczky, B. P. The basal ganglia control the detailed kinematics of learned motor skills. Nat Neurosci 1–14 (2021) doi:10.1038/s41593-021-00889-3.

34. Wolff, S. B. E., Ko, R. & Ölveczky, B. P. Distinct roles for motor cortical and thalamic inputs to striatum during motor skill learning and execution. Science Advances 8, eabk0231 (2022).

35. Turner, K. C., Frost, L., Linsenbardt, D., McIlroy, J. R. & Müller, R.-A. Atypically diffuse functional connectivity between caudate nuclei and cerebral cortex in autism. Behav Brain Funct 2, 34 (2006).

36. Chen, Y. et al. Altered resting-state functional organization within the central executive network in obsessive-compulsive disorder. Psychiatry Clin Neurosci 70, 448–456 (2016).

37. Remijnse, P. L. et al. Reduced orbitofrontal-striatal activity on a reversal learning task in obsessive-compulsive disorder. Arch Gen Psychiatry 63, 1225–1236 (2006).

38. Magrabi, A. et al. Alcohol dependence decreases functional activation of the caudate nucleus during model-based decision processes. Alcoholism: Clinical and Experimental Research 46, 749–758 (2022).

39. Addicott, M. A. et al. Attention-deficit/hyperactivity disorder and the explore/exploit trade-off. Neuropsychopharmacol. 46, 614–621 (2021).

40. Jami, A., Abbaszade, S. & Vahabie, A.-H. A review on exploration–exploitation trade-off in psychiatric disorders. BMC Psychiatry 25, 420 (2025).

41. Van Wouwe, N. C. et al. Deep Brain Stimulation of the Subthalamic Nucleus Improves Reward-Based Decision-Learning in Parkinson’s Disease. Front. Hum. Neurosci. 5, (2011).

42. Macpherson, T. & Hikida, T. Role of basal ganglia neurocircuitry in the pathology of psychiatric disorders. Psychiatry and Clinical Neurosciences 73, 289–301 (2019).

43. Wu, T., Hallett, M. & Chan, P. Motor automaticity in Parkinson’s disease. Neurobiol Dis 82, 226–234 (2015).

44. Puyjarinet, F. et al. Heightened orofacial, manual, and gait variability in Parkinson’s disease results from a general rhythmic impairment. npj Parkinsons Dis. 5, 19 (2019).

45. Prodoehl, J., Corcos, D. M. & Vaillancourt, D. E. Basal Ganglia Mechanisms Underlying Precision Grip Force Control. Neurosci Biobehav Rev 33, 900–908 (2009).

46. Joundi, R. A., Brittain, J.-S., Green, A. L., Aziz, T. Z. & Jenkinson, N. High-frequency stimulation of the subthalamic nucleus selectively decreases central variance of rhythmic finger tapping in Parkinson’s disease. Neuropsychologia 50, 2460–2466 (2012).

47. Pallanti, S. et al. Complex repetitive behavior: Punding after bilateral subthalamic nucleus stimulation in Parkinson’s disease. Parkinsonism & Related Disorders 16, 376–380 (2010).

48. Parolari, L., Schneeberger, M., Heintz, N. & Friedman, J. M. Functional analysis of distinct populations of subthalamic nucleus neurons on Parkinson’s disease and OCD-like behaviors in mice. Mol Psychiatry 26, 7029–7046 (2021).

49. Ardouin, C. et al. Pathological gambling in Parkinson’s disease improves on chronic subthalamic nucleus stimulation. Mov Disord 21, 1941–1946 (2006).

50. Borders, C., Hsu, F., Sweidan, A. J., Matei, E. S. & Bota, R. G. Deep brain stimulation for obsessive compulsive disorder: A review of results by anatomical target. Ment Illn 10, 7900 (2018).

51. Gadot, R. et al. Efficacy of deep brain stimulation for treatment-resistant obsessive-compulsive disorder: systematic review and meta-analysis. J Neurol Neurosurg Psychiatry 93, 1166–1173 (2022).

52. Mallet Luc et al. Subthalamic Nucleus Stimulation in Severe Obsessive– Compulsive Disorder. New England Journal of Medicine 359, 2121–2134 (2008).

53. Eusebio, A. et al. Effects of low-frequency stimulation of the subthalamic nucleus on movement in Parkinson’s disease. Experimental Neurology 209, 125–130 (2008).

54. Bonnevie, T. & Zaghloul, K. A. The Subthalamic Nucleus: Unravelling New Roles and Mechanisms in the Control of Action. Neuroscientist 25, 48–64 (2019).

55. Fife, K. H. et al. Causal role for the subthalamic nucleus in interrupting behavior. eLife Sciences 6, e27689 (2017).

56. Schmidt, R., Leventhal, D. K., Mallet, N., Chen, F. & Berke, J. D. Canceling actions involves a race between basal ganglia pathways. Nat Neurosci 16, 1118–1124 (2013).

57. Tan, S. K. H., Temel, Y., Blokland, A., Steinbusch, H. W. M. & Visser-Vandewalle, V. The subthalamic nucleus: From response selection to execution. Journal of Chemical Neuroanatomy 31, 155–161 (2006).

58. Watson, G. D. R. et al. Thalamic projections to the subthalamic nucleus contribute to movement initiation and rescue of parkinsonian symptoms. Science Advances 7, eabe9192 (2021).

59. Lambert, C. et al. Confirmation of functional zones within the human subthalamic nucleus: Patterns of connectivity and sub-parcellation using diffusion weighted imaging. NeuroImage 60, 83–94 (2012).

60. Bhuvanasundaram, R., Krzyspiak, J. & Khodakhah, K. Subthalamic Nucleus Modulation of the Pontine Nuclei and Its Targeting of the Cerebellar Cortex. J. Neurosci. 42, 5538–5551 (2022).

61. Nakano, K. et al. Topographical projections from the thalamus, subthalamic nucleus and pedunculopontine tegmental nucleus to the striatum in the Japanese monkey, Macaca fuscata. Brain Research 537, 54–68 (1990).

62. Kondabolu, K. et al. A Selective Projection from the Subthalamic Nucleus to Parvalbumin-Expressing Interneurons of the Striatum. eNeuro 10, (2023).

63. Wang, Q. et al. Genoarchitecture and input–output organization of the mouse basal ganglia and thalamic parafascicular nucleus. Nat Neurosci 29, 1248–1264 (2026).

64. Zhang, H. et al. STN–ANT plasticity is crucial for the motor control in Parkinson’s disease model. Sig Transduct Target Ther 6, 1–3 (2021).

65. Hunnicutt, B. J. et al. A comprehensive excitatory input map of the striatum reveals novel functional organization. eLife 5, e19103 (2016).

66. Guo, Q. et al. Whole-Brain Mapping of Inputs to Projection Neurons and Cholinergic Interneurons in the Dorsal Striatum. PLOS ONE 10, e0123381 (2015).

67. Mandelbaum, G. et al. Distinct Cortical-Thalamic-Striatal Circuits through the Parafascicular Nucleus. Neuron 102, 636–652.e7 (2019).

68. Kupferschmidt, D. A., Juczewski, K., Cui, G., Johnson, K. A. & Lovinger, D. M. Parallel, but Dissociable, Processing in Discrete Corticostriatal Inputs Encodes Skill Learning. Neuron 96, 476–489.e5 (2017).

69. Yin, H. H. et al. Dynamic reorganization of striatal circuits during the acquisition and consolidation of a skill. Nat Neurosci 12, 333–341 (2009).

70. Yin, H. H., Ostlund, S. B., Knowlton, B. J. & Balleine, B. W. The role of the dorsomedial striatum in instrumental conditioning. Eur. J. Neurosci. 22, 513–523 (2005).

71. Mizes, K. G. C., Lindsey, J., Escola, G. S. & Ölveczky, B. P. Dissociating the contributions of sensorimotor striatum to automatic and visually guided motor sequences. Nat Neurosci 26, 1791–1804 (2023).

72. Jurado-Parras, M.-T. et al. The Dorsal Striatum Energizes Motor Routines. Current Biology 0, (2020).

73. Hardcastle, K. et al. Differential kinematic coding in sensorimotor striatum across behavioral domains reflects different contributions to movement. Nat Neurosci 28, 1932–1945 (2025).

74. Rueda-Orozco, P. E. & Robbe, D. The striatum multiplexes contextual and kinematic information to constrain motor habits execution. Nat. Neurosci. 18, 453– 460 (2015).

75. Kondapavulur, S. et al. Transition from predictable to variable motor cortex and striatal ensemble patterning during behavioral exploration. Nat Commun 13, 2450 (2022).

76. Kawai, R. et al. Motor cortex is required for learning but not for executing a motor skill. Neuron 86, 800–812 (2015).

77. Murray, A. J. et al. Parvalbumin-positive CA1 interneurons are required for spatial working but not for reference memory. Nature Neuroscience 14, 297–299 (2011).

78. Schiavo, G. G. et al. Tetanus and botulinum-B neurotoxins block neurotransmitter release by proteolytic cleavage of synaptobrevin. Nature 359, 832–835 (1992).

79. Ciriachi, C., Svane-Petersen, D. & Rickhag, M. Genetic tools to study complexity of striatal function. Journal of Neuroscience Research 97, 1181–1193 (2019).

80. Poddar, R., Kawai, R. & Ölveczky, B. P. A fully automated high-throughput training system for rodents. PLoS One 8, e83171 (2013).

81. Redgrave, P. et al. Goal-directed and habitual control in the basal ganglia: implications for Parkinson’s disease. Nat. Rev. Neurosci. 11, 760–772 (2010).

82. Alkemade, A. et al. The functional microscopic neuroanatomy of the human subthalamic nucleus. Brain Struct Funct 224, 3213–3227 (2019).

83. Hassani, O.-K., François, C., Yelnik, J. & Féger, J. Evidence for a dopaminergic innervation of the subthalamic nucleus in the rat. Brain Research 749, 88–94 (1997).

84. Lee, K., Lee, M., Jiang, H. & Kim, H. F. Differential value processing in the rostral and caudal subthalamic nucleus during flexible and habitual decision-making in primates: Implications for cognitive dysfunction in Parkinson’s disease. 2026.10.19.760032 Preprint at 10.64898/2026.10.19.760032 (2026).

85. Collins, J., Zhou, G. Z., Reading, M., Cornish, J. L. & Prasad, A. A. Subthalamic Nucleus dynamically encodes reward value and cognitive flexibility. 2026.10.25.761735 Preprint at 10.64898/2026.10.25.761735 (2026).

86. Härmson, O. et al. Multi-level encoding of reward, effort, and choice across the frontal cortex and basal ganglia during cost-benefit decision-making. Cell Reports 44, 115209 (2025).

87. Darbaky, Y., Baunez, C., Arecchi, P., Legallet, E. & Apicella, P. Reward-related neuronal activity in the subthalamic nucleus of the monkey. Neuroreport 16, 1241– 1244 (2005).

88. Rosenbaum, P. R. & Rubin, D. B. The central role of the propensity score in observational studies for causal effects. Biometrika 70, 41–55 (1983).

89. Pereira, T. D. et al. SLEAP: A deep learning system for multi-animal pose tracking. Nat Methods 19, 486–495 (2022).

90. Aggleton, J. P. & Nelson, A. J. D. Why do lesions in the rodent anterior thalamic nuclei cause such severe spatial deficits? Neurosci Biobehav Rev 54, 131–144 (2015).

91. Jankowski, M. M. et al. The anterior thalamus provides a subcortical circuit supporting memory and spatial navigation. Front Syst Neurosci 7, 45 (2013).

92. Wolff, M., Alcaraz, F., Marchand, A. R. & Coutureau, E. Functional heterogeneity of the limbic thalamus: From hippocampal to cortical functions. Neurosci Biobehav Rev 54, 120–130 (2015).

93. Zingg, B. et al. AAV-Mediated Anterograde Transsynaptic Tagging: Mapping Corticocollicular Input-Defined Neural Pathways for Defense Behaviors. Neuron 93, 33–47 (2017).

94. Zingg, B., Peng, B., Huang, J., Tao, H. W. & Zhang, L. I. Synaptic Specificity and Application of Anterograde Transsynaptic AAV for Probing Neural Circuitry. J. Neurosci. 40, 3250–3267 (2020).

95. Zingg, B., Dong, H.-W., Tao, H. W. & Zhang, L. I. Application of AAV1 for Anterograde Transsynaptic Circuit Mapping and Input-Dependent Neuronal Cataloging. Current Protocols 2, e339 (2022).

96. Turner, K. M., Svegborn, A., Langguth, M., McKenzie, C. & Robbins, T. W. Opposing Roles of the Dorsolateral and Dorsomedial Striatum in the Acquisition of Skilled Action Sequencing in Rats. J. Neurosci. 42, 2039–2051 (2022).

97. Díaz-Hernández, E. et al. The Thalamostriatal Projections Contribute to the Initiation and Execution of a Sequence of Movements. Neuron 100, 739–752.e5 (2018).

98. Balleine, B. W. & Dezfouli, A. Hierarchical Action Control: Adaptive Collaboration Between Actions and Habits. Frontiers in Psychology 10, (2019).

99. Bergstrom, H. C. et al. Dorsolateral Striatum Engagement Interferes with Early Discrimination Learning. Cell Reports 23, 2264–2272 (2018).

100. Smith, K. S. & Graybiel, A. M. Habit formation coincides with shifts in reinforcement representations in the sensorimotor striatum. Journal of Neurophysiology 115, 1487–1498 (2016).

101. Balleine, B. W., Liljeholm, M. & Ostlund, S. B. The integrative function of the basal ganglia in instrumental conditioning. Behavioural Brain Research 199, 43–52 (2009).

102. Jeon, H. et al. Topographic connectivity and cellular profiling reveal detailed input pathways and functionally distinct cell types in the subthalamic nucleus. Cell Reports 38, 110439 (2022).

103. Emmi, A., Antonini, A., Macchi, V., Porzionato, A. & De Caro, R. Anatomy and Connectivity of the Subthalamic Nucleus in Humans and Non-human Primates. Frontiers in Neuroanatomy 14, (2020).

104. Kita, T. & Kita, H. The Subthalamic Nucleus Is One of Multiple Innervation Sites for Long-Range Corticofugal Axons: A Single-Axon Tracing Study in the Rat. J. Neurosci. 32, 5990–5999 (2012).

105. Nambu, A., Tokuno, H. & Takada, M. Functional significance of the cortico-subthalamo-pallidal ‘hyperdirect’ pathway. Neurosci. Res. 43, 111–117 (2002).

106. Gittis, A. H., Nelson, A. B., Thwin, M. T., Palop, J. J. & Kreitzer, A. C. Distinct Roles of GABAergic Interneurons in the Regulation of Striatal Output Pathways. J. Neurosci. 30, 2223–2234 (2010).

107. Koós, T. & Tepper, J. M. Inhibitory control of neostriatal projection neurons by GABAergic interneurons. Nat Neurosci 2, 467–472 (1999).

108. Gritton, H. J. et al. Unique contributions of parvalbumin and cholinergic interneurons in organizing striatal networks during movement. Nat Neurosci 22, 586–597 (2019).

109. Owen, S. F., Berke, J. D. & Kreitzer, A. C. Fast-Spiking Interneurons Supply Feedforward Control of Bursting, Calcium, and Plasticity for Efficient Learning. Cell 172, 683–695.e15 (2018).

110. Klug, J. R. et al. Differential inputs to striatal cholinergic and parvalbumin interneurons imply functional distinctions. eLife 7, e35657 (2018).

111. Choi, K., Holly, E. N., Davatolhagh, M. F., Beier, K. T. & Fuccillo, M. V. Integrated anatomical and physiological mapping of striatal afferent projections. European Journal of Neuroscience 49, 623–636 (2019).

112. Jung, F., Cao, X., Heymans, L. & Carlén, M. DMC-BrainMap is an open-source, end-to-end tool for multi-feature brain mapping in different species. Cell Rep Methods 6, 101302 (2026).

113. Chen, J. et al. The Allen Cell and Structure Segmenter: a new open source toolkit for segmenting 3D intracellular structures in fluorescence microscopy images. 491035 Preprint at 10.1101/491035 (2020).

114. McInnes, L., Healy, J. & Melville, J. UMAP: Uniform Manifold Approximation and Projection for Dimension Reduction. Preprint at 10.48550/arXiv.1802.03426 (2020).

115. Langfelder, P., Zhang, B. & Horvath, S. Defining clusters from a hierarchical cluster tree: the Dynamic Tree Cut package for R. Bioinformatics 24, 719–720 (2008).

