## Supplementary Tables for "Non-canonical Subthalamic Nucleus projections regulate motor variability"

**Supplementary Table 1. Statistical Results for Figure 2.**

**Figure 2E** *Mean IPI over the course of training. 2-Way RM ANOVA, Post hoc: Holm*
Group: F(2,29) = 8.042, p = 0.002. Block: F(3,87) = 81.388, p<0.001
Group x Block: F(6,87) = 3.708, p = 0.002

| Between Groups | | | | | | Within Group | | | | |
|---|---|---|---|---|---|---|---|---|---|---|
| | vs. | 0-2k | 8-10k | 18-20k | 28-30k | | vs. | Cont. | STN$^{DLS}$ | STN$^{ATN}$ |
| Control | STN$^{DLS}$ | 0.263 | 0.536 | 0.060 | 0.493 | 0-2k (#) | 8-10k | <.001 | 0.108 | 0.001 |
| | STN$^{ATN}$ | 0.091 | 0.001 | 0.093 | 0.493 | | 18-20k | <.001 | 0.048 | <.001 |
| STN$^{DLS}$ | STN$^{ATN}$ | 0.473 | 0.006 | 0.003 | 0.240 | | 28-30k | <.001 | 0.011 | <.001 |
| | | All | | | | 8-10k (~) | 18-20k | <.001 | 0.108 | 1.000 |
| Control | STN$^{DLS}$ | 0.827 | | | | | 28-30k | 0.002 | 0.087 | 1.000 |
| | STN$^{ATN}$ | 0.002 | | | | 18-20k ($) | 28-30k | 0.649 | 0.108 | 0.387 |
| STN$^{DLS}$ | STN$^{ATN}$ | 0.005 | | | | | | | | |

**Figure 2F** *IPI Variability over the course of training. 2-Way RM ANOVA, Post hoc: Holm*
Group: F(2,29) = 5.068, p = 0.013. Block: F(3,87) = 140.498, p<0.001.
Group x Block: F(6,87) = 5.098, p<0.001

| Between Groups (*) | | | | | | Within Group | | | | |
|---|---|---|---|---|---|---|---|---|---|---|
| | vs. | 0-2k | 8-10k | 18-20k | 28-30k | | vs. | Cont. | STN$^{DLS}$ | STN$^{ATN}$ |
| Control | STN$^{DLS}$ | 0.461 | 0.761 | 0.034 | 0.004 | 0-2k (#) | 8-10k | <.001 | 0.033 | <.001 |
| | STN$^{ATN}$ | 0.276 | 0.002 | 0.423 | 0.907 | | 18-20k | <.001 | 0.004 | <.001 |
| STN$^{DLS}$ | STN$^{ATN}$ | 0.605 | 0.006 | 0.013 | 0.008 | | 28-30k | <.001 | <.001 | <.001 |
| | | All | | | | 8-10k (~) | 18-20k | <.001 | 0.033 | 0.912 |
| Control | STN$^{DLS}$ | 0.445 | | | | | 28-30k | <.001 | 0.033 | 0.193 |
| | STN$^{ATN}$ | 0.039 | | | | 18-20k ($) | 28-30k | <.001 | 0.033 | 0.912 |
| STN$^{DLS}$ | STN$^{ATN}$ | 0.015 | | | | | | | | |

**Figure 2G** *Fraction of trials near target. 2-Way RM ANOVA, Post hoc: Holm*
Group: F(2,29) = 5.167, p = 0.012. Block: F(3,87) = 158.026, p<0.001.
Group x Block: F(6,87) = 6.175, p<0.001

| Between Groups | | | | | | Within Group | | | | |
|---|---|---|---|---|---|---|---|---|---|---|
| | vs. | 0-2k | 8-10k | 18-20k | 28-30k | | vs. | Cont. | STN$^{DLS}$ | STN$^{ATN}$ |
| Control | STN$^{DLS}$ | 0.899 | 0.824 | 0.194 | 0.004 | 0-2k (#) | 8-10k | <.001 | 0.006 | <.001 |
| | STN$^{ATN}$ | 0.374 | 0.003 | 0.635 | 0.588 | | 18-20k | <.001 | <.001 | <.001 |
| STN$^{DLS}$ | STN$^{ATN}$ | 0.374 | 0.003 | 0.157 | 0.022 | | 28-30k | <.001 | <.001 | <.001 |
| | | All | | | | 8-10k (~) | 18-20k | <.001 | 0.006 | 0.918 |
| Control | STN$^{DLS}$ | 0.173 | | | | | 28-30k | <.001 | 0.006 | 0.770 |
| | STN$^{ATN}$ | 0.173 | | | | 18-20k ($) | 28-30k | 0.003 | 0.356 | 0.770 |
| STN$^{DLS}$ | STN$^{ATN}$ | 0.010 | | | | | | | | |

**Figure 2I** *Number of trials needed to reach learning criterion. 1-Way ANOVA Post hoc: Holm*
Group: F(2,23) = 5.833, p=0.009

| Between Groups | | |
|---|---|---|
| Control | STN$^{DLS}$ | 0.543 |
| | STN$^{ATN}$ | 0.008 |
| STN$^{DLS}$ | STN$^{ATN}$ | 0.100 |

**Supplementary Table 2. Statistical Results for Figure 3.**

**Figure 3B** (Left): ΔIPI after reward (high reward context). *2-Way RM ANOVA, Post hoc: Holm*
Group: F(2,29) = 8.381, p = 0.001. Block: F(3,87) = 5.340, p = 0.002
Group x Block: F(6,87) = 4.420, p<0.001

| Between Groups | | | | | | Within Group | | | | |
|---|---|---|---|---|---|---|---|---|---|---|
| | vs. | 0-2k | 8-10k | 18-20k | 28-30k | | vs. | Cont. | STN$^{DLS}$ | STN$^{ATN}$ |
| Controls | STN$^{DLS}$ | 1.000 | 0.818 | 0.020 | <.001 | 0-2k | 8-10k | 0.436 | 1.000 | 0.020 |
| | STN$^{ATN}$ | 1.000 | 0.016 | 0.623 | 0.565 | | 18-20k | 0.181 | 1.000 | 0.062 |
| STN$^{DLS}$ | STN$^{ATN}$ | 1.000 | 0.016 | 0.016 | <.001 | | 28-30k | 0.114 | 1.000 | 0.008 |
| | | All | | | | 8-10k | 18-20k | 0.006 | 1.000 | 0.945 |
| Controls | STN$^{DLS}$ | 0.167 | | | | | 28-30k | 0.006 | 1.000 | 0.307 |
| | STN$^{ATN}$ | 0.001 | | | | 18-20k | 28-30k | 0.198 | 1.000 | 0.559 |
| STN$^{DLS}$ | STN$^{ATN}$ | 0.015 | | | | | | | | |

**Figure 3B** (Right): ΔIPI after no reward (high reward context). *2-Way RM ANOVA*
Group: F(2,29) = 0.471, p = 0.629. Block: F(3,87) = 12.545, p<0.001
Group x Block: F(6,87) = 0.996, p = 0.433

**Figure 3C** (Left): IPI variance (2-trial window) after reward *2-Way RM ANOVA, Post hoc: Holm*
Group: F(2,29) = 4.723, p = 0.017. Block: F(3,87) = 35.064, p<0.001
Group x Block: F(6,87) = 3.163, p = 0.007

| Between Groups | | | | | | Within Group | | | | |
|---|---|---|---|---|---|---|---|---|---|---|
| | vs. | 0-2k | 8-10k | 18-20k | 28-30k | | vs. | Cont. | STN$^{DLS}$ | STN$^{ATN}$ |
| Controls | STN$^{DLS}$ | 1.000 | 0.563 | 0.024 | 0.003 | 0-2k | 8-10k | 0.224 | 0.530 | 0.004 |
| | STN$^{ATN}$ | 1.000 | 0.029 | 0.548 | 0.684 | | 18-20k | 0.002 | 0.483 | 0.003 |
| STN$^{DLS}$ | STN$^{ATN}$ | 1.000 | 0.017 | 0.015 | 0.003 | | 28-30k | 0.001 | 0.253 | 0.003 |
| | | All | | | | 8-10k | 18-20k | <.001 | 0.530 | 0.652 |
| Controls | STN$^{DLS}$ | 0.077 | | | | | 28-30k | <.001 | 0.512 | 0.090 |
| | STN$^{ATN}$ | 0.296 | | | | 18-20k | 28-30k | 0.014 | 0.530 | 0.464 |
| STN$^{DLS}$ | STN$^{ATN}$ | 0.017 | | | | | | | | |

**Figure 3C** (Right): IPI variance (2-trial window) after no reward. *2-Way RM ANOVA*
Group: F(2,29) = 1.009, p = 0.377. Block: F(3,87) = 26.795, p<0.001
Group x Block: F(6,87) = 1.397, p = 0.225

**Figure 3D** (Left): IPI variance (5-trial window) after reward. *2-Way RM ANOVA, Post hoc: Holm*
Group: $F(2,29) = 2.963$, $p = 0.067$. Block: $F(3,87) = 29.783$, $p<0.001$
Group x Block: $F(6,87) = 2.384$, $p = 0.035$

| Between Groups | | | | | | Within Group | | | | |
|---|---|---|---|---|---|---|---|---|---|---|
| | vs. | 0-2k | 8-10k | 18-20k | 28-30k | | vs. | Cont. | $STN^{DLS}$ | $STN^{ATN}$ |
| Controls | $STN^{DLS}$ | 1.000 | 0.579 | 0.115 | 0.003 | 0-2k | 8-10k | 0.153 | 0.580 | 0.014 |
| | $STN^{ATN}$ | 1.000 | 0.063 | 0.783 | 0.913 | | 18-20k | 0.004 | 0.332 | 0.011 |
| $STN^{DLS}$ | $STN^{ATN}$ | 1.000 | 0.039 | 0.115 | 0.006 | | 28-30k | 0.002 | 0.167 | 0.011 |
| | | | | | | 8-10k | 18-20k | 0.003 | 0.580 | 0.905 |
| | | | | | | | 28-30k | <.001 | 0.570 | 0.558 |
| | | | | | | 18-20k | 28-30k | 0.039 | 0.580 | 0.558 |

**Figure 3D** (Right)**:** IPI variance (5-trial window) after no reward*. 2-Way RM ANOVA, Post hoc: Holm*
Group: $F(2,29) = 1.715$, $p = 0.198$. Block: $F(3,87) = 29.210$, $p<0.001$
Group x Block: $F(6,87) = 1.697$, $p = 0.131$

**Supplementary Table 3. Statistical Results for Figure 4.**

**Figure 4B** *Kinematic correlation after rewarded trials. Linear mixed-effects model (LME), random intercept per rat. Post hoc: Estimated marginal means from the LME, with holm correction.*
Group: $F(2,29) = 5.534$, $p = 0.009$. Block: $F(3,83.5) = 25.549$, $p<0.001$
Group x Block: $F(6,83.5) = 2.210$, $p = 0.050$

| Between Groups | | | | | | Within Group | | | | |
|---|---|---|---|---|---|---|---|---|---|---|
| | vs. | 0-2k | 8-10k | 18-20k | 28-30k | | vs. | Cont. | STN$^{DLS}$ | STN$^{ATN}$ |
| Controls | STN$^{DLS}$ | 0.604 | 0.273 | 0.109 | 0.025 | 0-2k | 8-10k | 0.02 | 0.424 | 0.012 |
| | STN$^{ATN}$ | 0.604 | 0.018 | 0.384 | 0.922 | | 18-20k | 0.005 | 0.362 | 0.014 |
| STN$^{DLS}$ | STN$^{ATN}$ | 0.411 | 0.004 | 0.042 | 0.033 | | 28-30k | <.001 | 0.362 | 0.014 |
| | | All | | | | 8-10k | 18-20k | 0.064 | 0.424 | 0.997 |
| Controls | STN$^{DLS}$ | 0.079 | | | | | 28-30k | 0.052 | 0.4 | 0.928 |
| | STN$^{ATN}$ | 0.174 | | | | 18-20k | 28-30k | 0.496 | 0.761 | 0.928 |
| STN$^{DLS}$ | STN$^{ATN}$ | 0.008 | | | | | | | | |

**Figure 4C** *Kinematic correlation after unrewarded trials. Linear mixed-effects model (LME), random intercept per rat. Post hoc: Estimated marginal means from the LME, with holm correction.*
Group: $F(2,29.1471) = 6.130$, $p = 0.006$. Block: $F(3,83.56) = 29.033$, $p<0.001$
Group x Block: $F(6,83.5) = 1.544$, $p = 0.174$

| Between Groups | | |
|---|---|---|
| | | All |
| Controls | STN$^{DLS}$ | 0.024 |
| | STN$^{ATN}$ | 0.368 |
| STN$^{DLS}$ | STN$^{ATN}$ | 0.007 |

**Figure 4D** *Number of Modes. Linear mixed-effects model (LME), random intercept per rat. Post hoc: Estimated marginal means from the LME, with holm correction.*
Group: $F(2,29) = 5.056$, $p = 0.013$. Block: $F(3,87) = 1.187$, $p = 0.320$
Group x Block: $F(6,87) = 1.087$, $p = 0.376$

| Between Groups | | |
|---|---|---|
| | | All |
| Controls | STN$^{DLS}$ | 0.025 |
| | STN$^{ATN}$ | 0.916 |
| STN$^{DLS}$ | STN$^{ATN}$ | 0.025 |

**Figure 4E** *Within block main mode. Linear mixed-effects model (LME), random intercept per rat. Post hoc: Estimated marginal means from the LME, with holm correction.*
Group: $F(2,29.08) = 4.293$, $p = 0.023$. Block: $F(3,83.47) = 1.443$, $p = 0.236$
Group x Block: $F(6,83.45) = 1.323$, $p = 0.256$

| Between Groups | | |
|---|---|---|
| | | All |
| Controls | STN$^{DLS}$ | 0.048 |
| | STN$^{ATN}$ | 0.809 |
| STN$^{DLS}$ | STN$^{ATN}$ | 0.048 |

**Figure 4F** *Final block main mode. Linear mixed-effects model (LME), random intercept per rat. Post hoc: Estimated marginal means from the LME, with holm correction.*
Group: F(2,28.88) = 3.699, p = 0.037. Block: F(3,83.22) = 37.121, p<0.001
Group x Block: F(6,83.2) = 2.589, p = 0.024

| Between Groups | | | | | | Within Group | | | | |
|---|---|---|---|---|---|---|---|---|---|---|
| | vs. | 0-2k | 8-10k | 18-20k | 28-30k | | vs. | Cont. | STN$^{DLS}$ | STN$^{ATN}$ |
| Controls | STN$^{DLS}$ | 0.574 | 0.104 | 0.085 | 0.149 | 0-2k | 8-10k | 0.128 | 0.290 | <.001 |
| | STN$^{ATN}$ | 0.751 | 0.046 | 0.562 | 0.761 | | 18-20k | <.001 | 0.007 | <.001 |
| STN$^{DLS}$ | STN$^{ATN}$ | 0.734 | 0.002 | 0.056 | 0.248 | | 28-30k | <.001 | <.001 | <.001 |
| | | All | | | | 8-10k | 18-20k | 0.008 | 0.133 | 1.000 |
| Controls | STN$^{DLS}$ | 0.090 | | | | | 28-30k | <.001 | 0.002 | 1.000 |
| | STN$^{ATN}$ | 0.495 | | | | 18-20k | 28-30k | 0.128 | 0.169 | 1.000 |
| STN$^{DLS}$ | STN$^{ATN}$ | 0.047 | | | | | | | | |

**Figure 4G** *Mode Stereotypy. Linear mixed-effects model (LME), random intercept per rat. Post hoc: Estimated marginal means from the LME, with holm correction.*
Group: F(2,29.3) = 2.306, p = 0.117. Block: F(3,83.88) = 17.029, p<0.001
Group x Block: F(6,83.86) = 2.437, p = 0.032

| Between Groups | | | | | | Within Group | | | | |
|---|---|---|---|---|---|---|---|---|---|---|
| | vs. | 0-2k | 8-10k | 18-20k | 28-30k | | vs. | Cont. | STN$^{DLS}$ | STN$^{ATN}$ |
| Controls | STN$^{DLS}$ | 1 | 0.349 | 0.208 | 0.037 | 0-2k | 8-10k | 0.004 | 1 | <.001 |
| | STN$^{ATN}$ | 1 | 0.144 | 0.754 | 0.528 | | 18-20k | <.001 | 0.639 | 0.002 |
| STN$^{DLS}$ | STN$^{ATN}$ | 1 | 0.039 | 0.208 | 0.161 | | 28-30k | <.001 | 0.852 | 0.003 |
| | | | | | | 8-10k | 18-20k | 0.111 | 1 | 1 |
| | | | | | | | 28-30k | 0.017 | 1 | 1 |
| | | | | | | 18-20k | 28-30k | 0.348 | 1 | 1 |

**Supplementary Table 4. Statistical Results for Figure 5.**

**Figure 5C** *Mean IPI over the course of training. 2-Way RM ANOVA, Post hoc: Holm*
Group: $F(1,20) = 2.045$, $p = 0.168$. Block: $F(3,60) = 110.406$, $p<.001$
Group x Block: $F(3,60) = 3.002$, $p = 0.037$

| Between Groups | | | | | | Within Group | | | |
|---|---|---|---|---|---|---|---|---|---|
| | vs. | 0-2k | 8-10k | 18-20k | 28-30k | | vs. | Control | $ATN^{DMS}$ |
| Control | $ATN^{DMS}$ | 0.667 | 0.035 | 0.720 | 0.947 | 0-2k | 8-10k | <.001 | <.001 |
| | | | | | | | 18-20k | <.001 | <.001 |
| | | | | | | | 28-30k | <.001 | <.001 |
| | | | | | | 8-10k | 18-20k | <.001 | 0.126 |
| | | | | | | | 28-30k | 0.001 | 0.106 |
| | | | | | | 18-20k | 28-30k | 0.649 | 0.934 |

**Figure 5D** *IPI variability over the course of training. 2-Way RM ANOVA, Post hoc: Holm*
Group: $F(1,20) = 2.447$, $p = 0.133$. Block: $F(3,60) = 162.804$, $p<0.001$
Group x Block: $F(3,60) = 4.693$, $p = 0.005$

| Between Groups | | | | | | Within Group | | | |
|---|---|---|---|---|---|---|---|---|---|
| | vs. | 0-2k | 8-10k | 18-20k | 28-30k | | vs. | Control | $ATN^{DMS}$ |
| Control | $ATN^{DMS}$ | 0.965 | 0.007 | 0.322 | 0.510 | 0-2k | 8-10k | <.001 | <.001 |
| | | | | | | | 18-20k | <.001 | <.001 |
| | | | | | | | 28-30k | <.001 | <.001 |
| | | | | | | 8-10k | 18-20k | <.001 | <.001 |
| | | | | | | | 28-30k | <.001 | <.001 |
| | | | | | | 18-20k | 28-30k | <.001 | 0.064 |

**Figure 5E** *Fraction of trials near target. 2-Way RM ANOVA, Post hoc: Holm*
Group: $F(1,20) = 5.607$, $p = 0.028$. Block: $F(3,60) = 182.793$, $p<0.001$.
Group x Block: $F(3,60) = 6.889$, $p<0.001$

| Between Groups | | | | | | Within Group | | | |
|---|---|---|---|---|---|---|---|---|---|
| | | 0-2k | 8-10k | 18-20k | 28-30k | | vs. | Control | $ATN^{DMS}$ |
| Control | $ATN^{DMS}$ | 0.729 | 0.003 | 0.056 | 0.387 | 0-2k | 8-10k | <0.001 | <0.001 |
| All | | | | | | | 18-20k | <0.001 | <0.001 |
| Control | $ATN^{DMS}$ | 0.028 | | | | | 28-30k | <0.001 | <0.001 |
| | | | | | | 8-10k | 18-20k | <0.001 | 0.026 |
| | | | | | | | 28-30k | <0.001 | 0.026 |
| | | | | | | 18-20k | 28-30k | 0.003 | 0.304 |

**Figure 5G** Number of trials needed to reach learning criterion. 1-Way ANOVA
Group: $F(1,19) = 2.200$, $p = 0.154$

**Figure 5H** (Left)**:** ΔIPI after reward (high reward context)*.2-Way RM ANOVA, Post hoc: Holm*
Group: $F(1,20) = 1.629$, $p = 0.217$. Block: $F(3,60) = 13.097$, $p<0.001$
Group x Block: $F(3,60) = 2.815$, $p = 0.047$

| Between Groups | | | | | | Within Group | | | |
|---|---|---|---|---|---|---|---|---|---|
| | vs. | 0-2k | 8-10k | 18-20k | 28-30k | | vs. | Control | ATN$^{DMS}$ |
| Control | ATN$^{DMS}$ | 0.391 | 0.026 | 0.324 | 0.385 | 0-2k | 8-10k | 0.436 | 0.148 |
| | | | | | | | 18-20k | 0.181 | 0.036 |
| | | | | | | | 28-30k | 0.114 | 0.021 |
| | | | | | | 8-10k | 18-20k | 0.006 | 0.148 |
| | | | | | | | 28-30k | 0.006 | 0.040 |
| | | | | | | 18-20k | 28-30k | 0.198 | 0.148 |

**Figure 5H** (Right)**:** ΔIPI after no reward (high reward context)*. 2-Way RM ANOVA.*
Group: $F(1,20) = 0.324$, $p = 0.576$. Block: $F(3,60) = 13.100$, $p<0.001$
Group x Block: $F(3,60) = 0.571$, $p = 0.636$

**Figure 5I** *Kinematic correlation after reward. Linear mixed-effects model (LME), random intercept per rat.*
Group: $F(1,19.666) = 0.077$, $p = 0.785$. Block: $F(3,57.0186) = 19.976$, $p<0.001$.
Group x Block: $F(3,57.019) = 1.142$, $p = 0.340$

**Supplementary Table 5. Statistical Results for Figure 6**

**Figure 6C** *Mean IPI over the course of training. 2-Way RM ANOVA*
Group: F(1,21) = 2.640, p = 0.119. Block: F(3,63) = 63.082, p<0.001
Group x Block: F(3,63) = 1.537, p = 0.2136

**Figure 6D** *IPI variability over the course of training. 2-Way RM ANOVA, Post hoc: Holm*
Group: F(1,21) = 4.676, p = 0.042. Block: F(3,63) = 127.013, p<0.001.
Group x Block: F(3,60) = 3.485, p = 0.021

| Between Groups | | | | | | Within Group | | | |
|---|---|---|---|---|---|---|---|---|---|
| | vs. | 0-2k | 8-10k | 18-20k | 28-30k | | vs. | Cont. | $^{STN}ATN^{DMS}$ |
| Controls | $^{STN}ATN^{DMS}$ | 0.150 | 0.012 | 0.287 | 0.927 | 0-2k | 8-10k | <.001 | 0.001 |
| | | All | | | | | 18-20k | <.001 | <.001 |
| Controls | $^{STN}ATN^{DMS}$ | 0.042 | | | | | 28-30k | <.001 | <.001 |
| | | | | | | 8-10k | 18-20k | <.001 | 0.067 |
| | | | | | | | 28-30k | <.001 | 0.043 |
| | | | | | | 18-20k | 28-30k | <.001 | 0.067 |

**Figure 6E** *Fraction of trials near target over the course of training. 2-Way RM ANOVA, Post hoc: Holm*
Group: F(1,21) = 2.172, p = 0.015. Block: F(3,63) = 115.896, p<0.001
Group x Block: F(3,63) = 3.084, p = 0.034

| Between Groups | | | | | | Within Group | | | |
|---|---|---|---|---|---|---|---|---|---|
| | vs. | 0-2k | 8-10k | 18-20k | 28-30k | | vs. | Cont. | $^{STN}ATN^{DMS}$ |
| Controls | $^{STN}ATN^{DMS}$ | 0.495 | 0.045 | 0.207 | 0.889 | 0-2k | 8-10k | <.001 | <.001 |
| | | | | | | | 18-20k | <.001 | <.001 |
| | | | | | | | 28-30k | <.001 | <.001 |
| | | | | | | 8-10k | 18-20k | <.001 | 0.126 |
| | | | | | | | 28-30k | <.001 | 0.095 |
| | | | | | | 18-20k | 28-30k | 0.003 | 0.335 |

**Figure 6G** Number of trials needed to reach learning criterion. 1-Way ANOVA
Group: F(1,19) = 5.175, p = 0.034

**Figure 6H** (Left): ΔIPI after reward (high reward context).*2-Way RM ANOVA.*
Group: F(1,21) = 1.436, p = 0.244. Block: F(3,63) = 10.368, p<0.001
Group x Block: F(3,63) = 1.411, p = 0.248

**Figure 6H** (Right): ΔIPI after no reward (high reward context).*2-Way RM ANOVA.*
Group: F(1,21) = 0.376, p = 0.546. Block: F(3,63) = 7.991, p<0.001
Group x Block: F(3,63) = 0.976, p = 0.410

**Figure 6I** *Kinematic correlation after reward. Linear mixed-effects model (LME), random intercept per rat. Post hoc: Estimated marginal means from the LME, with holm correction.*
Group: F(1,21.15) = 0.256, p = 0.618. Block: F(3,61.34) = 28.605, p<0.001.
Group x Block: F(3,61.34) = 0.967, p = 0.414

**Supplementary Table 6. Statistical Results for Extended Data Figure 2.**

**Extended Data Figure 2A** *Mean IPI over the course of training. 2-Way RM ANOVA*
Group: F(4,17) = 1.00, p = 0.434. Block: F(3,51) = 67.965, p<0.001.
Group x Block: F(12,51) = 1.000, p = 0.462

**Extended Data Figure 2B** *IPI variability over the course of training. 2-Way RM ANOVA, Post hoc: Holm*
Group: F(4,17) = 0.169, p = 0.951. Block: F(3,51) = 165.904, p<0.001.
Group x Block: F(6,87) = 2.324, p = 0.018

| Between Groups | | | | | | Within Group | | | | | | |
|---|---|---|---|---|---|---|---|---|---|---|---|---|
| **Shams** | vs. | 0-2k | 8-10k | 18-20k | 28-30k | | vs | $STN^{DLS}$ | $STN^{ATN}$ | $ATN^{DMS}$ | $STN^{Thal}$ | $STN^{DMS}$ |
| $STN^{DLS}$ | $STN^{ATN}$ | 0.398 | 1 | 1 | 1 | 0-2k | 8-10k | 0.12 | 0.017 | 0.053 | 0.009 | 0.025 |
| | $ATN^{DMS}$ | 1 | 1 | 1 | 1 | | 18-20k | 0.012 | 0.023 | 0.01 | 0.005 | 0.052 |
| | $STN^{Thal}$ | 1 | 1 | 1 | 1 | | 28-30k | 0.012 | 0.02 | 0.005 | 0.003 | 0.052 |
| | $STN^{DMS}$ | 1 | 1 | 1 | 0.708 | 8-10k | 18-20k | 0.104 | 0.127 | 0.015 | 0.028 | 0.405 |
| $STN^{ATN}$ | $ATN^{DMS}$ | 1 | 1 | 1 | 1 | | 28-30k | 0.12 | 0.084 | 0.015 | 0.03 | 0.405 |
| | $STN^{Thal}$ | 1 | 1 | 1 | 1 | 18-20k | 28-30k | 0.137 | 0.127 | 0.053 | 0.068 | 0.405 |
| | $STN^{DMS}$ | 1 | 1 | 1 | 0.237 | | | | | | | |
| $ATN^{DMS}$ | $STN^{Thal}$ | 1 | 1 | 1 | 1 | | | | | | | |
| | $STN^{DMS}$ | 1 | 1 | 1 | 0.081 | | | | | | | |
| $STN^{Thal}$ | $STN^{DMS}$ | 1 | 1 | 1 | 0.371 | | | | | | | |

**Extended Data Figure 2C** Fraction of trials near target. *2-Way RM ANOVA*
Group: F(4,17) = 0.126, p = 0.971. Block: F(3,51) = 146.741, p<0.001.
Group x Block: F(6,87) = 0.833, p = 0.617

**Extended Data Figure 2E** Number of trials needed to reach learning criterion. 1-Way ANOVA
Group: F(4,15) = 0.240, p = 0.911

**Extended Data Figure 2F** *Mean ITI over the course of training. 2-Way RM ANOVA*
Group: F(4,17) = 0.497, p = 0.738. Block: F(3,51) = 55.013, p<0.001
Group x Block: F(6,87) = 1.113, p = 0.370

**Extended Data Figure 2G** *ITI Variability over the course of training. 2-Way RM ANOVA, Post hoc: hom*
Group: F(4,17) = 0.004, p = 0.100. Block: F(3,51) = 107.065, p<0.001.
Group x Block: F(6,87) = 2.086, p = 0.035

| Between Groups | | | | | | Within Group | | | | | | |
|---|---|---|---|---|---|---|---|---|---|---|---|---|
| **Shams** | vs. | 0-2k | 8-10k | 18-20k | 28-30k | | vs | $STN^{DLS}$ | $STN^{ATN}$ | $ATN^{DMS}$ | $STN^{Thal}$ | $STN^{DMS}$ |
| $STN^{DLS}$ | $STN^{ATN}$ | 1.000 | 1.000 | 1.000 | 1.000 | 0-2k | 8-10k | 0.126 | 0.163 | 0.236 | 0.164 | 0.391 |
| | $ATN^{DMS}$ | 1.000 | 1.000 | 1.000 | 1.000 | | 18-20k | 0.004 | 0.001 | 0.027 | 0.009 | 0.185 |
| | $STN^{Thal}$ | 1.000 | 1.000 | 1.000 | 1.000 | | 28-30k | 0.004 | <.001 | 0.039 | 0.010 | 0.185 |
| | $STN^{DMS}$ | 1.000 | 1.000 | 1.000 | 1.000 | 8-10k | 18-20k | 0.102 | 0.163 | 0.089 | 0.164 | 0.185 |
| $STN^{ATN}$ | $ATN^{DMS}$ | 1.000 | 1.000 | 1.000 | 1.000 | | 28-30k | 0.126 | 0.163 | 0.084 | 0.172 | 0.064 |
| | $STN^{Thal}$ | 1.000 | 1.000 | 1.000 | 1.000 | 18-20k | 28-30k | 0.198 | 0.163 | 0.294 | 0.578 | 0.607 |
| | $STN^{DMS}$ | 0.319 | 1.000 | 1.000 | 0.369 | | | | | | | |
| $ATN^{DMS}$ | $STN^{Thal}$ | 1.000 | 1.000 | 1.000 | 1.000 | | | | | | | |
| | $STN^{DMS}$ | 0.946 | 1.000 | 1.000 | 1.000 | | | | | | | |
| $STN^{Thal}$ | $STN^{DMS}$ | 0.423 | 1.000 | 1.000 | 1.000 | | | | | | | |

**Supplementary Table 7. Statistical Results for Extended Data Figure 3.**

**Extended Data Figure 3A** number of trials per session *over the course of training. 2-Way RM ANOVA*
Group: F(2,29) = 0.995, p = 0.382. Block: F(3,87) = 5.464, p = 0.026
Group x Block: F(6,87) = 0.729, p = 0.491

**Extended Data Figure 3B** *Mean reward per trial per session over the course of training. 2-Way RM ANOVA*
Group: F(2,29) = 1.623, p = 0.215. Block: F(3,87) = 0.454, p = 0.506.
Group x Block: F(6,87) = 1.404, p = 0.262

**Extended Data Figure 3C** *Fraction of 1 tap trials over the course of training. 2-Way RM ANOVA*
Group: F(2,29) = 2.091, p = 0.142. Block: F(3,87) = 40.218, p<0.001
Group x Block: F(6,87) = 0.768, p = 0.597

**Extended Data Figure 3D** *Fraction of 2 tap trials over the course of training. 2-Way RM ANOVA Post hoc: Holm*
Group: F(2,29) = 1.615, p = 0.216. Block: F(3,87) = 17.646, p<0.001
Group x Block: F(6,87) = 2.392, p = 0.034

| Between Groups | | | | | | Within Group | | | | |
|---|---|---|---|---|---|---|---|---|---|---|
| | vs. | 0-2k | 8-10k | 18-20k | 28-30k | | vs. | Cont. | $STN^{DLS}$ | $STN^{ATN}$ |
| Controls | $STN^{DLS}$ | 1.000 | 0.863 | 0.387 | 0.021 | 0-2k | 8-10k | 0.014 | 0.127 | 0.065 |
| | $STN^{ATN}$ | 1.000 | 0.218 | 0.916 | 0.276 | | 18-20k | <0.001 | 0.541 | 0.065 |
| $STN^{DLS}$ | $STN^{ATN}$ | 1.000 | 0.241 | 0.916 | 0.276 | | 28-30k | <0.001 | 1.000 | 0.038 |
| | | | | | | 8-10k | 18-20k | <0.001 | 1.000 | 1.000 |
| | | | | | | | 28-30k | <0.001 | 1.000 | 1.000 |
| | | | | | | 18-20k | 28-30k | 0.261 | 1.000 | 1.000 |

**Extended Data Figure 3E** *Fraction of 3 tap trials over the course of training. 2-Way RM ANOVA*
Group: F(2,29) = 0.086, p = 0.433. Block: F(3,87) = 4.818, p = 0.004
Group x Block: F(6,87) = 1.331, p = 0.252

**Extended Data Figure 3F** *Average Velocity. 1-Way ANOVA,*
Group: F(2,29) = 0.145, p = 0.866.

**Extended Data Figure 3G** *Average Acceleration. 1-Way ANOVA,*
Group: F(2,29) = 0.123, p = 0.885

**Extended Data Figure 3H** *Mean ITI over the course of training. 2-Way RM ANOVA*
Group: F(2,29) = 0.567, p = 0.573. Block: F(3,87) = 52.319, p<0.001
Group x Block: F(6,87) = 1.637, p = 0.147

**Extended Data Figure 3I** *ITI variability over the course of training. 2-Way RM ANOVA, Post hoc: Holm*
Group: F(2,29) = 3.738, p = 0.036. Block: F(3,87) =136.577, p<0.001
Group x Block: F(6,87) = 3.439, p = 0.004

| Between Groups | | | | | | Within Group | | | | |
|---|---|---|---|---|---|---|---|---|---|---|
| | vs. | 0-2k | 8-10k | 18-20k | 28-30k | | vs. | Controls | $STN^{DLS}$ | $STN^{ATN}$ |
| Controls | $STN^{DLS}$ | 0.568 | 0.686 | 0.100 | 0.209 | 0-2k | 8-10k | <0.001 | 0.014 | <0.001 |
| | $STN^{ATN}$ | 0.279 | 0.017 | 0.328 | 0.888 | | 18-20k | <0.001 | 0.005 | <0.001 |
| $STN^{DLS}$ | $STN^{ATN}$ | 0.568 | 0.041 | 0.026 | 0.246 | | 28-30k | <0.001 | <0.001 | <0.001 |
| | vs. | All | | | | 8-10k | 18-20k | <0.001 | 0.078 | 0.271 |
| Controls | $STN^{DLS}$ | 0.480 | | | | | 28-30k | <0.001 | <0.001 | 0.271 |
| | $STN^{ATN}$ | 0.091 | | | | 18-20k | 28-30k | 0.008 | 0.015 | 0.994 |
| $STN^{DLS}$ | $STN^{ATN}$ | 0.041 | | | | | | | | |

**Supplementary Table 8. Statistical Results for Extended Data Figure 5.**

| |
|---|
| **Extended Data Figure 5C** *Mean IPI over the course of training. 2-Way RM ANOVA*<br>Group: F(1,16) = 3.356, p = 0.086. Block: F(3,48) = 44.026, p<0.001.<br>Group x Block: F(3,48) = 1.449, p = 0.240 |
| **Extended Data Figure 5D** *IPI variability over the course of training. 2-Way RM ANOVA*<br>Group: F(1,16) = 0.436, p = 0.518. Block: F(3,48) = 90.422, p<0.001.<br>Group x Block: F(3,48) = 2.264, p = 0.093 |
| **Extended Data Figure 5E** *Fraction of trials near target. 2-Way RM ANOVA*<br>Group: F(1,16) = 0.198, p = 0.662. Block: F(3,48) = 90.863, p<0.001.<br>Group x Block: F(3,48) = 0.358, p = 0.784 |
| **Extended Data Figure 5G** Number of trials needed to reach learning criterion. 1-Way ANOVA<br>Group: F(1,12) = 0.816, p = 0.384 |
| **Extended Data Figure 5H** *Mean ITI over the course of training. 2-Way RM ANOVA*<br>Group: F(1,16) = 0.430, p = 0.521. Block: F(3,48) = 51.771, p<0.001.<br>Group x Block: F(3,48) = 1.457, p = 0.238 |
| **Extended Data Figure 5I** *ITI variability over the course of training. 2-Way RM ANOVA*<br>Group: F(1,16) = 1.047, p = 0.321. Block: F(3,48) = 63.840, p<0.001.<br>Group x Block: F(3,48) = 0.737, p = 0.535 |

**Supplementary Table 9. Statistical Results for Extended Data Figure 7.**

| |
|---|
| **Extended Data Figure 7C** *Mean IPI over the course of training. 2-Way RM ANOVA*<br>Group: F(2,21) = 0.394, p = 0.679. Block: F(3,63) = 58.933, p<0.001.<br>Group x Block: F(6,63) = 0.785, p = 0.585 |
| **Extended Data Figure 7D** *IPI variability over the course of training. 2-Way RM ANOVA*<br>Group: F(2,21) = 0.588, p = 0.564. Block: F(3,63) = 126.83, p<0.001.<br>Group x Block: F(6,63) = 1.592, p = 0.164 |
| **Extended Data Figure 7E** *Fraction of trials near target. 2-Way RM ANOVA*<br>Group: F(2,21) = 0.653, p = 0.530. Block: F(3,63) = 124.568, p<0.001.<br>Group x Block: F(6,63) = 1.317, p = 0.263 |
| **Extended Data Figure 7G** Number of trials needed to reach learning criterion. 1-Way ANOVA<br>Group: F(2,19) = 0.227, p = 0.7992 |
| **Extended Data Figure 7H** *Mean ITI over the course of training. 2-Way RM ANOVA*<br>Group: F(2,21) = 1.628, p = 0.220. Block: F(3,63) = 45.976, p<0.001<br>Group x Block: F(6,63) = 1.033, p = 0.412 |
| **Extended Data Figure 7I** *ITI variability over the course of training. 2-Way RM ANOVA*<br>Group: F(2,21) = 0.373, p = 0.693. Block: F(3,63) = 131.907, p<0.001.<br>Group x Block: F(6,63) = 1.651, p = 0.148 |

**Supplementary Table 10. Statistical Results for Extended Data Figure 8**

**Extended Data Figure 8A** (Left)**:** ΔIPI after reward (across all reward contexts). *2-Way RM ANOVA, Post hoc: Holm*
Group: F(2,29) = 6.786, p = 0.004. Block: F(3,87) = 19.023, p<0.001
Group x Block: F(6,87) = 3.790, p = 0.002

| Between Groups | | | | | | Within Group | | | | |
|---|---|---|---|---|---|---|---|---|---|---|
| | vs. | 0-2k | 8-10k | 18-20k | 28-30k | | vs. | Cont. | $STN^{DLS}$ | $STN^{ATN}$ |
| Controls | $STN^{DLS}$ | 1.000 | 0.473 | 0.024 | 0.002 | 0-2k | 8-10k | 0.714 | 1.000 | 0.009 |
| | $STN^{ATN}$ | 1.000 | 0.013 | 0.467 | 0.569 | | 18-20k | 0.015 | 1.000 | 0.009 |
| $STN^{DLS}$ | $STN^{ATN}$ | 1.000 | 0.005 | 0.011 | 0.002 | | 28-30k | 0.010 | 1.000 | 0.004 |
| | | All | | | | 8-10k | 18-20k | <.001 | 1.000 | 0.683 |
| Controls | $STN^{DLS}$ | 0.03 | | | | | 28-30k | <.001 | 1.000 | 0.206 |
| | $STN^{ATN}$ | 0.205 | | | | 18-20k | 28-30k | 0.033 | 1.000 | 0.585 |
| $STN^{DLS}$ | $STN^{ATN}$ | 0.004 | | | | | | | | |

**Extended Data Figure 8A** (Right): ΔIPI after no reward (across all reward contexts). *2-Way RM ANOVA*
Group: F(2,29) = 1.325, p = 0.281. Block: F(3,87) = 15.030, p<0.001
Group x Block: F(6,87) = 1.484, p = 0.019

**Extended Data Figure 8B** (Left): ΔIPI after reward (low reward context). *2-Way RM ANOVA, Post hoc: Holm*
Group: F(2,29) = 6.635, p = 0.004. Block: F(3,87) = 36.140, p<0.001.
Group x Block: F(6,87) = 2.433, p = 0.032

| Between Groups | | | | | | Within Group | | | | |
|---|---|---|---|---|---|---|---|---|---|---|
| | vs. | 0-2k | 8-10k | 18-20k | 28-30k | | vs. | Cont. | $STN^{DLS}$ | $STN^{ATN}$ |
| Controls | $STN^{DLS}$ | 1.000 | 0.337 | 0.027 | 0.003 | 0-2k | 8-10k | 0.204 | 0.374 | 0.015 |
| | $STN^{ATN}$ | 1.000 | 0.088 | 0.431 | 0.420 | | 18-20k | 0.003 | 0.247 | 0.004 |
| $STN^{DLS}$ | $STN^{ATN}$ | 1.000 | 0.022 | 0.011 | 0.001 | | 28-30k | 0.002 | 0.136 | 0.001 |
| | vs. | All | | | | 8-10k | 18-20k | 0.001 | 0.247 | 0.470 |
| Controls | $STN^{DLS}$ | 0.005 | | | | | 28-30k | <0.001 | 0.232 | 0.040 |
| | $STN^{ATN}$ | 0.269 | | | | 18-20k | 28-30k | 0.078 | 0.280 | 0.470 |
| $STN^{DLS}$ | $STN^{ATN}$ | 0.025 | | | | | | | | |

**Extended Data Figure 8B** (Right): ΔIPI after no reward (low reward context). *2-Way RM ANOVA*
Group: F(2,29) = 1.683, p = 0.204. Block: F(3,87) = 15.6768, p<0.001.
Group x Block: F(6,87) = 1.624, p = 0.150

**Extended Data Figure 8C** (Left): IPI variance (3-trial window) after reward. *2-Way RM ANOVA, Post hoc: Holm.* Group: F(2,29) = 3.73, p = 0.036. Block: F(3,87) = 31.800, p<0.001.
Group x Block: F(6,87) = 2.468, p = 0.030

| Between Groups | | | | | | Within Group | | | | |
|---|---|---|---|---|---|---|---|---|---|---|
| | vs. | 0-2k | 8-10k | 18-20k | 28-30k | | vs. | Cont. | $STN^{DLS}$ | $STN^{ATN}$ |
| Controls | $STN^{DLS}$ | 1 | 0.494 | 0.066 | 0.003 | 0-2k | 8-10k | 0.145 | 0.570 | 0.010 |
| | $STN^{ATN}$ | 1 | 0.054 | 0.706 | 0.852 | | 18-20k | 0.003 | 0.285 | 0.007 |
| $STN^{DLS}$ | $STN^{ATN}$ | 1 | 0.025 | 0.066 | 0.003 | | 28-30k | 0.002 | 0.186 | 0.007 |
| | vs. | All | | | | 8-10k | 18-20k | 0.002 | 0.570 | 0.791 |
| Controls | $STN^{DLS}$ | 0.097 | | | | | 28-30k | 0.001 | 0.570 | 0.180 |
| | $STN^{ATN}$ | 0.453 | | | | 18-20k | 28-30k | 0.047 | 0.570 | 0.424 |
| $STN^{DLS}$ | $STN^{ATN}$ | 0.043 | | | | | | | | |

**Extended Data Figure 8C** (Right): IPI variance (3-trial window) after no reward. *2-Way RM ANOVA*
Group: F(2,29) = 1.269, p = 0.296. Block: F(3,87) = 30.347, p<0.001.
Group x Block: F(6,87) = 1.651, p = 0.143

**Extended Data Figure 8D** (Left): IPI variance (4-trial window) after reward. *2-Way RM ANOVA, Post hoc: Holm*
Group: F(2,29) = 3.215, p = 0.055. Block: F(3,87) = 29.546, p<0.001.
Group x Block: F(6,87) = 2.397, p = 0.034

| Between Groups | | | | | | Within Group | | | | |
|---|---|---|---|---|---|---|---|---|---|---|
| | vs. | 0-2k | 8-10k | 18-20k | 28-30k | | vs. | Cont. | $STN^{DLS}$ | $STN^{ATN}$ |
| Controls | $STN^{DLS}$ | 1.000 | 0.534 | 0.105 | 0.003 | 0-2k | 8-10k | 0.150 | 0.594 | 0.012 |
| | $STN^{ATN}$ | 1.000 | 0.061 | 0.774 | 0.949 | | 18-20k | 0.004 | 0.319 | 0.010 |
| $STN^{DLS}$ | $STN^{ATN}$ | 1.000 | 0.033 | 0.105 | 0.006 | | 28-30k | 0.003 | 0.189 | 0.010 |
| | | | | | | 8-10k | 18-20k | 0.003 | 0.594 | 0.933 |
| | | | | | | | 28-30k | <0.001 | 0.594 | 0.526 |
| | | | | | | 18-20k | 28-30k | 0.043 | 0.594 | 0.526 |

**Extended Data Figure 8D** (Right): IPI variance (4-trial window) after no reward. *2-Way RM ANOVA*
Group: F(2,29) = 1.563, p = 0.227. Block: F(3,87) = 30.078, p<0.001.
Group x Block: F(6,87) = 1.714, p = 0.127

**Extended Data Figure 8E** (Left): ∆IPI after reward. *2-Way RM ANOVA*
Group: F(1,20) = 1.182, p = 0.290. Block: F(3,60) = 23.487, p<0.001.
Group x Block: F(3,60) = 2.491, p = 0.069

**Figure 8E** (Right): ∆IPI after no reward. *2-Way RM ANOVA*
Group: F(1,20) = 1.128, p = 0.300. Block: F(3,60) = 14.885, p<0.001.
Group x Block: F(3,60) = 0.393, p = 0.758

**Extended Data Figure 8F** (Left): ∆IPI after reward (low reward context). *2-Way RM ANOVA.*
Group: F(1,20) = 0.848, p = 0.368. Block: F(3,60) = 29.512, p<0.001
Group x Block: F(3,60) = 1.733, p = 0.170

**Extended Data Figure 8F** (Right): ∆IPI after no reward (low reward context). *2-Way RM ANOVA.*
Group: F(1,20) = 1.661, p = 0.212. Block: F(3,60) = 13.180, p<0.001
Group x Block: F(3,60) = 0.328, p = 0.805

**Extended Data Figure 8G** (Left): IPI variance (2 trial window) after reward. *2-Way RM ANOVA. Post hoc: Holm*
Group: F(1,20) = 0.624, p = 0.439. Block: F(3,60) = 34.189, p<0.001
Group x Block: F(3,60) = 3.103, p = 0.033

| Between Groups | | | | | | Within Group | | | |
|---|---|---|---|---|---|---|---|---|---|
| | vs. | 0-2k | 8-10k | 18-20k | 28-30k | | vs. | Cont. | $ATN^{DMS}$ |
| Control | $ATN^{DMS}$ | 0.337 | 0.036 | 0.308 | 0.350 | 0-2k | 8-10k | 0.224 | 0.039 |
| | | | | | | | 18-20k | 0.002 | 0.008 |
| | | | | | | | 28-30k | 0.001 | 0.008 |
| | | | | | | 8-10k | 18-20k | <.001 | 0.044 |
| | | | | | | | 28-30k | <.001 | 0.003 |
| | | | | | | 18-20k | 28-30k | 0.014 | 0.155 |

**Extended Data Figure 8G** (Right): IPI variance (2 trial window) after no reward. *2-Way RM ANOVA.*
Group: F(1,20) = 0.772, p = 0.390. Block: F(3,60) = 26.771, p<0.001
Group x Block: F(3,60) = 0.678, p = 0.569

**Extended Data Figure 8H** (Left): IPI variance (5 trial window) after reward. *2-Way RM ANOVA.*
Group: F(1,20) = 0.585, p = 0.453. Block: F(3,60) = 31.190, p<0.001
Group x Block: F(3,60) = 2.258, p = 0.091

**Extended Data Figure 8H** (Right): IPI variance (5 trial window) after no reward. *2-Way RM ANOVA.*
Group: F(1,20) =0.942, p = 0.343. Block: F(3,60) = 29.033, p<0.001
Group x Block: F(3,60) = 1.199, p = 0.318

**Extended Data Figure 8I** (Left): ∆IPI after reward (across all reward contexts). 2-Way RM ANOVA
Group: F(1,21) = 2.738, p = 0.113. Block: F(3,63) = 21.747, p<0.001.
Group x Block: F(3,63) = 1.967, p = 0.128

**Extended Data Figure 8I** (Right): ∆IPI after no reward (across all reward contexts). 2-Way RM ANOVA
Group: F(1,21) = 0.732 p = 0.402. Block: F(3,63) = 11.553, p<0.001.
Group x Block: F(3,63) = 1.336, p =0.271

**Extended Data Figure 8J** (Left): ∆IPI after reward (low reward context). *2-Way RM ANOVA. Post hoc: Holm*
Group: F(1,21) = 1.971, p = 0.175. Block: F(3,63) = 37.046, p<0.001
Group x Block: F(3,63) = 2.980, p = 0.040

| Between Groups | | | | | | Within Group | | | |
|---|---|---|---|---|---|---|---|---|---|
| | vs. | 0-2k | 8-10k | 18-20k | 28-30k | | vs. | Cont. | $^{STN}ATN^{DMS}$ |
| Controls | $^{STN}ATN^{DMS}$ | 0.585 | 0.007 | 0.226 | 0.456 | 0-2k | 8-10k | 0.204 | 0.004 |
| | | | | | | | 18-20k | 0.003 | 0.004 |
| | | | | | | | 28-30k | 0.002 | <.001 |
| | | | | | | 8-10k | 18-20k | 0.001 | 0.197 |
| | | | | | | | 28-30k | <.001 | 0.052 |
| | | | | | | 18-20k | 28-30k | 0.078 | 0.197 |

**Extended Data Figure 8J** (Right): ∆IPI after no reward (low reward context).*2-Way RM ANOVA.*
Group: F(1,21) = 0.415, p = 0.527. Block: F(3,63) = 11.98, p<0.001
Group x Block: F(3,63) = 2.023, p = 0.120

**Extended Data Figure 8K** (Left): IPI variance (2 trial window) after reward. *2-Way RM ANOVA.*
Group: F(1,21) = 2.627, p = 0.120. Block: F(3,63) = 35.615, p<0.001
Group x Block: F(3,63) = 2.608, p = 0.059

**Extended Data Figure 8K** (Right): IPI variance (2 trial window) after no reward. *2-Way RM ANOVA.*
Group: F(1,21) = 1.195, p = 0.287. Block: F(3,63) = 23.197, p<0.001
Group x Block: F(3,63) = 1.689, p = 0.178

**Extended Data Figure 8L** (Left): IPI variance (5 trial window) after reward. *2-Way RM ANOVA.*
Group: F(1,21) = 2.091, p = 0.163. Block: F(3,63) = 28.167, p<0.001
Group x Block: F(3,63) = 2.228, p = 0.094

**Extended Data Figure 8L** (Right): IPI variance (5 trial window) after no reward. *2-Way RM ANOVA.*
Group: F(1,21) = 1.371, p = 0.255. Block: F(3,63) = 26.043, p<0.001
Group x Block: F(3,63) = 2.202, p = 0.096

**Supplementary Table 11. Statistical Results for Extended Data Figure 9**

**Extended Data Figure 9A** *Kinematic correlation (All Trials). Linear mixed-effects model (LME), random intercept per rat. Post hoc: Estimated marginal means from the LME, with holm correction.*
Group: F(2,29.01) = 5.577, p = 0.009. Block: F(3,83.42) = 26.920, p<0.001.
Group x Block: F(6,83.39) = 1.962, p = 0.0803

| | vs. | All |
|---|---|---|
| Controls | STN$^{DLS}$ | 0.04 |
| | STN$^{ATN}$ | 0.323 |
| STN$^{DLS}$ | STN$^{ATN}$ | <0.001 |

**Extended Data Figure 9B** *Kinematic correlation (All Trials). LME, random intercept per rat*
Group: F(1,20.09) = 0.137, p = 0.715. Block: F(3,58.28) = 22.44, p<0.001.
Group x Block: F(3,58.28) = 0.77, p = 0.514

**Extended Data Figure 9C** *Kinematic correlation after no reward. LME, random intercept per rat*
Group: F(1,19.97) = 0.074, p = 0.788. Block: F(3,58.14) = 27.48, p<0.001.
Group x Block: F(3,58.14) = 0.248, p = 0.862

**Extended Data Figure 9D** *Number of Modes. .LME, random intercept per rat*
Group: F(1,20) = 0.290, p = 0.596. Block: F(3,60) = 1.70, p = 0.177.
Group x Block: F(3,60) = 0.110, p = 0.954

**Extended Data Figure 9E** *Within block main mode. LME, random intercept per rat*
Group: F(1,19.94) = 0.505, p = 0.485. Block: F(3,58.13) = 2.308, p = 0.086.
Group x Block: F(3,58.13) = 0.880, p = 0.457

**Extended Data Figure 9F** *Final block main mode. LME, random intercept per rat*
Group: F(1,19.93) = 1.795, p = 0.195. Block: F(3,58.06) = 34.12, p<0.001.
Group x Block: F(3,58.06) = 0.146, p = 0.9321

**Extended Data Figure 9G** *Within mode correlation. LME, random intercept per rat*
Group: F(1,20.27) = 0.059, p = 0.811. Block: F(3,58.59) = 33.395, p<0.001.
Group x Block: F(3,58.59) = 1.267, p = 0.294

**Extended Data Figure 9H** *Kinematic correlation (All Trials). LME, random intercept per rat*
Group: F(1,21.1) = 0.372, p = 0.548. Block: F(3,61.26) = 37.87, p<0.001.
Group x Block: F(3,61.26) = 1.162, p = 0.332

**Extended Data Figure 9I** *Kinematic correlation after no reward. LME, random intercept per rat*
Group: F(1,21.07) = 0.378, p = 0.545. Block: F(3,61.23) = 44.916, p<0.001.
Group x Block: F(3,61.23) = 0.930, p = 0.432

**Extended Data Figure 9J** *Number of Modes. LME, random intercept per rat*
Group: F(1,21) = 2.495, p = 0.129. Block: F(3,63) = 1.018, p = 0.391.
Group x Block: F(3,63) = 2.027, p = 0.119

**Extended Data Figure 9K** *Within block main mode. LME, random intercept per rat*
Group: F(1,21.1) = 1.848, p = 0.188. Block: F(3,61.29) = 1.813, p = 0.154.
Group x Block: F(3,61.29) = 0.524, p = 0.667

**Extended Data Figure 9L** *Final block main mode. LME, random intercept per rat*
Group: F(1,20.98) = 0.662, p = 0.425. Block: F(3,61.13) = 28.703, p<0.001.
Group x Block: F(3,61.13) = 0.519, p = 0.671

**Extended Data Figure 9M** *Within mode correlation. LME, random intercept per rat*
Group: F(1,21.11) = 1.724, p = 0.203. Block: F(3,61.35) = 29.285, p<0.001.
Group x Block: F(3,61.35) = 1.048, p = 0.378
